# Enhancer viruses and a transgenic platform for combinatorial cell subclass-specific labeling

**DOI:** 10.1101/525014

**Authors:** Lucas T. Graybuck, Tanya L. Daigle, Adriana E. Sedeño-Cortés, Miranda Walker, Brian Kalmbach, Garreck H. Lenz, Thuc Nghi Nguyen, Emma Garren, Tae Kyung Kim, La’ Akea Siverts, Jacqueline L. Bendrick, Thomas Zhou, Marty Mortrud, Shenqin Yao, Ali H. Cetin, Rachael Larsen, Luke Esposito, Bryan Gore, Eric Szelenyi, Elyse Morin, John K. Mich, Nick Dee, Jeff Goldy, Kimberly Smith, Zizhen Yao, Viviana Gradinaru, Susan M. Sunkin, Ed Lein, Boaz P. Levi, Jonathan T. Ting, Hongkui Zeng, Bosiljka Tasic

## Abstract

The rapid pace of cell type identification by new single-cell analysis methods has not been met with efficient experimental access to the newly discovered types. To enable flexible and efficient access to specific neural populations in the mouse cortex, we collected chromatin accessibility data from individual cells and clustered the single-cell data to identify enhancers specific for cell classes and subclasses. When cloned into adeno-associated viruses (AAVs) and delivered to the brain by retro-orbital injections, these enhancers drive transgene expression in specific cell subclasses in the cortex. We characterize several enhancer viruses in detail to show that they result in labeling of different projection neuron subclasses in mouse cortex, and that one of them can be used to label the homologous projection neuron subclass in human cortical slices. To enable the combinatorial labeling of more than one cell type by enhancer viruses, we developed a three-color Cre-, Flp- and Nigri-recombinase dependent reporter mouse line, *Ai213*. The delivery of three enhancer viruses driving these recombinases via a single retroorbital injection into a single *Ai213* transgenic mouse results in labeling of three different neuronal classes/subclasses in the same brain tissue. This approach combines unprecedented flexibility with specificity for investigation of cell types in the mouse brain and beyond.

## Introduction

To understand brain function, we need to define brain cell types and build genetic tools to selectively label and perturb them for further study (Tasic et al., 2018; Zeng and Sanes, 2017). In mice, transgenic recombinase driver and reporter lines have been used to great effect to label cell populations that share marker gene expression (Daigle et al., 2018; Gong et al., 2007; Taniguchi et al., 2011). However, the creation, maintenance, and sharing of transgenic mouse lines is costly. Establishment of lines that label more than one cell type or class requires laborious crosses, which yield a low frequency of experimental animals with three or four transgenes due to the laws of Mendelian genetics.

Recent advances in single-cell profiling, such as single-cell RNA sequencing (scRNA-seq) (Tasic et al., 2016; Tasic et al., 2018) have defined cell types based on genome-wide gene expression and unsupervised clustering. In mouse cortex, we defined more than 100 cell types (Tasic et al., 2018), which were organized in a taxonomy with two major neuronal classes (GABAergic and glutamatergic) divided into many subclasses (groups of related cell types). This characterization included the discovery of many new marker genes for all levels of the taxonomy, but experimental access to cell types still largely depends on transgenic lines. Here, we provide a high-quality dataset for single-cell version of the Assay for Transposase-Accessible Chromatin (scATAC-seq) for the adult mouse cortex. Then we demonstrate that a combination of scRNA-seq and scATAC-seq can be used to identify functional enhancer elements with expected specificity. These elements can be introduced into recombinant adeno-associated viruses (AAVs) to generate cortical cell class- or subclass-specific viral labeling tools. These tools can be delivered using minimally invasive retro-orbital injections to consistently label genetically defined cell populations in mice (**Figure 1**). Moreover, these enhancer viruses can be co-delivered into dual reporter mice to independently label different cell classes within the same animal. Finally, to enable even greater flexibility and experimental simplicity for combinatorial cell type labeling, and future enhancer testing, we generated a novel transgenic mouse line, *Ai213*, which encodes three fluorescent reporters gated independently by three different recombinases, Cre, Flp and Nigri, in a single genomic locus. When used in combination with enhancer viruses, this new line enables efficient and flexible investigation of cell types in the mouse nervous system.

**Figure 1.**
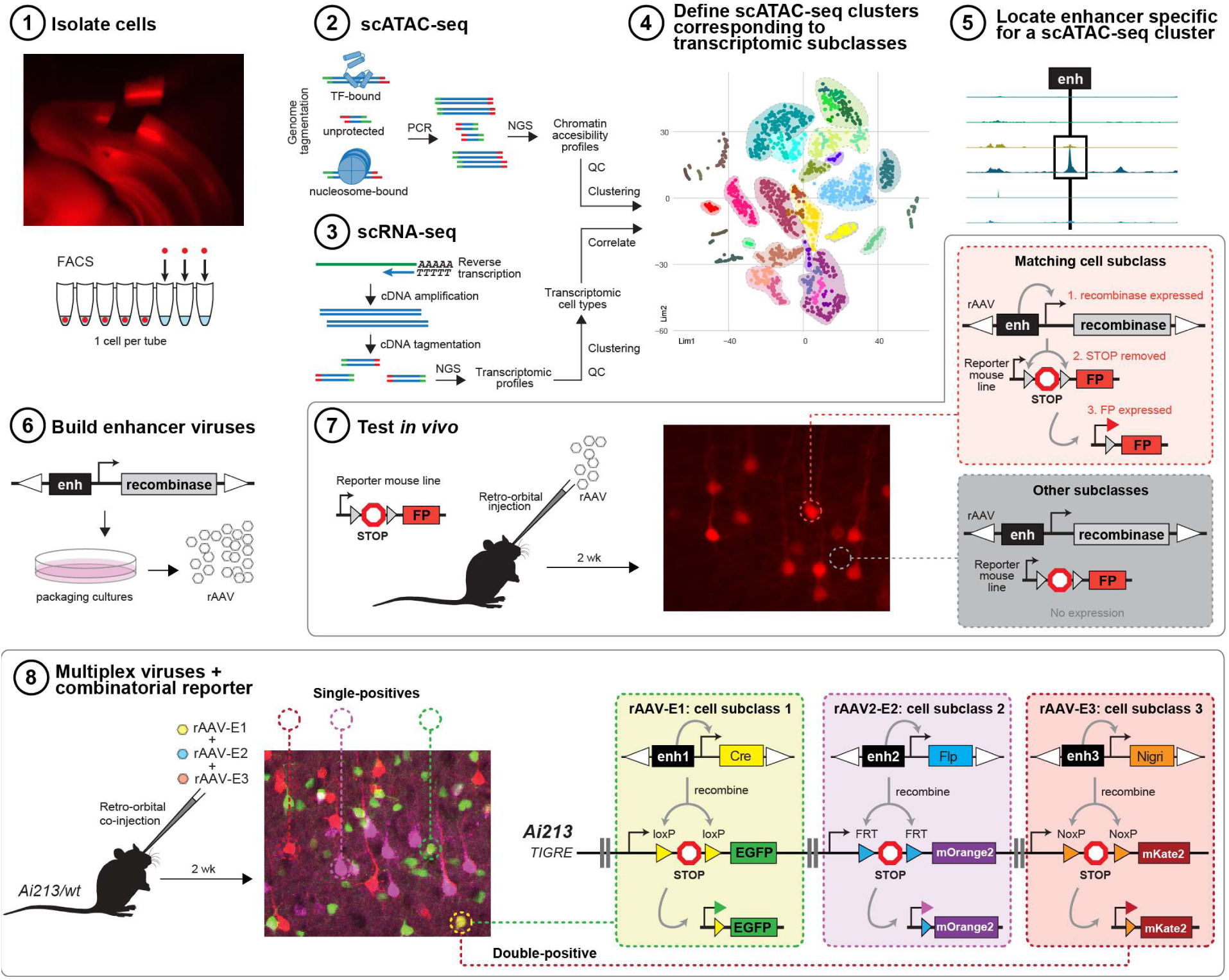
Overview of enhancer discovery for viral tool development. **1-4)**To build cell type-specific labeling tools, we isolated cells from adult mouse cortex, performed scATAC-seq, clustered samples, and compared them to scRNA-seq datasets to assign identity to the clusters. **5)** We then pooled single cells matching a transcriptomic type or subclass and searched the genome for specific putative enhancers. **6)** These regions were cloned upstream of a minimal promoter to drive a fluorescent protein or a recombinase in an AAV genomic backbone, which was used to generate recombinant AAVs. **7)** These viral tools were delivered by retro-orbital or stereotaxic injection to label specific cortical populations. We tested the enhancer viruses for specificity to assess if expected cell types were labeled. In principle, in cells with a matching cell type, enhancers recruit their cognate transcription factors to drive cell type-specific expression. In other cells, viral genomes are present, but transcripts are not expressed. **8)** To label up to three different types of cells with minimal mouse breeding, three different enhancer viruses encoding three different recombinases were mixed and delivered to a new triple-recombinase reporter line, *Ai213*.

## Results

### Single-cell ATAC-seq of adult mouse cortex

We isolated individual neuronal and non-neuronal cells from transgenically labeled mouse cortex by FACS (**Figure S1**) and examined them using scATAC-seq (**Figure 1**) (Buenrostro et al., 2015; Cusanovich et al., 2015). To generate scATAC-seq data that would be directly comparable to our recently published scRNA-seq dataset (Tasic et al., 2018), we focused our dissections on primary visual cortex (VISp) for glutamatergic cell types, but allowed broader cortical sampling for GABAergic cell types. This strategy is based on our observation that GABAergic cell types are shared across the anterior-posterior poles of mouse cortex, whereas the glutamatergic cell types differ between cortical regions (Tasic et al., 2018). To access both abundant and rare cell types, we utilized 25 transgenic Cre or Flp recombinase-expressing driver lines or their combinations crossed to appropriate reporter lines (**Figure S2** and **Table S1**), many of which we previously characterized by single-cell RNA-seq (Tasic et al., 2018). This strategy allowed us to use the same method to interrogate both abundant and rare cell types. In addition, to selectively examine neurons with specific projections, we performed ATAC-seq on cells retrogradely labeled via injections of recombinase-expressing viruses into recombinase reporter mice (Retro-ATAC-seq; **Figure S2, Table S2**, and **Table S3**). Retro-ATAC-seq cells were collected only from VISp.

In total, we collected 3,603 single cells from 25 driver-reporter combinations in 60 mice, 126 retrogradely labeled cells from injections into 3 targets across 7 donors, and 96 cells labeled by one retro-orbital injection of a viral tool generated in this study (**Table S1**). After FACS, individual cells were subjected to ATAC-seq, and were sequenced in 60-96 sample batches (see **STAR Methods**). Our method yielded scATAC-seq datasets of comparable or better quality than previously published scATAC-seq data (Buenrostro et al., 2015; Cusanovich et al., 2015; Pliner et al., 2018) (**Figure S3**). We performed quality-control (QC) filtering to select 2,416 samples with >10,000 uniquely mapped paired-end fragments (median fragment count per cell = 113,184), >10% of which were longer than 250 base pairs (bp), and with >25% of fragments overlapping high-depth cortical DNase-seq peaks generated by ENCODE (Yue et al., 2014) (**Figure 2A, Figure S2**, and **Table S4**).

**Figure 2.**
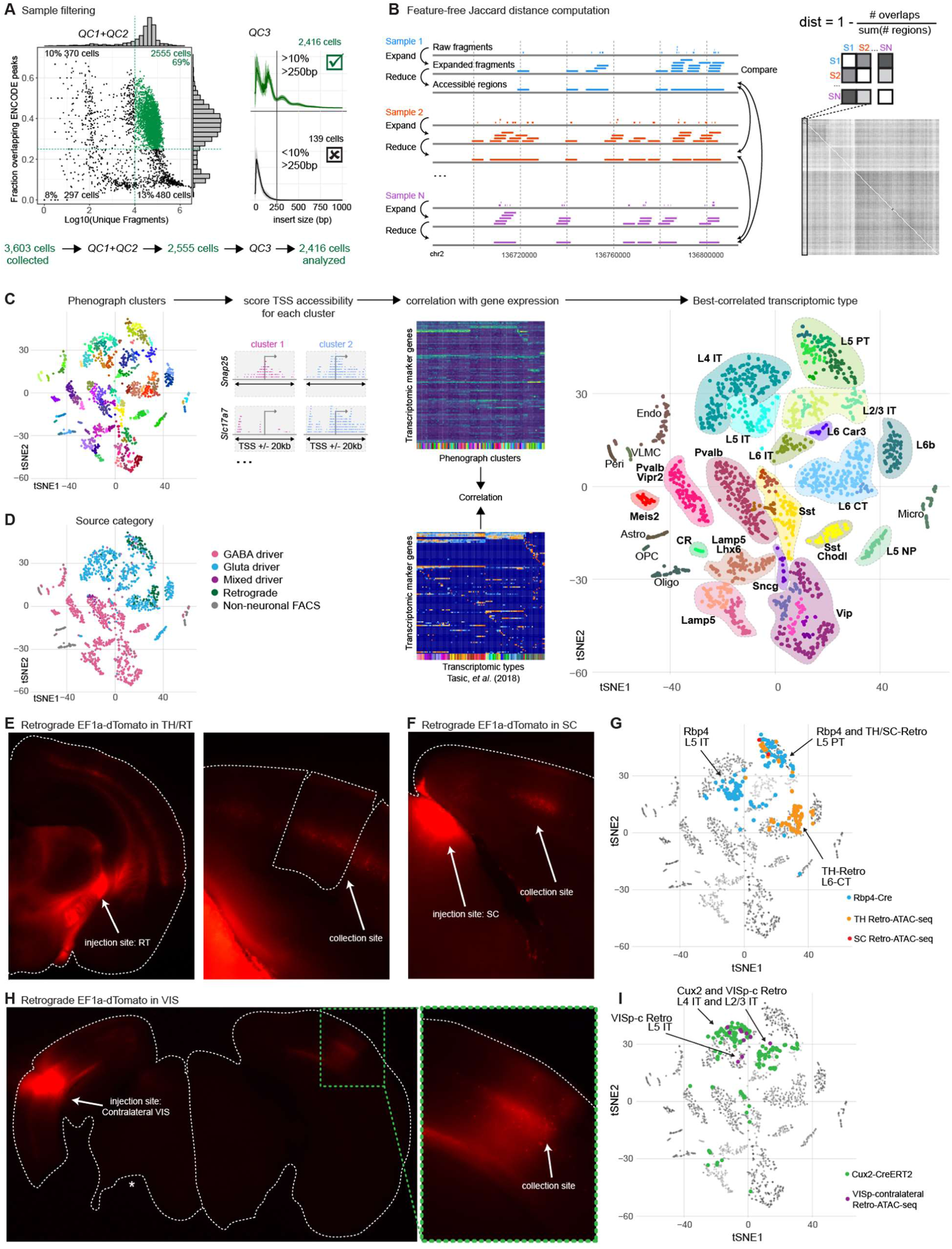
Identification of cell classes, subclasses and types in scATAC-seq data by correlation with scRNA-seq. (**A**) Quality control criteria for scATAC-seq. For analysis, we retained samples with >10,000 uniquely mapped fragments (QC1) that overlapped ENCODE whole-cortex DNase-seq peaks with > 25% of fragments (QC2), and which had nucleosomal fragment size structure identified by > 10% of all aligned fragments with an insert size > 250bp (QC3). (**B**) Samples that passed QC were prepared for clustering by downsampling to 10,000 unique fragments, extending fragment lengths to 1 kb, and reducing overlapping fragments to identify accessible regions. Samples were then compared by computing the number of overlaps between accessible regions, and a Jaccard distance was computed. These distances were used as input for *t-*SNE projection. (**C**) Samples were clustered in *t-*SNE space using the RPhenograph implementation of Jaccard-Louvain clustering. To identify the cell types within these clusters, cells from each cluster were pooled, and we counted the number of fragments within 20 kb of each TSS. We then selected marker genes for transcriptomic clusters from Tasic *et al.* (2018) (Tasic et al., 2018), and performed correlations between TSS accessibility scores and log-transformed gene expression. The scRNA-seq cluster with the highest correlation score was assigned as the identity for each Phenograph cluster, and clusters with the same transcriptomic mapping were combined for downstream analyses. (**D**) *t-*SNE as in (**C**) with samples labeled according to the predominant cell class of their source driver line or experimental source. Cells from the same class are grouped as expected. (**E**) Native fluorescence images of coronal sections dissected for scATAC-seq sample collection from a mouse with retrograde labeling of projections to the thalamus (TH). Cells were labeled by injection of CAV-Cre into TH of an *Ai14* mouse. The left panel shows the full hemisphere near the injection site, while the left panel shows the cortical region (VISp) containing retrogradely labeled cell bodies that was used for collection. RT, reticular nucleus of the thalamus, a thalamic sub-region to which L5 PT cells project. (**F**) Native fluorescence image of a mouse dissected for scATAC-seq sample collection from a SC retrograde experiment. (**G**) Labeling of scATAC-seq cells in *t*-SNE space to highlight cells collected using *Rbp4-Cre* (Rbp4, blue), which labels both L5 IT and L5 PT neurons; using retrograde labeling from the lateral reticular nucleus of the thalamus (RT) to VISp (orange); or retrograde labeling from superior colliculus (SC) to VISp (red). As expected, cells isolated from *Rbp4-Cre* were present in L5 IT and L5 PT ATAC-seq clusters, but not in the L5 NP cluster (Tasic et al., 2018). Cells labeled by retrograde injections into thalamus (TH) were present in L5 PT and L6 CT clusters, but not L5 IT or L5 NP. Retrograde injection into superior colliculus (SC) labeled cells in L5 PT. (**H**) Native fluorescence images of a mouse dissected for scATAC-seq sample collection from a VISp-contralateral retrograde experiment. In this example, cells were collected only from the hemisphere opposite (R) the single-hemisphere injection site (L). Asterisk: tissue lost in vibratome sectioning. The lower panel shows a closer view of the collection site. (**I**) As for (C), but highlighting cells labeled using *Cux2-CreERT2* (Cux2, green), which labels both L2/3 IT and L4 IT neurons, and cells with retrograde labeling from VISp in the contralateral hemisphere (VISp-c, purple). As expected, cells labeled by VISP-c injection were found in L4 IT, L5 IT, and L2/3 IT scATAC-seq clusters.

### Identification of cell classes and subclasses in scATAC-seq data

To identify putative cell class- and subclass-specific regulatory elements, we clustered the scATAC-seq data using a feature-free method for computation of pairwise distances (**Figure 2B** and **STAR Methods**). These distances were used for principal component analysis (PCA) and *t-*distributed stochastic neighbor embedding (*t-*SNE), followed by Phenograph clustering (Levine et al., 2015) (**Figure 2C** and **STAR Methods**). This clustering segregated cells into multiple clusters as expected based on previous transcriptomic analyses (**Figure 2D**). For quantitative comparison to scRNA-seq data at the cluster level, we computed accessibility scores for each gene by counting the number of reads in each cluster that were found near each RefSeq transcription start site (TSS ± 20 kb). The accessibility scores were then correlated with median marker gene expression values for each VISp scRNA-seq cluster from Tasic *et al.* (Tasic et al., 2018) and scATAC-seq cluster identity was assigned based on the scRNA-seq cluster with the highest correlation (**Figure 2C** and **STAR Methods**). For each driver line, we found that subclass-level composition in scATAC-seq data was similar to the one determined based on scRNA-seq data (**Figure S4**).

Transcriptomic types of glutamatergic neurons in the cortex preferentially reside in specific layers and project to specific brain areas. In VISp, intratelencephalic (IT, also called cortico-cortical) neurons reside in all layers except L1 and send axon projections to other cortical regions. Pyramidal tract (PT, also called extra-telencephalic or subcerebral projection neurons) neurons reside in L5 and project subcortically to thalamus and tectum, whereas an additional subclass of subcortically-projecting neurons, termed cortico-thalamic (CT) neurons, reside in L6 and project to thalamus(Harris et al., 2019; Tasic et al., 2018). Previously, we associated transcriptomic identity with neuronal projection patterns by Retro-seq, which combines retrograde tracing with scRNA-seq (Economo et al., 2018; Tasic et al., 2018). To examine if our cell-type assignment for ATAC-seq data is congruent with neuronal projection properties, we performed scATAC-seq on retrogradely labeled neurons (Retro-ATAC-seq, **Figure 2E-I**). We injected a virus into a cortical or subcortical projection target and processed cells from VISp as previously described by our ATAC-seq protocol (see **STAR Methods**). We focused on injections that would differentiate subcortically projecting neurons (project from VISp to thalamus, TH) from cortico-cortically projecting neurons (project to contralateral VISp, VISp-c). Consistent with our annotation of ATAC-seq clusters based on comparison to scRNA-seq, cells labeled by injections into TH (**Figure 2E**) and SC (**Figure 2F**) clustered with cells labeled as L6 CT and L5 PT subclasses (**Figure 2G**), whereas injections into VISp-c (**Figure 2H**) labeled cells in the L2/3 IT, L4 IT, and L5 IT subclasses (**Figure 2I**). Likewise, our subclass assignments for the L5, L2/3, and L6 to our ATAC-seq clusters agreed with the transgenic sources from which the cells were derived. For example, the cells derived from Rbp4-Cre were mostly present in L5 IT and L5 PT subclasses (**Figure 2G**), whereas cells derived from Cux2-CreERT2 fell into L2/3 IT and L4 IT subclasses (**Figure 2I**) (Gray et al., 2017).

### Identification of putative subclass-specific enhancers

To identify putative enhancers specific for cell classes (i.e. GABAergic and glutamatergic), subclasses (i.e. related subpopulations of GABAergic and glutamatergic cells) or types (i.e. one distinct population of cells within a subclass), we aggregated the data from different scATAC-seq clusters for peak calling and examination of chromatin accessibility patterns (**Figure 3, S5**). These aggregated scATAC-seq profiles corresponded well to published ATAC-seq from cortical cell populations (Gray et al., 2017; Mo et al., 2015) (**Figure S6**).

**Figure 3.**
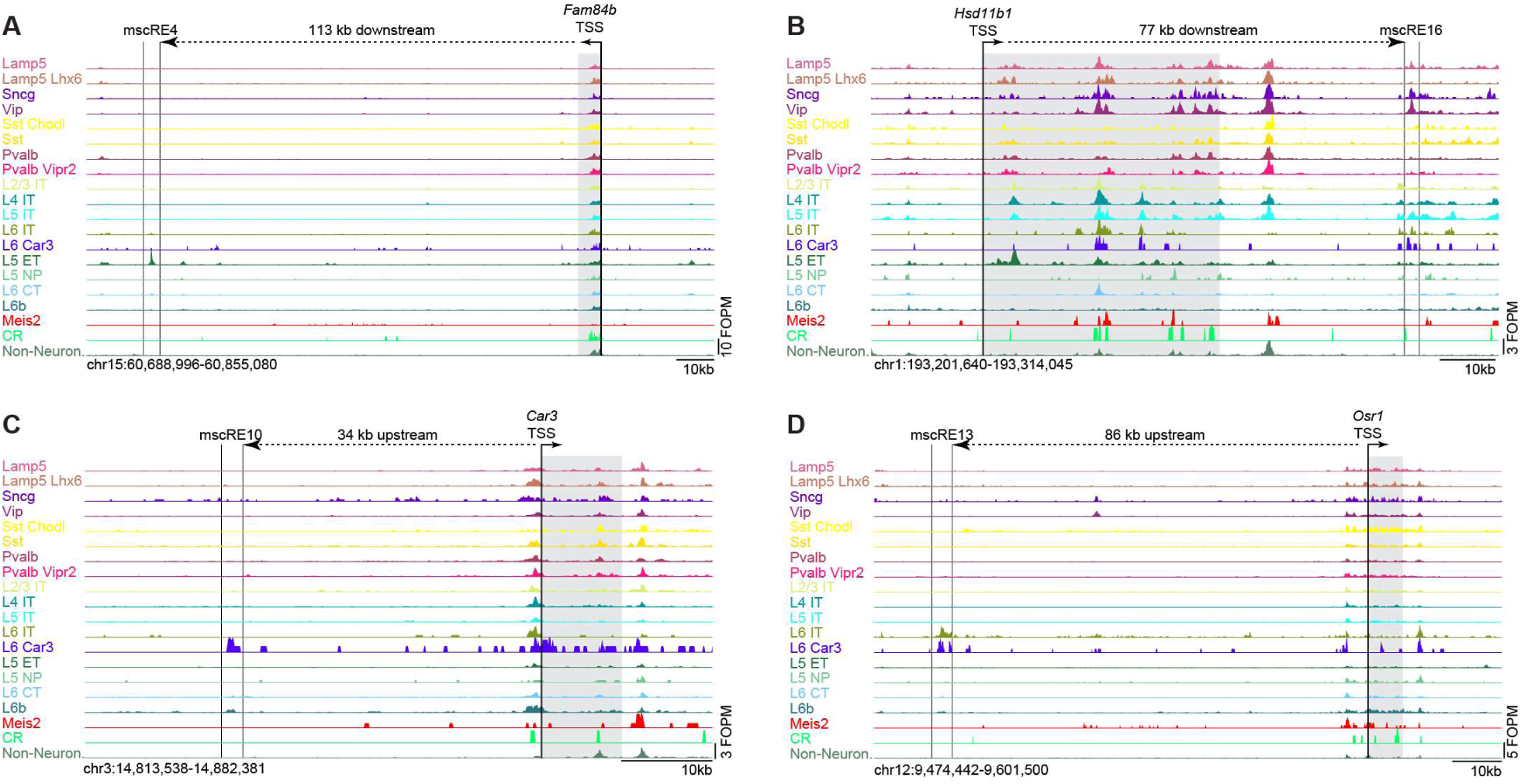
scATAC-seq peak selection defines mscREs. Chromatin accessibility in clusters based on single cell ATAC-seq data for select genomic regions containing mscRE4, mscRE16, mscRE10 and mscRE13. The nearby gene, which is the likely target of each enhancer is shaded, with the transcription start site (TSS) and the direction of transcription designated by arrow. The distance between each TSS and mscRE is indicated. For a complete set of mscREs examined in this study, see **Figure S5**.

To identify putative subclass-specific enhancers, which we call mouse single-cell regulatory elements (mscREs), we interrogated the genomic neighborhoods of subclass-specific marker genes identified in our previous transcriptomic study (Tasic et al., 2018). We focused on mscREs that were 500-600 bp-long, and were preferentially accessible in L5 PT, L5 IT, L6 Car3, or L6 IT subclasses compared to all other subclasses, and had high sequence conservation (Siepel et al., 2005) (**Figure S5; Table S5**). Genome accessibility profiles per ATAC-seq cluster for 16 such elements are shown in **Figure S5**, including 4 that were extensively characterized in this study: mscRE4, mscRE10, mscRE13 and mscRE16 (**Figure 3**).

### Enhancer-driven fluorophores for cell subclass labeling

To functionally test mscREs, we initially cloned them upstream of a minimal β-globin promoter (Yee and Rigby, 1993) driving fluorescent proteins SYFP2 or EGFP in a recombinant AAV genome (**Figure 3A; Table S7**). To enable delivery of these viruses by retro-orbital injection, the recombinant genomes were packaged into the PHP.eB serotype of AAV, which can cross the blood-brain barrier (Chan et al., 2017). We screened nine mscREs for the L5 PT subclass, two for L5 IT, three for L6 Car3, and four for L6 IT (**Table S7** and **Figure S7**). Two weeks after retro-orbital injection of virus, we analyzed native or immunohistochemistry-enhanced fluorescence in brain slices. Eight of these enhancers labeled cells in L5 and two in L6 (**Figure S7**). We selected two from each layer for additional screening and experiments (L5: mscRE4 and mscRE16; L6: mscRE10 and mscRE13).

To determine the specificity of the enhancer viruses driving fluorophores, we performed additional retro-orbital injections of viruses, dissected L5 of VISp, isolated labeled cells by FACS, and performed SMART-Seq v4-based scRNA-seq as described previously (**Figure 4A**) (Tasic et al., 2018). We classified these scRNA-seq expression profiles by bootstrapped centroid mapping to our reference transcriptomic taxonomy (see **STAR Methods**) (Tasic et al., 2018). We observed labeling in L5, and found that the mscRE4-SYFP2 virus yielded >91% specificity for L5 PT cells within L5 (**Figure 4B,C** and **Table S8**). Using the same strategy, we collected cells labeled by mscRE4-SYFP2 for scATAC-seq. As previously observed by scRNA-seq analysis, 55 of 61 high-quality mscRE4 scATAC-seq profiles clustered together with other L5 PT samples (90.2%, **Figure S8**). We also confirmed labeling of L5 PT cells by electrophysiological characterization of labeled vs. unlabeled cells in the mouse cortex (**Figure 4D-E** and **Figure S9**). Cells labeled by the mscRE4-SYFP2 virus had physiological characteristics of thick-tufted cortical L5 PT neurons (high resonance frequency and low input resistance), whereas unlabeled cells more closely matched L5 IT neurons (Baker et al., 2018; Dembrow et al., 2010). These data collectively demonstrate that L5 PT neurons are labeled and can be examined reliably by the mscRE4-SYFP enhancer virus.

**Figure 4.**
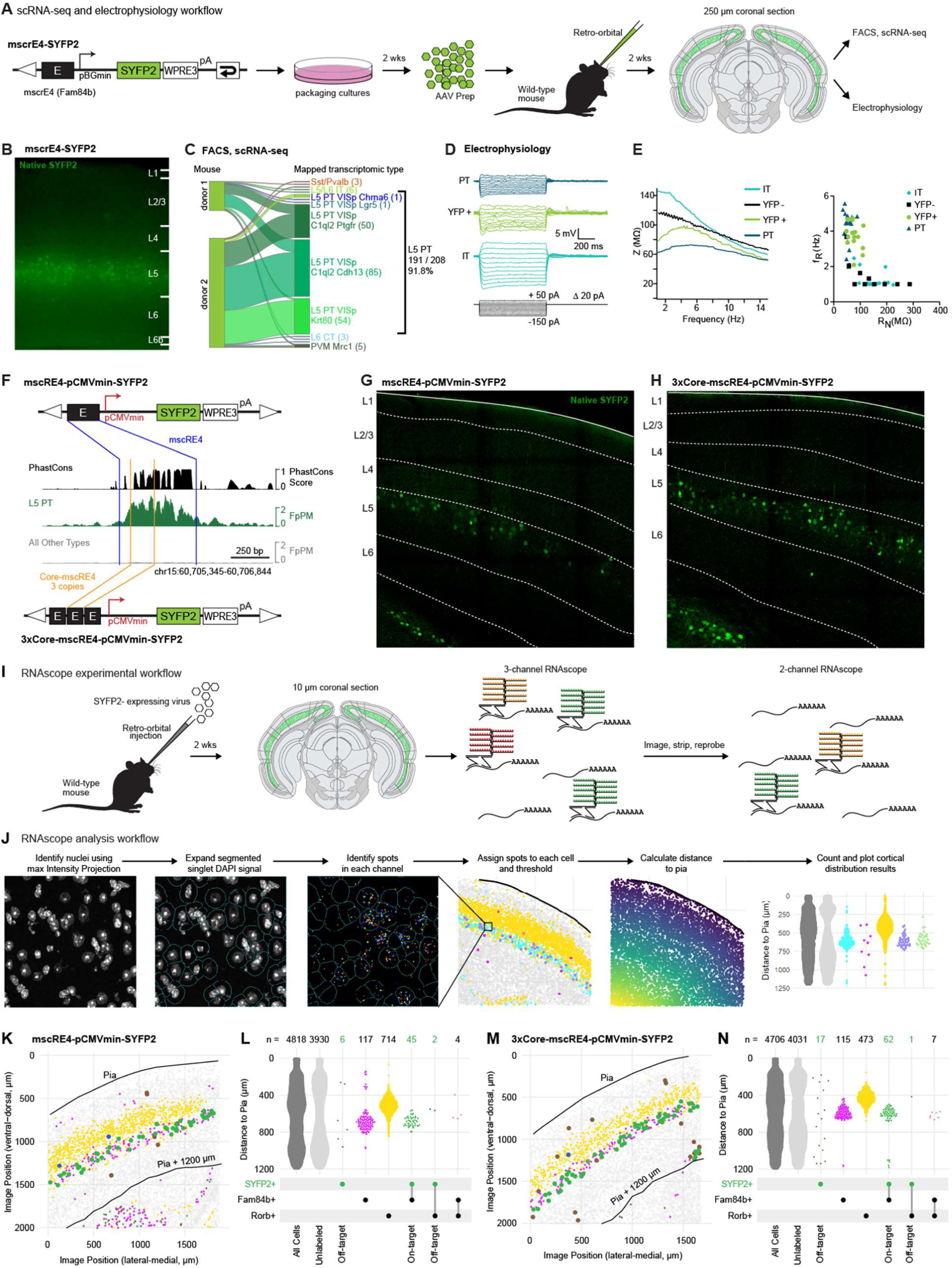
Direct fluorophore labeling of L5 PT neurons by enhancer viruses. (**A**) Experimental workflow for testing the enhancer virus containing putative *Fam84b* enhancer, mscRE4, in a self-complementary AAV backbone (scAAV) with a beta-globin minimal promoter (pBGmin) driving SYFP2. WPRE3: short woodchuck hepatitis virus posttranscriptional regulatory element; pA: polyadenylation site. After packaging, purification, and titering, the virus was retro-orbitally injected into a wild-type mouse, and tissue examined two weeks later by scRNA-seq or electrophysiology. (**B**) Two weeks post-injection, distinct labeling was seen in L5, which was microdissected, dissociated, and sorted by FACS for SYFP2+ cells for scRNA-seq. scRNA-seq data (n = 219 cells) from n=2 animals, were mapped to the Tasic *et al.* (2018) cell type reference. 91.8% of all profiled cells mapped to L5 PT cell types. (**D**) Electrophysiological characterization of cells in cortical slices labeled with the mscRE4-SYPF2. Example voltage responses to a series of hyperpolarizing and depolarizing current injections for a YFP+ and a YFP– neuron. Example voltage responses obtained from unlabeled PT-like and IT-like neurons are also shown for reference. (**E**) Example impedance amplitude profiles obtained from a YFP+ and a YFP– neuron in VISp (Left). For comparison, impedance amplitude profiles from an unlabeled PT-like and an IT-like neuron from somatosensory cortex are also shown. Resonance frequency (*f*_R_) is plotted as a function of input resistance (R_N_, right). (**F**) Schematics of viral genomes constructed to evaluate concatenation of mscRE4 in order to increase SYFP2 expression levels. Note that compared to construct in (A), CMV minimal promoter, and non-self-complementary AAV backbone was used. Center, Tn5 transposon footprinting of the mscRE4 enhancer region, bounded by blue bars. A subset of this region, termed Core-mscRE4, was selected based on accessibility and conservation, bounded by orange bars. Tracks show PhastCons conservation scores scaled between 0-1 (black), L5 PT scATAC-seq accessibility (green), and non-L5 PT scATAC-seq accessibility (gray). Accessibility tracks are scaled to Footprints Per Million Reads (FpPM). (**G-H**) Native fluorescence imaging 3 weeks post-injection of mscRE4-pCMVmin-SYFP2 constructs packaged in PHP.eB capsids and delivered retro-orbitally. Two viruses: single copy, full-length mscRE4 enhancer in pCMVmin-SYFP2 construct (H) and the 3xCore-mscRE4-pCMVmin-SYFP2 construct (G) were titer-matched for injections, and imaging was performed on the same microscope with identical exposure settings on the same day. (**I**) Experimental workflow for evaluation of the viruses in F for specificity and completeness of labeling *in situ* by RNAscope. Viruses were retro-orbitally injected into wild-type mice. Two weeks later, two rounds of RNAscope were performed on fresh-frozen brains. (**J**) RNAscope analysis workflow included identification of nuclei and expansion of nuclear boundaries to mimic cell boundaries. Fluorescent spots corresponding to specific mRNA molecules in each channel were counted and assigned to individual cells. Threshold was applied to each probe to account for background signal, and positive cells were plotted along distance from pia. (**K**) In-tissue positions of cells labeled by RNAscope in a cortical region from an animal injected with mscRE4-pCMVmin-SYFP2. Each spot corresponds to a single cell, and is colored according to the probe combination that labeled each cell: gray, unlabeled; bright green, *SYFP2*+ only; magenta, *Fam84b*+ only; yellow, *Rorb*+ only; dark green, *SYFP2*+ and *Fam84b*+; pale green, *Fam84b*+ and *Rorb*+; orange, *Fam84b*+ and *Rorb*+. *SYFP2*+ points are enlarged for visibility. (**L**) Data from (K) plotted as cell counts (top) relative to pia for cells positive for each combination of probes. Points are jittered on the x-axis using quasirandom positioning. Probe combinations and on/off-target indications are shown below the plot. *SYFP2*+ labeling is shown in green for emphasis. *Scnna1* and *Hsd11b1* were used to delineate L4 and L5 boundaries (not shown, counts in **Table S9**). (**M**) Same as (K) but for 3xCore-mscRE4-pCMVmin-SYFP2 virus. (**N**) Data from (M) plotted like in (L).

Stereotaxic injections can be used for spatially targeted labeling of cell types by providing virus to a specific location in the brain, rather than the whole brain. We tested injection of the mscRE4 and mscRE16 fluorophore viruses directly into VISp and found that we could achieve bright labeling using stereotaxic injection. However, specificity of labeling was considerably lower than with retro-orbital delivery (mscRE4: 31.7% and 38% on target at 50 nL and 25 nL injections, respectively), likely reflecting the larger multiplicity of infection achieved by direct brain injections (**Figure S10).**

While stereotaxic delivery enhanced the brightness of labeling, it decreased the specificity, and cannot provide labeling throughout the brain, but only in the tissue surrounding the injection site. We sought to enhance the efficacy of retro-orbital labeling by optimizing the design of the AAV viral genomes using two complementary approaches: use of a stronger minimal promoter, and integration of multiple copies of mscRE4 to increase the amount of fluorophore expression without reducing cell type specificity. As an alternative to the β-globin minimal promoter used in our initial constructs, we selected the cytomegalovirus (CMV) minimal promoter, which was empirically determined to mediate stronger enhancer-driven transgene expression (data not shown). To ensure additional copies of mscRE4 would fit in the limited space of the AAV genome with diverse gene expression cargo, we selected a short core sequence (155 bp) from the mscRE4 enhancer and inserted 3 copies in a construct driving SYFP2 (**Figure 4F**). We found that the 3xCore increased native fluorescence in L5 PT cells above that with a single copy of the full mscRE4 sequence (**Figure 4G-H**).

To assess the specificity and completeness of L5 PT cell labeling *in situ*, we used single molecule RNA fluorescence *in situ* hybridization with RNAscope (**Figure 4I**). RNAscope was performed using two rounds of labeling with a total of 5 probes: one for *SYFP2*, which is expressed in cells labeled by the enhancer viruses; and 4 endogenous mRNAs that were selected based on scRNA-seq data to distinguish L5 IT from L5 PT cells. In the first round, we detected *SYFP* mRNA and *Fam84b*, which is expressed in L5 PT cell types, as well as *Rorb*, which is expressed in L4 and most L5 IT cell types. In the second round, to better delineate the boundary between L4 and L5, we detected *Scnn1a*, which labels L4 IT cell types and *Hsd11b1*, which is expressed in L5 IT cells. We identified nuclei using maximum intensity projection of the DAPI channel and expanded them digitally to approximate the cytoplasm (**Figure 4J**). Spots from each imaging channel were then counted and assigned to each cell, and thresholds for each channel were applied based on a channel-specific background count to define cells that express examined transcripts (**Figure 4K,M** and **Table S9**). Position of each cell was also assessed as the distance of the center of the nucleus from the pial surface in each image (**Figure 4L,N**). We found that the CMV minimal promoter had 84% on-target specificity in cortex n = 8,363 total cells examined; 68 SYFP+; n = 2 sections; results of one experiment are shown in **Figure 4K** with counts for both in **Figure 4L** and **Supp. Table 9**). Three copies of the mscRE4 enhancer core further increased SYFP expression (**Figure 4H,G**) with a modest loss of specificity: 78% of *SYFP2*+ cells in cortex were also *Fam84b*+ (n = 4,706 total cells examined; 80 SYFP+ cells; n = 1 section; shown in **Figure 4M,N** and **Supp. Table 9**).

### Viral enhancer-driven recombinases

To determine whether enhancer viruses expressing an exogenous recombinase or tetracycline could be combined with transgenic reporter lines to enable high and reliable expression of many other genes (e.g., activity reporters, opsins), some of which are too large to fit into the limited genome size of recombinant AAVs (Daigle et al., 2018; Madisen et al., 2015), we generated recombinase or the tetracycline-dependent transcriptional activator (tTA2)-expressing enhancer viruses. We cloned mscRE4 into viral constructs containing a minimal β-globin promoter driving a destabilized Cre (dgCre), a mouse codon-optimized Cre (iCre), a mouse codon-optimized Flp (FlpO), or tTA2, and packaged them into AAV PHP.eB serotype (**Figure 5A**). Viruses were delivered by retro-orbital injection into conditional reporter mice for each recombinase or transcription factor: *Ai14* for dgCre and iCre (Madisen et al., 2010); *Ai65F* for FlpO and *Ai63* for tTA2 (Daigle et al., 2018). Native tdTomato fluorescence for each enhancer virus was assessed two weeks post-injection in fixed brain tissue (**Figure 5A**). tTA2- and FlpO-expressing enhancer viruses achieved the highest specificity for L5, with tTA2 giving the most restricted labeling pattern. The iCre-expressing virus labelled excitatory types mainly in L5 and L6, with sparser expression observed in L2/3, whereas the dgCre-expressing virus gave widespread non-specific labeling in cortex and was not pursued further.

**Figure 5.**
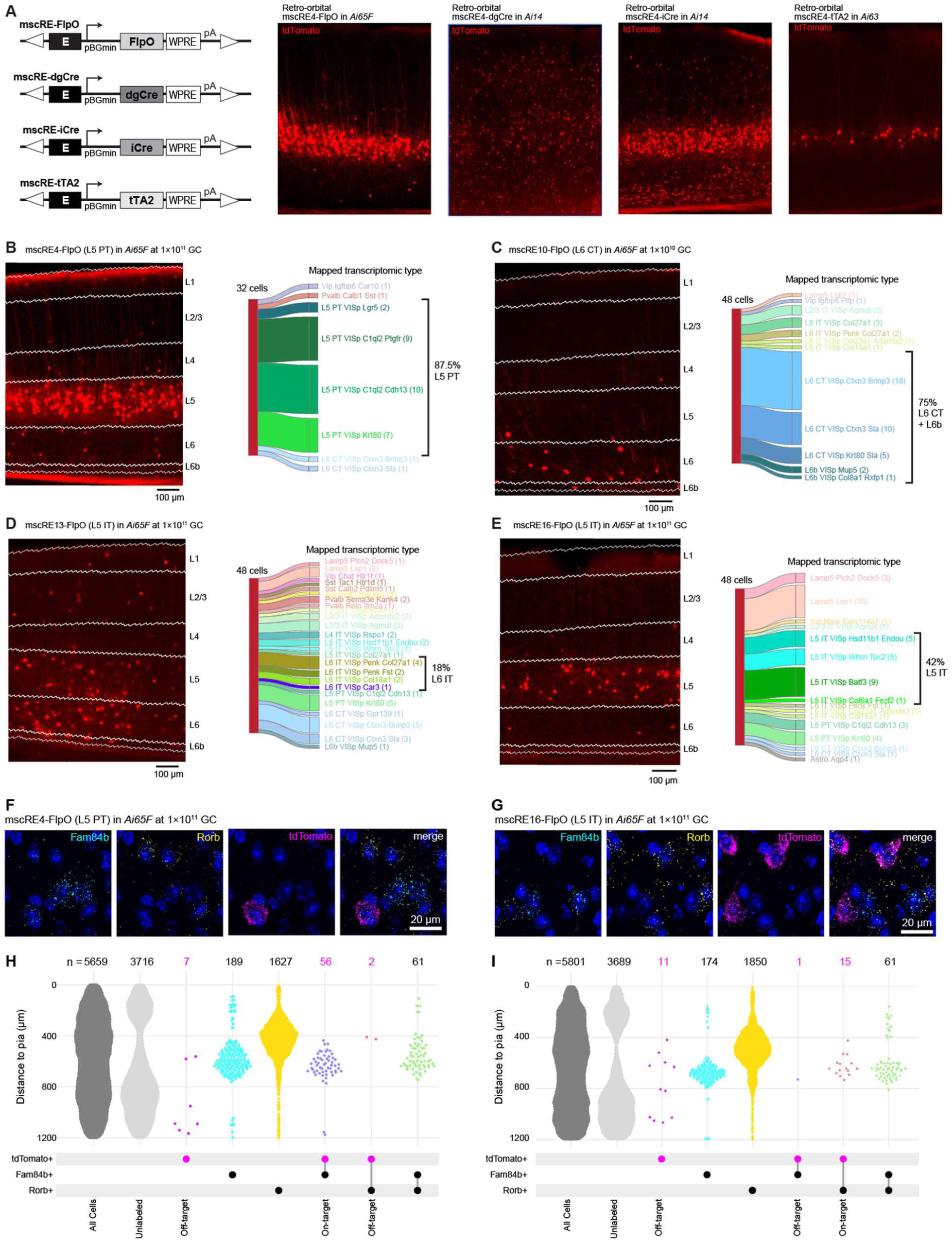
Cell subclass labeling by enhancer-driven recombinase or transcription factor viruses. (**A**) Schematics of viral genomes used for comparison of enhancer-driven FlpO, dgCre, iCre, or tTA2. E: enhancer, pBGmin: minimal beta-globin promoter; WPRE: woodchuck hepatitis virus posttranscriptional regulatory element; pA: polyadenylation site. The viral genomes were packaged into PHP.eB-serotype rAAVs, and injected retro-orbitally into reporter mice. Images show native fluorescence in visual cortex 2 weeks post-injection. (**B**) Cortical imaging of a mscRE4-FlpO injection shows labeling throughout L5 of the posterior cortex. Layer overlays from the Adult Mouse Allen Brain Reference Atlas shows labeling restricted primarily to L5. *tdTomato*+ cells were dissected from the full cortical depth and were collected by FACS for scRNA-seq. Transcriptomic profiles were mapped to reference cell types from Tasic *et al.* (2018) (Tasic et al., 2018). 87.5% of cells (28 of 32) mapped to L5 PT cell types. (**C**) Cortical imaging of mscRE10-FlpO injection shows labeling in L6 of the cortex. scRNA-seq of *tdTomato*+ cells shows that L6 CT and L6b cell types are the most frequently labeled subclasses of neurons (75%, n = 36 of 48). (**D**) Cortical imaging of mscRE13-FlpO injection shows deep-layer enriched labeling of cells. However, scRNA-seq of *tdTomato*+ cells shows that there is not strong enrichment for any single cell subclass, and only 18% of cells are labeled as predicted (n = 9 L6 IT cells of 48 labeled cells) (**E**) Cortical imaging of mscRE16-FlpO injection shows labeling in L5 of the cortex. scRNA-seq of *tdTomato*+ cells shows that L5 IT cell types are the most frequently labeled subclass of neurons (42%, n = 20 of 48), but other subclasses are also labeled at this titer (Lamp5, 27%; L6 IT, 6%; L5 PT, 15%). (**F**) Representative images from the cortex of an animal injected with mscRE4-FlpO probed for *Fam84b, Rorb*, and *tdTomato* expression using RNAscope. The same field of cells is shown in all 4 panels, and the scale bar at the right corresponds to all 4 panels. (**G**) Representative images from the cortex of an animal injected with mscRE16-FlpO probed for *Fam84b, Rorb*, and *tdTomato* expression using RNAscope. The same field of cells is shown in all 4 panels, and the scale bar at the right corresponds to all 4 panels. (**H**) Positions relative to the pial surface for cells labeled by injection with mscRE4-FlpO with RNAscope probe combinations: gray, unlabeled; magenta, *tdTomato*+ only; cyan, *Fam84b*+ only; yellow, *Rorb*+ only; purple, *tdTomato*+ and *Fam84b*+; orange, *tdTomato*+ and *Rorb*+; green, *Fam84b*+ and *Rorb*+. Counts of cells with each probe combination are above the plot. Points are jittered on the x-axis using quasirandom positioning. Probe combinations and on/off-target indications are shown below the plot. *tdTomato*+ labeling is shown in magenta for emphasis. (**I**) Positions relative to pia for cells labeled by injection with mscRE16-FlpO and probed with RNAscope. Colors and labeling are the same as in (H).

We applied the same strategy to evaluate mscRE10, mscRE13, and mscRE16 as drivers of FlpO, iCre, and/or tTA2 by retro-orbital injection at two different titers (1×10^10^ and 1×10^11^ total genome copies, GC). We found that the specificity and completeness of labeling depended on both the total GC of virus delivered, the recombinase-reporter combination used in these experiments (**Figure S11**), and age of mice utilized at the time of injection (data not shown). Based on these experiments, we chose a single titer for mscRE4, mscRE10, mscRE13, and mscRE16 FlpO viruses for in-depth characterization, and injected additional animals for brain-wide imaging (**Figure 6A-B, Figure S12**) and scRNA-seq (**Figure 5B-E**).

**Figure 6.**
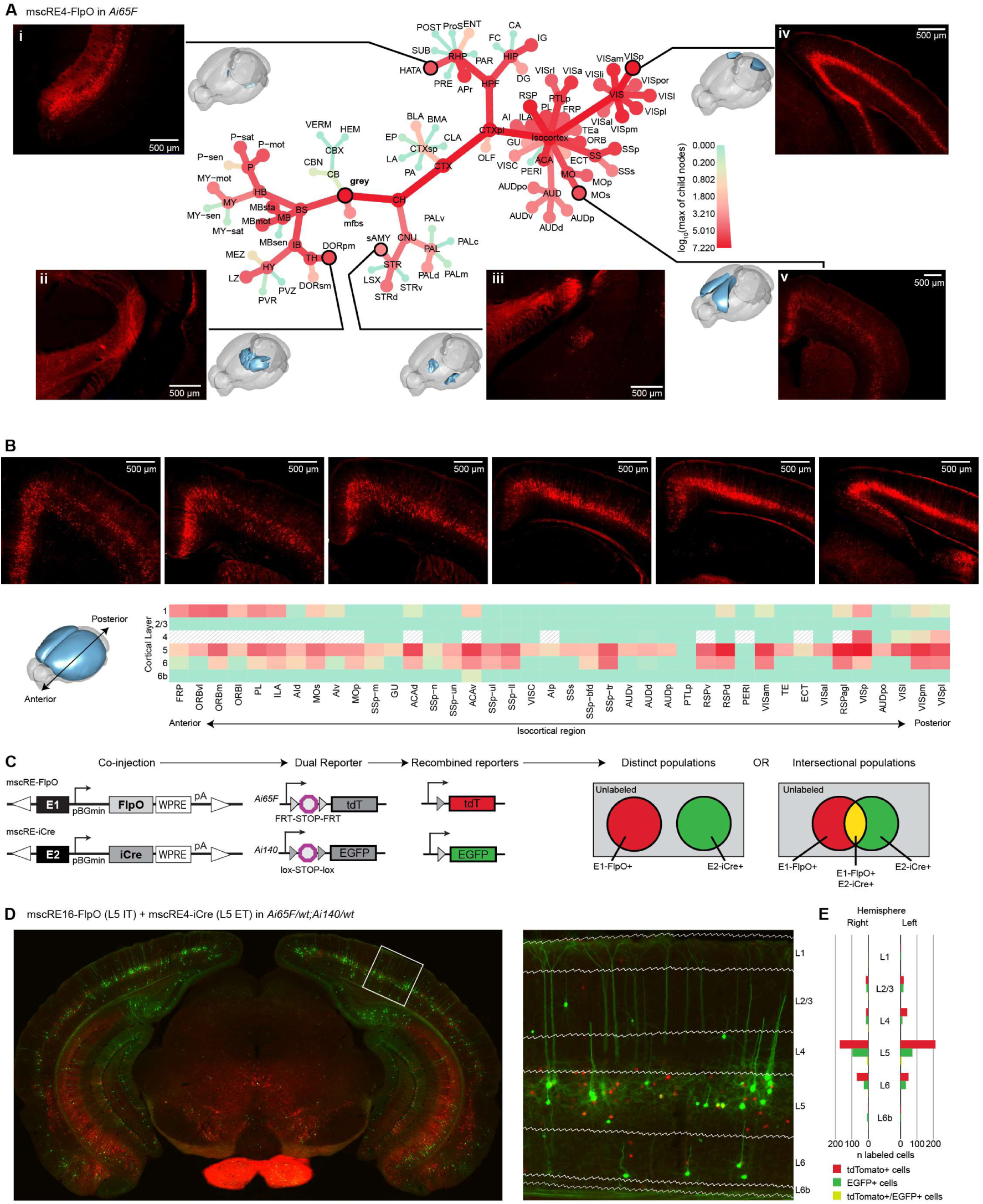
Characterization of brain-wide and combinatorial labeling of cell types. (**A**) Whole-brain two-photon imaging using TissueCyte. Sections throughout the whole brain of an *Ai65F* mouse after retro-orbital injection of mscRE4-FlpO were aligned to the Allen Institute Common Coordinate Framework (CCF), and labeling intensity per region was represented within Allen Brain Atlas structural ontology dendrogram. “Grey” is the root of the plot, representing all grey-matter regions, and each branching represents child structures within each region. The combination of size and color of nodes (see legend) represent the maximum signal found among all children of the nodes, which allows one to follow the tree to the source of high signal within each structure. Insets display selected regions of high or specific signal. Region acronyms correspond to the Adult Mouse Allen Brain Reference Atlas. (**B**) Further division of the isocortical regions in the TissueCyte dataset to the level of cortical layers allows cortex-wide quantification of layer-specific signal. Representative cortical sections from the TissueCyte dataset are shown along the top, from most anterior to most posterior (left to right). The heatmap shows quantification of the signal in each region and layer. Note that agranular regions are thought to lack a typical layer 4 (indicated as white with gray hashing). (**C**) Diagrams showing the use of co-injected recombinase viruses in a dual-reporter system for co-labeling or intersectional labeling of cell types. In this experiment, one virus driving FlpO and a second driving iCre are co-injected into a mouse with genetically encoded Flp-dependent and Cre-dependent reporters. In target cell types, enhancers will drive the recombinases, which will permanently label their target cell types. If the enhancers selected are mutually exclusive, distinct populations will be labeled. If they overlap, intersectional labeling is possible. (**D**) Native fluorescence imaging of an *Ai65F;Ai140* dual-reporter mouse line retro-orbitally injected with mscRE16-FlpO (red) and mscRE4-iCre (green). These enhancers are expected to label mostly mutually exclusive cell types in L5 of the cortex. The region in the white box corresponds the inset image, showing strong labeling of cells in L5. (**E**) Cell counts within each layer for all cortical regions labeled with EGFP (mscRE4; L5 PT), tdTomato (mscRE16; L5 IT), or both in the image in (D).

We found that 3 out of 4 of these viruses had a high degree of layer- and subclass-specificity in the cortex, with 87.5% of cells labeled by mscRE4-FlpO corresponding to L5 PT cells (**Figure 5B**), and 42% of cells labeled by mscRE16-FlpO corresponding to L5 IT cells (**Figure 5E**), with little overlap in cell types labeled. Viruses with mscRE13 proved to be largely non-specific, with only 18% of sequenced cells mapping to the expected L6 IT subclass and labeling of a broad distribution of cortical cell types (**Figure 5C**). We found that 75% of cells labeled by mscRE10-FlpO were L6 CT or L6b cells, as predicted by the scATAC-seq data (**Figure 5D and Figure S5**). However, mscRE10-FlpO did not label L6 IT Car3 cells, despite the proximity of the enhancer to the *Car3* gene and mscRE10 accessibility in this subclass in the scATAC-seq data (**Figure S5**).

We have observed previously that some cell types, especially L5 PT and parvalbumin-expressing cells, are sensitive to isolation needed for scRNA-seq and may be selectively depleted among profiled neurons (Tasic et al., 2018). Therefore, to assess the specificity and completeness of mscRE4-FlpO and mscRE16-FlpO *in situ*, we injected each virus into *Ai65F* reporter mice and analyzed the tissue by RNAscope as described above (**Figure 5F-I, Table S9**). We found that mscRE4-FlpO virus labelling was highly specific for L5 PT cells (83%; 97 of 117 *tdT*+ cells were also *Fam84b*+, **Figure 5H**, counts for both are in **Table S9**), consistent with scRNA-seq results in **Figure 5B**, and very similar to direct labeling by the mscRE4-pCMVmin-SYFP2, as shown in **Figure 4K-L**. Although quite specific, the labeling was incomplete: only 18% of the cells that could in principle be labeled were labeled (97 of 551 *Fam84b*+ cells were also *tdT*+). For the mscRE16-FlpO virus, we found 54% of labeled cells to be L5 IT cells (**Figure 5I, Table S9**). Co-labeling of *tdT*+ with *Fam84b*+ was observed and the off-target rate was found to be 13% for L5 PT cells (6 out of 46 cells **Figure 5I**), consistent with the scRNA-seq results (7 of 48 cells, 15%, **Figure 5E**).

### Characterization of brain-wide labeling of cell types by retro-orbital injection of enhancer viruses

To assess brain-wide labeling specificity, we injected mscRE4-FlpO, mscRE10-FlpO, and mscRE16-FlpO viruses into *Ai65F* animals and performed whole-brain two-photon tomography imaging of native tdTomato fluorescence using the TissueCyte system (mscRE4, **Figure 6A-B**; mscRE10 and mscRE16, **Figure S12**). Images were registered to the Allen Mouse Common Coordinate Framework (CCFv3) for quantification of expression in each region of the CCFv3. For the mscRE4-FlpO virus, we observed distinct labeling in L5 throughout mostly posterior cortex, labelling in L5, L6 and L2/3 cells in more anterior cortex (**Figure 6A**, VISp, **panel iv**; MOs, **panel v**; and **Figure 6B**), and specific expression in several subcortical regions (**Figure 6A**, HATA, **panel i**; sAMY, **panel iii**). As *Ai65F* expresses tdT within the cytoplasm, we also observed strong labeling of axonal tracts and projection targets of L5 PT neurons (e.g., **Figure 6A** DORpm, **panel ii**). TissueCyte imaging revealed that mscRE10 labeling was sparse throughout the cortex and restricted to the cortex and claustrum (**Figure S12A-B**), while mscRE16-FlpO labeled distinct subcortical populations (**Figure S12C**), and had stronger labeling of L5 cells in the primary somatosensory and primary motor cortex regions (**Figure S12D**).

We investigated if retroorbital viral delivery and subsequent neuronal labeling had effect on gene expression by comparing scRNA-seq data from matched virus-labeled and transgene-labeled cell types. We found no significant induction of innate immune genes when virus was delivered retro-orbitally (**Figure S13A-E**). In contrast, and as observed previously (Daigle et al., 2018), we found significant viral volume-dependent innate immune gene induction when the virus was delivered by stereotaxic injections (**Figure S13F-I**). Therefore, brain-wide delivery of AAV by retro-orbital injection does not induce a long-lasting innate immune response, resulting in unperturbed transcriptional profiles of labeled cell types.

### Combinatorial labeling of cell subclasses

To simplify breeding and experimental schemes, we tested if enhancer viruses can be combined with one another and with transgenic reporter lines to label multiple cell types simultaneously in mouse the brain *in vivo* (**Figure 6C**). The enhancer viruses mscRE4-iCre (to label L5 PT cells) and mscRE16-FlpO (to label L5 IT cells) were co-injected retro-orbitally into *Ai65F/wt;Ai140/wt* double transgenic reporter mice (**Figure 6D**) and labelling was visualized in brain sections two weeks post-injection. Here we found largely distinct labeling of L5 PT cells (in green) and L5 IT cells (in red) throughout the cortex and in multiple subcortical regions (**Figure 6D-E)**, demonstrating that multiple enhancer-driven viruses can be used simultaneously to label defined neuronal subclasses in one animal. We also observed a few co-labelled cells in cortex and midbrain (in yellow), showing that both enhancers can be active within a cell.

While the above strategy which utilizes *Ai65F/wt;Ai140/wt* animals enables two color labeling, it still requires the creation of double-transgenic reporter animals. To simplify transgenic requirements and significantly expand the number of cell types that could potentially be labeled within a single animal, we generated *Ai213*, a novel triple-recombinase reporter transgenic mouse line with independent recombinase gating of three different fluorophores in the *TIGRE* locus (Zeng et al., 2008). Generating this complex transgene required re-engineering the *TIGRE* locus to create a new landing pad embryonic stem (ES) cell line with Bxb1 integrase attP sites (Ghosh et al., 2003) to enable site-specific, unidirectional and efficient integrase-mediated cassette exchange in the locus. We then engineered a large (∼26 kb) targeting vector with three independent transgene expression cassettes flanked by Bxb1 integrase attB sites that each contained a recombinase-gated, CAG promoter-driven fluorophore (**Figure 7A**): EGFP is gated by a loxP-STOP-loxP (excised by Cre recombinase), mOrange2 is gated by a FRT-STOP-FRT (excised by Flp recombinase), and mKate2 is gated by a nox-STOP-nox (excised by Nigri recombinase) (Karimova et al., 2016). Bxb1 integrase-mediated cassette exchange into the *TIGRE* locus yielded a single-copy 24 kb-transgene and subsequently, the modified ES cells gave rise to the *Ai213* reporter line (**Methods**).

**Figure 7.**
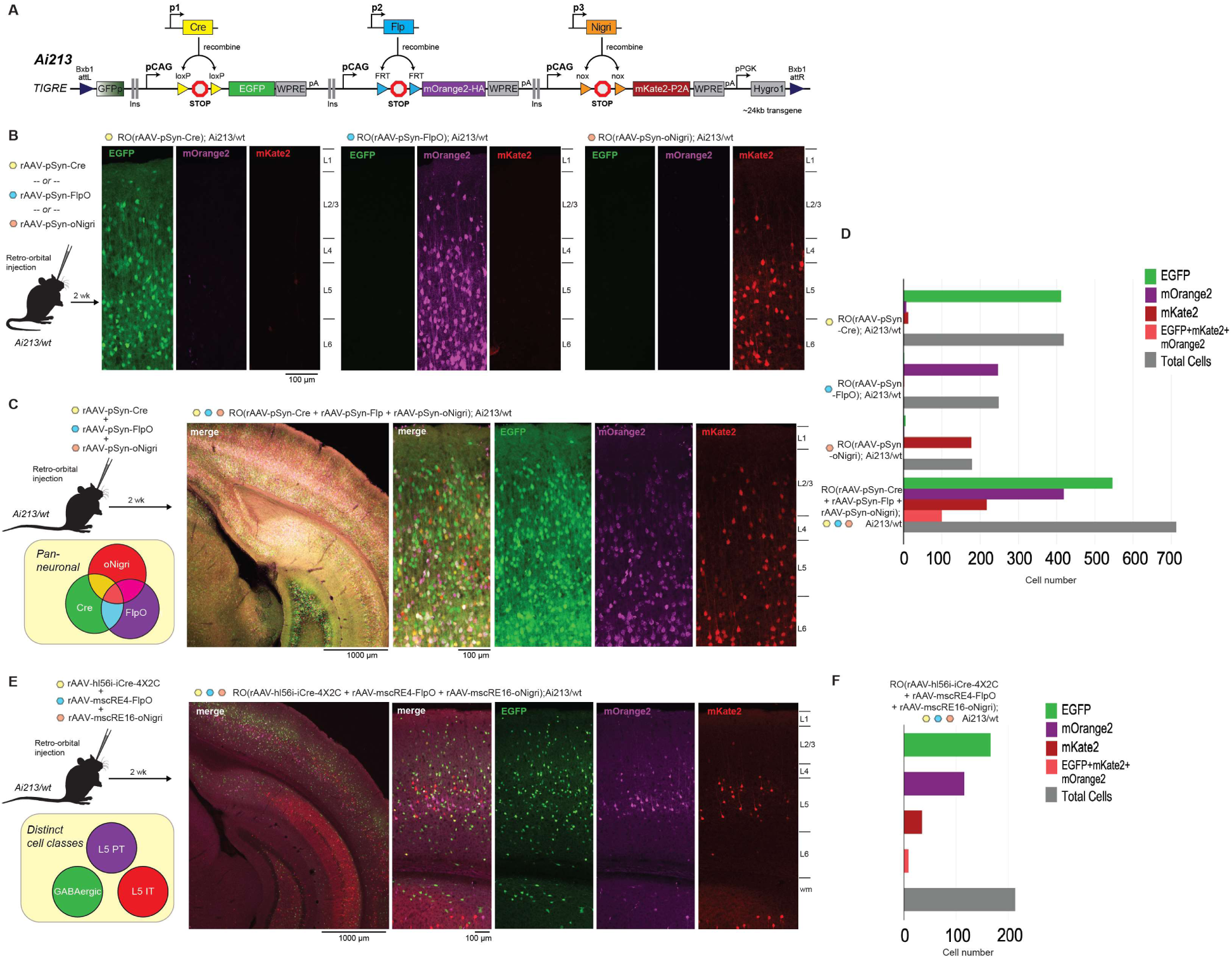
Combinatorial cell class labeling with a novel 3-color reporter line, *Ai213*. (**A**) Schematic diagrams illustrating the three-independent transgene expression units engineered into the *TIGRE* locus to create the *Ai213* reporter line. mOrange2 was tagged with the HA epitope and mKate2 was tagged with a P2A epitope to enable detection of each protein with antibodies in addition to direct fluorescence. (**B,C**) *Ai213* heterozygous mice were injected retro-orbitally with either pSyn-Cre (B, left panel), pSyn-FlpO (B, middle panel) or pSyn-oNigri (B, right panel) viruses or all three viruses in combination (C); final concentration of each virus was 1 × 10^11^ GCs. Native fluorescence was imaged by confocal microscopy in brain sections (focus on VISp) two weeks post-injection. Images were acquired using the same instrument settings; scale bars and layer boundaries are indicated. Injection of the separate synapsin promoter-driven viruses into Ai213 mice shows the expected recombinase dependence, with little cross-recombination at the non-cognate STOP cassettes. Coadministration of all three synapsin promoter-driven viruses into *Ai213* mice demonstrates labeling throughout the mouse cortex. The number of cells labeled with specified fluorophores from genotypes and viruses indicated on the left from images in (B) and (C). All counts were collected from at least one image per condition. (**E**) *Ai213* heterozygous mice were injected retro-orbitally with a mixture of the hl56i-iCre-4×2C (pan-GABAergic), mscRE4-FlpO (L5 PT), and mscRE16-oNigri (L5 IT) viruses each at a final concentration of 1X 10^11^ GCs. Native fluorescence was examined like in (B). Mostly mutually exclusive labeling of the expected cell types was observed in VISp. (**F**) Same as in (D) for data in (E).

To validate the *Ai213* reporter line, we initially evaluated the labelling achieved with retro-orbital delivery of pan-neuronal, synapsin promoter-driven rAAVs that expressed either Cre, FlpO, or a mouse codon-optimized Nigri (oNigri) recombinase (**Figure 7B**). Here we observed the robust expression of only one fluorophore as expected per recombinase in cortex and miniscule cross-recombination of Cre on FRT or nox sites, FlpO on loxP or nox sites, or oNigri on loxP or FRT sites (**Figure 7C**). When these three viruses were mixed together and co-injected into *Ai213* mice, we observed strong expression of all three fluorophores in the cortex, with most cells labeled by individual fluorophores (**Figure 7C-D**). Despite matching titers, many more EGFP+ cells were observed relative to mOrange2+ and mKate2+ overall (**Figure 7C-D**), which is likely a reflection of Cre being a more efficient recombinase compared to FlpO and oNigri as reported (Karimova et al., 2016). We reasoned that low fluorophore co-expression could be due to poor co-transduction efficiency of different viruses. To test if we can increase the labeling by improving viral transduction efficiency, we delivered the same combination of three viruses directly into the cerebral ventricle of *Ai213* neonates (**Figure S14**). This approach has been shown previously to yield widespread transduction of neurons throughout the mouse brain (Kim et al., 2014). As expected, we observed more cells labeled by each fluorophore in brain sections from ICV-injected *Ai213* mice, as well as many more double and triple labeled cells, showing that the three expression cassettes in *Ai213* can all be expressed from the TIGRE locus in the same cells *in vivo* (**Figure S14B-D**).

To determine if distinct cell types could be labelled simultaneously with *Ai213*, we retro-orbitally co-injected a trio of subclass-specific enhancer-driven recombinase viruses into an adult *Ai213* transgenic mouse (**Figure 7E**): rAAV-mscRE4-FlpO, which targets FlpO to L5 PT cells (magenta, mOrange2); and rAAV-mscRE16-oNigri, which targets L5 IT cells (red, mKate2); and rAAV-hi56i-iCre-4×2C, which targets iCre expression to GABAergic cell types (green, EGFP). The rAAV-hi56i-iCre-4×2C vector incorporates a micro RNA-binding element (mAGNET) that suppresses expression in excitatory cells (Sayeg et al., 2015), and is described in greater detail in **Figure S15**. Two weeks after injection, we observed mostly non-overlapping expression of the three fluorophores with expected layer specificity: EGFP broadly labeled GABAergic cells throughout the cortex, while mOrange2 and mKate2 were found primarily in L5 in mostly non-overlapping cell populations within VISp (**Figure 7F**). Collectively, these data demonstrate the broad utility of *Ai213* and show that multiplexed-enhancer-reporter experiments for labeling three distinct cell classes or subclasses in a single transgenic animal can be accomplished with ease.

### Labeling of L5 PT neurons in human *ex vivo* slices

To test if an enhancer element discovered in mouse can be functionally conserved across mammalian species, we tested whether the mscRE4 enhancer can label L5 PT neurons in human neocortex. In human neocortex, L5 PT neurons are relatively rare, comprising <1% of the neuronal population in middle temporal gyrus (MTG) (Hodge et al., 2020). To label this rare neuronal population we applied the mscRE4-FlpO virus together with a Flp-dependent EGFP reporter virus directly to human MTG *ex-vivo* slice cultures (**Figure 8A**). Labeled neurons were sparsely observed throughout L2-6 and thus labeling was not nearly as specific as observed in mouse VISp. However, sparse neurons that were fluorescently labeled in L5 with large somata consistent with human PT neurons were readily targeted for patch-clamp electrophysiology. Example biocytin fills demonstrated that these large EGFP+ labeled neurons were thick-tufted neurons (**Figure 8B**), similar to rodent PT neuron morphology although much larger, with apical dendrites extending ∼2 mm to reach the pial surface. Furthermore, the labeled neurons exhibited electrophysiological properties consistent with those of PT neurons in rodents, including a low R_N_ and a resonant frequency of ∼5 Hz. In contrast, non-labeled neurons possessed properties consistent with non-PT neurons in rodent (i.e. a higher R_N_ and a lower resonant frequency; **Figure 8C-F**). Finally, for a subset of experiments we extracted the nucleus through the recording pipette at the end of the electrical recording for RNA sequencing. Three out of 4 EGFP+ labeled neurons mapped to a putative PT transcriptomic neuron type and one out of 7 EGFP-labeled neurons mapped to a putative L6 IT cluster (**Figure 8G**; (Hodge et al., 2020)). The other neurons did not yield sufficiently high quality of RNA-seq data to enable high-confidence mapping. These results demonstrate the feasibility of applying the mscRE4 enhancer in an AAV vector to label and functionally characterize L5 PT neurons across species.

**Figure 8.**
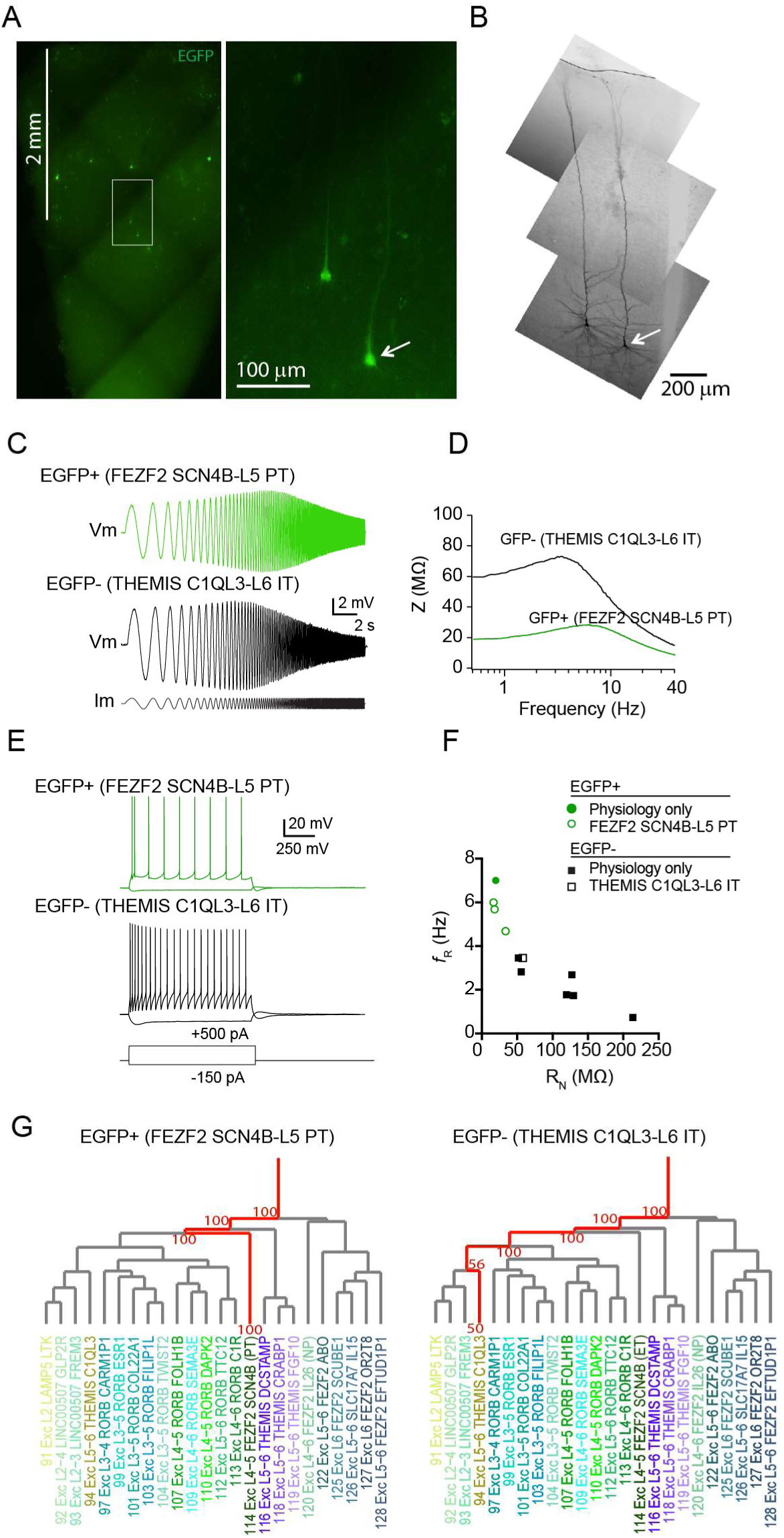
mscRE4-based virus labels L5 PT neurons in human middle temporal gyrus. (**A**) mscRE4-Cre enhancer virus in combination with a conditional reporter virus drives fluorescent protein expression in human MTG. EGFP+ neurons were targeted for patch-seq/standard patch-clamp experiments. (**B**) Biocytin fills of two double labeled neurons in human MTG. (**C**) Example voltage responses to a chirp stimulus for an EGFP+ neuron and a non-labeled (EGFP-) L5 pyramidal neuron. (**D**) Impedance amplitude profiles for the neurons in C. (**E**) Voltage response to a suprathreshold depolarizing current injection and hyperpolarzing current injection for the neurons in C. (**F**) Resonance frequency as a function of input resistance. (**G**) Representative EGFP-labeled neuron mapped to a putative ET transcriptomic cell type while the non-labeled neuron mapped to a L6 IT transcriptomic cell type.

## Discussion

All complex multicellular organisms perform an almost miraculous feat: transforming a single genome into a multitude of highly specialized cell types and tissues. This diversity of interpretation of a single, finite source – the genome – is enabled by developmentally regulated epigenetic programs that selectively reveal specific regions of the genome to enable specific gene expression (Klemm et al., 2019). Enhancers and other distal regulatory elements act as “adjectives” that modify the emphasis our cells place on genes to drive cell type-specific expression programs, thus regulating the construction of highly diverse and specialized tissues such as the brain (Preissl et al., 2018; Visel et al., 2013). Identifying and extracting these regulatory elements, through identification in specific tissues and by conservation (Dickel et al., 2018), allows us to reuse the cell’s own reading of regulatory elements to build genetic tools so that we can interrogate specific, epigenetically similar groups of cells repeatedly in different animals.

Enhancer-driven cell type selectivity has a rich history of use in *Drosophila*, where *cis*-regulatory modules have been used in the GAL4-UAS system to express transgenes in specific cell types (Pfeiffer et al., 2010). In *Drosophila*, these regulatory modules have been found using large screens of conserved regulatory modules (CRMs) located proximal to genes (Pfeiffer et al., 2008), or identified using enhancer traps. The latter strategy has also been employed in mice to generate hundreds of reporter lines, many with region- or type-restricted expression patterns (Kelsch et al., 2012; Shima et al., 2016). However, these methods require extensive screening of individual enhancer-driven expression patterns, which is resource-intensive. Extending this concept to prospective labeling of mammalian cell types using viral drivers can accelerate identification of cell type-specific enhancers, especially for cell types with no existing genetic tools.

We have demonstrated that a combination of scRNA-seq and scATAC-seq identifies functional enhancers that preferentially label predicted cell classes and subclasses. Previous methods for enhancer identification, such as STARR-seq (Arnold et al., 2013), CRE-seq (Shen et al., 2016), or EDGE (Blankvoort et al., 2018) rely on large screens of genomic elements identified in bulk tissues with retrospective identification of labeled cell types. In contrast, our methodology extends concepts previously used in the retina, where defined populations of cell types were used to identify putative regulatory elements for screening (Hartl et al., 2017; Jüttner et al., 2019; Kishi et al., 2019), and in *Drosophila*, where whole-organism scATAC-seq revealed clade-specific regulatory elements that were validated by transgenesis (Cusanovich et al., 2018). Utilizing single-cell measurements of chromatin accessibility in the mouse cortex allowed us to efficiently screen for enhancer-driven labeling of targeted cell subclasses. And while we generated many cell class-specific viral tools, the success rate largely depended on the cell class targeted, and at best was 78% for L5 PT cells. Methodological improvements in single-cell measurements of chromatin accessibility will continue to improve our ability to prospectively select regulatory elements and decrease the time and cost of screening (Gasperini et al., 2020). Recent developments of large-scale methods for paired measurements of scATAC-seq and scRNA-seq (e.g. sci-CAR (Cao and Jonathan S. Packer, 2017), scACAT-seq (Liu et al., 2019), and SNARE-seq (Chen et al., 2019)) as well as methods for linking distal enhancers to the promoters they regulate (e.g. single-cell Hi-C (Nagano et al., 2013) and scHi-C (Ramani et al., 2017) should enhance our understanding of gene regulation and inform enhancer selection.

The high-quality scATAC-seq dataset in this study provides a foundational resource for locating enhancer elements to target diverse cell types in the mouse cortex. Through the use of a single-cell-per-well method, we were able to enrich for rare cell types in cortex and validate cortical projection types using Retro-ATAC-seq. We created new tools to resolve cell subclasses in L5 of the mouse cortex, some with specificity comparable to available recombinase driver lines (Feng et al., 2000; Tasic et al., 2016; Tasic et al., 2018). These viral tools are immediately useful in conjunction with existing reporter and/or driver lines to augment and expand the existing genetic toolkit (Daigle et al., 2018; Gong et al., 2007; Madisen et al., 2015; Madisen et al., 2012; Madisen et al., 2010; Taniguchi et al., 2011). Concatenation of these natural regulatory elements resulted in higher reporter expression, as previously demonstrated for similar AAV tools targeting cell types in muscle (Salva et al., 2007).

### Current limitations of enhancer viruses

Several key limitations must be carefully considered in experimental designs using the tools presented here: variations in the extent and specificity of labeling due to virus titer, delivery method and animal age, as well as incompleteness of labeling. We have noticed marked variation in the specificity and completeness of labeling at different titers of retro-orbital virus injection (**Figure S11**), and decreased specificity at high multiplicity of infection in stereotaxic injection experiments (**Figure S10**), as well as perturbations in transcriptomic state in stereotaxic experiments (**Figure S13**). In addition, we have seen increased infectivity in younger animals (P28 vs P56, data not shown). It is therefore imperative that any researcher using these tools perform experiments to select the delivery method, titer, age of animals, and genetic strain (Huang et al., 2019) to provide the level of specificity required for their experimental needs.

In experiments with the *Ai213* conditional reporter line, the retro-orbital delivery of Synapsin promoter-driven viruses, which should label all neurons, resulted in incomplete labeling and notable differences in recombination efficiency between recombinases (**Figure 7C**). The latter was expected as Cre was reported to be a more efficient recombinase than FlpO and were reported to be more efficient than Nigri (Cre>Flp>Nigri). ICV delivery of pan-neuronal viruses to neonates dramatically improved the extent of labelling observed in the brain (**Figure S14**) and therefore may be an alternative approach to consider when higher transduction efficiency and more complete labeling a given cell class or type is needed. It is notable however, that neither viral delivery method resulted in a substantial number of cells labeled by two or three fluorophores even when same promoter (synapsin, in this case) was used to drive each of the three recombinases. This may be due to incomplete viral infection and may be improved by delivering more FlpO and Nigri viruses relative to Cre virus to compensate for the differences in recombinase efficiency.

### Combinatorial and cross-species studies

In some cases presented here, the viral tools have a potential to supersede germline transgenesis for labeling and perturbation of specific cell types. L5 PT cells can be labeled using other methods, including injection of retrograde tracing viruses into the thalamus, spinal cord, or pons to label subcortically projecting neurons (Economo et al., 2018), or through the use of driver lines that label L5 PT cells (Sorensen et al., 2015), though the latter may include off-target expression in other cell subclasses or types (e.g., L5 IT with Rbp4-Cre_KL100 (Tasic et al., 2016) or L6 cell types with. Thy1-EGFP-H (Porrero et al., 2010)). Our mscRE4-FlpO virus and the optimized 3xCore-mcsRE4-SYFP2 virus provide an alternative approach to highly specific labeling of L5 PT cells in the cortex, with the added flexibility provided by retro-orbital delivery of a virus. In addition, the mscRE4-FlpO virus labels L5 PT cells in human neocortical slices (**Figure 8**) and therefore may be portable for use across species as has been shown for other enhancer-based viral tools (Dimidschstein et al., 2016; Mich et al., 2019; Vormstein-Schneider, 2019).

To complement the quickly evolving landscape of viral drivers, we developed *Ai213*, the first triple-recombinase reporter contained in a single genomic locus. By using *Ai213*, we demonstrated that multiple enhancer viruses can be combined to label mutually exclusive cell classes *in vivo* without interfering with the function of one another. This approach provides an alternative to complex breeding strategies for the generation of triple or quadruple transgenic animals and enables the rapid screening of up to three enhancer viruses in a single animal. In addition, the use of Cre and Flp-dependent fluorophores in *Ai213* makes this line compatible with existing transgenic driver lines, further opening strategies that include both well-characterized recombinase drivers and new viral tools. The targeted enhancer-driven viral reagents and the new transgenic mouse line developed here open new frontiers for the combinatorial exploration of brain cell types in mouse and beyond.

## Acknowledgments

We could not have performed this study without the support of the Allen Institute Animal Care Team for mouse husbandry, the Allen Institute Transgenic Colony Management team for colony management, the Allen Institute Laboratory Animal Services team for preparation and delivery of experimental animals; Tamara Caspar, Kirsten Chrichton, Matthew Kroll, Josef Sulc, and Herman Tung for tissue processing; Nadiya Shapovalova, Daniel Hirschstein, and Susan Bort for FACS; Natalie Weed and Victoria Omstead for retro-orbital injections; Refugio Martinez and Ximena Opitz-Araya for cloning; and Darren Bertagnolli, Michael Tieu, Delissa McMillen, Thanh Pham, Christine Rimorin, Katelyn Ward, Alexandra Glandon, and Amy Torkelson for scRNA-seq processing. The authors also thank Andrew Hill and Darren Cusanovich for assistance in accessing and reusing data published in Cusanovich and Hill *et al.* (2018). We thank Advanced Cell Diagnostics for validation of the *Fam84b* probe and for early access to the RNAscope HiPlex assays.

## Author contributions

B.T., L.T.G., and T.L.D. designed the study; J.T.T., B.P.L, and J.K.M. provided viral genome constructs and cloning protocols; G.H.L. and M.W. cloned enhancers and generated viral constructs. T.L.D., L.S., and G.H.L., M.W. designed and generated the Ai213 transgenic mouse line. J.T.T. identified the mscRE4 enhancer core and performed cloning experiments for the concatemer and provided the hI56i-iCre-4×2C virus. A.S.-C., T.N., and L.T.G. performed scATAC-seq experiments. B.K. and J.T. performed electrophysiology experiments. S.Y., M.M., and T.Z. performed viral packaging and purification. T.N., T.K., and M.M. performed retrograde injections. M.W. and L.S. performed retro-orbital injections. T.N., and B.G. E.S. performed ICV and stereotaxic injections. M.W., J.B., E.M., L.S. and T.L.D. performed IHC experiments, imaging, cell counting and analysis. T.N. and E.G. performed mFISH validation experiments. N.D. managed tissue processing for RNA-seq experiments. K.S. managed RNA-seq experiments. R.L. and L.E. managed the transgenic line colonies. A.H.C. managed virus production. Z.Y. and L.T.G. performed RNA-seq analysis. L.T.G. and A.S.-C. performed scATAC-seq analysis. B.K. performed electrophysiology data analysis. S.M.S. provided project management. H.Z. and E.L. lead the Cell Types Program at the Allen Institute. L.T.G., T.L.D., and B.T. wrote the manuscript, with input from all coauthors.

## Data and materials availability

scATAC-seq and scRNA-seq data will be deposited to GEO. Software code used for data analysis and visualization is available from GitHub at https://github.com/AllenInstitute/graybuck2019analysis/. An R package for analysis of low-coverage accessibility and transcriptomics (lowcat) is available on GitHub at https://github.com/AllenInstitute/lowcat/, and an R package for generating figures based on the Allen Institute Common Coordinate Framework (cocoframer) is available at https://github.com/AllenInstitute/cocoframer/.

## Supplementary Material

## STAR Methods

### Mouse breeding and husbandry

Mice were housed under Institutional Care and Use Committee protocols 1508, 1802, and 1806 at the Allen Institute for Brain Science, with no more than five animals per cage, maintained on a 12 hr day/night cycle, with food and water provided *ad libitum*. Animals with anophthalmia or microphthalmia were excluded from experiments. Animals were maintained on a C57BL/6J genetic background.

### Retrograde labeling

We performed stereotaxic injection of CAV-Cre (gift of Miguel Chillon Rodrigues, Universitat Autònoma de Barcelona) (Hnasko et al., 2006) into brains of heterozygous or homozygous *Ai14* mice using stereotaxic coordinates obtained from the Paxinos adult mouse brain atlas (Paxinos and Franklin, 2013). Specific coordinates used for each injection are provided in **Table S3**. tdTomato-positive single cells were isolated from VISp by FACS. Example FACS gating is provided in **Figure S1**.

### Single cell ATAC

Single-cell suspensions of cortical neurons were generated as described previously (Gray et al., 2017), with the exception of the addition of trehalose to the dissociation and sorting medium for some samples, and the use of papain in place of pronase for some samples as shown in **Table S2**. Where trehalose was used, we made a 132 mM trehalose stock solution by mixing 12.49 g trehalose dihydrate (Sigma-Aldrich Cat#T9531) with 250 mL of water. ACSF + trehalose solutions were made by mixing 50 mL of trehalose stock solution with 450 mL artificial cerebrospinal fluid (ACSF). When used, ACSF + trehalose was used in place of all ACSF solutions. Where papain was used for dissociations, we added 4.17 mL ACSF (with or without trehalose) to a 125 U vial of Papain (Worthington Biochemical PDS Kit Cat#LK003176) for a final concentration of 30 U/mL. After microdissection, samples were digested in the 30 U/mL papain solution for 30 min at 32°C. We then sorted individual cells using FACS with gating of DAPI-negative and fluorophore-positive labeling (tdTomato, EGFP, or SYFP2) to select for live neuronal cells or DAPI-negative and fluorophore-negative labeling for live non-neuronal cells. Example FACS gating is provided in **Figure S1**.

For GM12878 scATAC, cells were obtained from Coriell Institute, and were grown in T25 culture flasks in RPMI 1640 Medium (Gibco, Thermo Fisher Cat#11875093) supplemented with 10% fetal bovine serum (FBS; Hyclone Cat#SH30070.03) and 1% Penicillin Streptomycin (Life Technologies Cat#15140-122). At ∼80% confluence, cells were transferred to a 15 mL conical tube, centrifuged, and washed with PBS containing 1% FBS. Cells were then resuspended in PBS with 1% FBS and 2 ng/mL DAPI (DAPI*2HCl, Life Technologies Cat#D1306) for FACS sorting of DAPI-negative live cells. Single cells were sorted into 200 μL 8-well strip tubes containing 1.5 μL tagmentation reaction mix (0.75 μL Tagment DNA Buffer (Illumina Cat# 15027866), 0.2 μL Nextera Tn5 Enzyme (Illumina TDE1, Cat# 15027865), 0.55 μL water). After collection, cells were briefly spun down in a bench-top centrifuge, then immediately tagmented at 37 °C for 30 minutes in a thermocycler. After tagmentation, we added 0.6 μL Proteinase K stop solution to each tube (5 mg/mL Proteinase K solution (Qiagen Cat#19131), 50 mM EDTA, 5 mM NaCl, 1.25% SDS) followed by incubation at 40°C for 30 minutes in a thermocycler. We then purified the tagmented DNA using Agencourt AMPure XP beads (Beckman Coulter Cat#A63881) at a ratio of 1.8:1 resuspended beads to reaction volume (3.8 μL added to 2.1 μL), with a final elution volume of 11 μL of Buffer EB (Qiagen Cat# 19086). Libraries were indexed and amplified by the addition of 15 uL 2X Kapa HiFi HotStart ReadyMix (Kapa Biosystems, Cat# KK2602) and 2 uL Nextera i5 and i7 indexes (Illumina Cat# FC-121-1012) to each tube, followed by incubation at 72°C for 3 minutes and PCR (95°C for 1 minute, 22 cycles of 98°C for 20 seconds, 65°C for 15 seconds, and 72°C for 15 seconds, then final extension at 72°C for 1 minute). After amplification, sample concentrations were measured using a Quant-iT PicoGreen assay (Thermo Fisher Cat#P7589) in duplicate. For each sample, the mean concentration was calculated by comparison to a standard curve, and the mean and standard deviation of concentrations was calculated for each batch of samples. Samples with a concentration greater than 2 standard deviations above the mean were not used for downstream steps, as these were found in early experiments to dominate sequencing runs. These cells have high, non-specific sequence diversity and low overlap with ENCODE peaks, suggesting that they have lost coherent chromatin structure. All other samples were pooled by combining 5 μL of each sample in a 1.5 mL tube. We then purified the combined library by adding Ampure XP beads in a 1.8:1 ratio, with final elution in 50 μL of Buffer EB (Qiagen Cat# 19086). The mixed library was then quantified using a BioAnalyzer High Sensitivity DNA kit (Agilent Cat# 5067-4626) according to manufacturer’s instructions.

### scATAC sequencing, alignment, and filtering

Mixed libraries, containing 60 to 96 samples, were sequenced on an Illumina MiSeq at a final quantity of 20-30 pmol in paired-end mode with 50 nt reads. After sequencing, raw FASTQ files were aligned to the GRCm38 (mm10) mouse genome using Bowtie v1.1.0 (Langmead et al., 2009)as described previously (Gray et al., 2017). After alignment, duplicate reads were removed using samtools rmdup (Li et al., 2009), which yielded only single copies of uniquely mapped paired reads in BAM format. For analysis, we removed samples with fewer than 10,000 paired-end fragments (20,000 reads) and with more than 10% of sequenced fragments longer than 250 bp. An additional filter was created using ENCODE whole cortex DNase-seq HotSpot peaks (sample ID ENCFF651EAU from experiment ID ENCSR00COF) (Yue et al., 2014). Samples with less than 25% of paired-end fragments that overlapped DNase-seq peaks were removed from downstream analysis. Cells passing these criteria both had sufficient number of unique reads for downstream analysis, and had high-quality chromatin accessibility profiles as assessed by fragment size analysis (**Figure S2**). As an additional QC check, we compared aggregate scATAC-seq data to bulk ATAC-seq data from matching Cre-driver lines, where available. We found that aggregate single-cell datasets matched well to previously published bulk datasets (**Figure S6**).

### Jaccard distance calculation, PCA and tSNE embedding, and density-based clustering

To compare scATAC-seq samples, we downsampled all cells to an equal number of uniquely aligned fragments (10,000 per sample), extended these fragments to a total length of 1kb centered on the middle of each fragment, then collapsed any overlapping fragments within each sample into regions based on the outer boundaries of overlapping fragments. We then counted the number of overlapping regions between every pair of samples, and divided by the total number of regions in both samples to obtain a Jaccard similarity score. These scores were converted to Jaccard distances (1 – Jaccard similarity), and the resulting matrix was used as input for PCA, followed by *t*-distributed stochastic neighbor embedding (*t*-SNE). After *t*-SNE, samples were clustered in the *t*-SNE space using the RPhenograph package (Levine et al., 2015) with k = 6 to obtain small groups of similar neighbors (Levine et al., 2015). Phenograph cluster assignments used for correlation with transcriptomic data are shown in **Table S6**.

### Correlation of scATAC-seq with scRNA-seq

Phenograph-defined neighborhoods were assigned to cell subclasses and clusters by comparison of accessibility near transcription start site (TSS) to median expression values of scRNA-seq clusters at the cell type level (e.g., L5 CF Chrna6) from mouse primary visual cortex (Tasic et al., 2018). To score each TSS, we retrieved TSS locations from the RefSeq Gene annotations provided by the UCSC Genome Browser database, and generated windows from TSS ± 20kb. We then counted the number of fragments for all samples within each cluster that overlapped these windows. For comparison, we selected differentially expressed marker genes from the Tasic *et al.* scRNA-seq dataset (Tasic et al., 2018) using the scrattch.hicat package for R. We then correlated the Phenograph cluster scores with the log-transformed median exon read count values for this set of marker genes for each scRNA-seq cluster from primary visual cortex, and assigned the transcriptomic cell type with the highest-scoring correlation. We found that this strategy of neighbor assignment and correlation allowed us to resolve cell types within the scATAC-seq data close to the resolution of the scRNA-seq data, as types that were split too far would be assigned to the same transcriptomic subclass or type by correlation.

### scATAC-seq grouping and peak calling

For downstream analysis, we grouped cell type assignments to the subclass level, except for highly distinct cell types (Lamp5 Lhx6, Sst Chodl, Pvalb Vipr2, L6 IT Car3, CR, and Meis2). Unique fragments for all cells within each of these subclass/distinct type groups were aggregated to BAM files for analysis. Aligned reads from single cell subclasses/clusters were used to create tag directories and peaks of chromatin accessibility were called using HOMER (Heinz et al., 2010) with settings “findPeaks -region -o auto”. The resulting peaks were converted to BED format.

### Population ATAC of Sst neurons

We performed population ATAC-seq of neurons from *Sst-IRES2-Cre/wt;Ai14/wt* mice as described previously (Gray et al., 2017). Briefly, cells from the visual cortex of an adult mouse were microdissected and FACS-isolated into 8-well strips as described above, but with 500 cells per well instead of single cells as for scATAC-seq. Cell membranes were lysed by addition of 25 5 µL Cell Lysis Buffer (10 mM Tris-HCl pH 7.4, 10 mM NaCl, 3 mM MgCl2, and 0.1% IGEPAL CA-630) to ∼5 µL of FACS-sorted cells, and nuclei were pelleted before resuspension in the same tagmentation buffer described above at a higher volume (25 µL). Tagmentation was carried out at 37°C for 1 hour, followed by addition of 5 µl of Cleanup Buffer (900 mM NaCl, 300 mM EDTA), 2 µl 5% SDS, and 2 µl Proteinase K and incubation at 40°C for 30 minutes, and cleanup with AMPure XP beads at a ratio of 1.8:1 beads to reaction volume. Samples were amplified using KAPA HotStart Ready Mix and 2 µl each of Nextera i5 and i7 primers (Illumina), quantified using a Bioanalyzer, and sequenced on an Illumina MiSeq in paired-end mode with 50 nt reads.

### Comparisons to bulk ATAC-seq data

For comparison to previously published studies, we used data from GEO accession GSE63137 for Camk2a, Pvalb, and Vip neuron populations, and GEO accession GSE87548 (Gray et al., 2017) for Cux2, Scnn1a-Tg3, Rbp4, Ntsr1, Gad2, mES, and genomic controls. For these comparisons, we also included population ATAC-seq of Sst neurons, described above. For each population, we merged reads from all replicates and downsampled each region to 6.4 million reads. We then called peaks using HOMER as described for aggregated scATAC-seq data, above. We used the BED-formatted peaks for scATAC-seq aggregates with or without bulk ATAC-seq datasets as input for comparisons using the DiffBind package for R as described previously (Gray et al., 2017). For all samples, bulk genomic DNA ATAC-seq was used as a background control from (Gray et al., 2017).

### Identification of mouse single-cell regulatory elements

We performed a targeted search for mouse single cell regulatory elements (mscREs) by performing pairwise differential expression analysis of scRNA-seq clusters from (Tasic et al., 2018) to identify uniquely expressed genes in L5 CF, L5 IT, and L6 IT subclasses as well as the L6 IT Car3 cell type across all glutamatergic subclasses. We then searched for unique peaks within 1 Mbp of each marker gene, and manually inspected these peaks for low or no accessibility in off-target cell types and for conservation (phastCons scores). If a region of high conservation overlapped the peak region, but the peak was not centered on the highly conserved region, we adjusted the peak selection to include neighboring highly conserved sequence. For cloning, we centered our primer search on 500 bp regions centered at the middle of the selected peak regions, and included up to 100 bp on either side for primer selection. Final region selections and PCR primers are provided in **Table S6**.

### Recombinant viral genome construction

Enhancers were cloned from C57Bl/6J mouse genomic DNA using enhancer-specific primers (**Table S6**) and Phusion high-fidelity polymerase (NEB Cat#M0530S). Individual enhancers were then inserted into an rAAV or scAAV backbone that contained a minimal beta-globin promoter or minimal CMV promoter, gene, a bovine growth hormone polyA (BGHpA), and a woodchuck post-transcriptional regulatory element (WPRE or WPRE3) using standard molecular cloning approaches. Viral genome construct details are available in **Table S7**. The 3xCore of the mscRE4 enhancer was created by custom gene synthesis and inserted into the rAAV backbone. For AAV vectors with the DLX enhancer, the DLX enhancer sequence was PCR amplified from human genomic DNA and cloned into the rAAV or scAAV backbone. To create rAAV-DLX-minBglobin-iCre-4×2C-WPRE-BGHpA, the 4×2C sequence (Sayeg et al., 2015) was generated by custom gene synthesis and inserted 3’ of the iCre cDNA sequence in rAAV-DLX-minBglobin-iCre-WPRE-BGHpA. All plasmid sequences were verified via Sanger sequencing and restriction digests were performed to confirm intact inverted terminal repeat (ITR) sites.

### Viral packaging and tittering

To generate purified rAAVs of the PHP.eb serotype, 105 μg of AAV viral genome plasmid, 190 μg of the pHelper plasmid (encodes adenoviral replication proteins; Agilent Cat#240071), and 105 μg of the pUCmini-iCAP-PHP.eB plasmid (encodes engineered PHP.eB capsid protein; Addgene Cat#103005) (Chan et al., 2017) were mixed with 5 mL of Opti-MEM I media with reduced serum and GlutaMAX (ThermoFisher Scientific Cat#51985034) and 1.1 mL of a solution of 1 mg/mL 25 kDa linear Polyethylenimine (PEI; Polysciences Cat#23966-1) in PBS at pH 4-5. This co-transfection mixture was incubated at room temperature for 10 minutes, then 0.61 mL of this co-transfection mixture was added to one 15-cm dish of ∼70-80%% confluent HEK293T cells (ATCC Number CRL-3216. A total of ten, 15 cm plates of cells were transfected for a small-scale packaging run. 24 hours post-transfection, cell medium was replaced with DMEM containing high glucose, L-glutamine and sodium pyruvate (ThermoFisher Scientific Cat# 11995073) and supplemented with 4% fetal bovine serum (FBS;Hyclone Cat#SH30070.03) and 1% Antibiotic-Antimycotic solution (Life Technologies Pen Strep Cat#15140-122). Cells were collected 72 hours post-transfection by manually scraping them from the culture dish into 5 mL of culture medium and were then pelleted by centrifugation at 1500 rpm at 4°C for 15 minutes. The cell pellet was resuspended in a buffer containing 150 mM NaCl, 10 mM Tris, and 10 mM MgCl2, pH 7.6, and was frozen in dry ice. Samples were thawed quickly in a 37°C water bath, then passed through a syringe with a 21-23 gauge needle 5 times, followed by 3 more freeze/thaw cycles, and a 30 minute incubation with 50 U/ml Benzonase nuclease (Sigma-Aldrich Cat#E8263) at 37°C to degrade DNA and RNA not contained within the viral particles. The suspension was then centrifuged at 3,000 × g to remove cellular debris and the virus-containing supernatant was purified using a layered iodixanol step gradient (15%, 25%, 40%, and 60%) by centrifugation at 58,000 rpm in a Beckman 70Ti rotor for 90 minutes at 18°C. Following ultracentrifugation, the virus-containing layer at the 40-60% interface was collected and concentrated using an Amicon Ultra-15 centrifugal filter unit (100 kDa cutoff; EMD Millipore Cat#UFC910024) and by centrifugation at 3,000 rpm at 4°C. The concentrated virus was diluted in PBS containing 5% glycerol and 35 mM NaCl before storage at −80°C. Crude lysate preps of rAAVs of the PHP.eb serotype were generated using a late harvest protocol similar to that found in (Jüttner et al., 2019). In brief, the same plasmids specified above were used at a ratio of 1:1:2 viral genome plasmid:serotype plasmid:helper plasmid to transfect one, 15 cm dish of HEK293T cells at ∼70-80% conf. 24 hours post-transfection, the cell culture medium was changed from DMEM containing sodium pyruvate and supplemented with 10% FBS to DMEM containing sodium pyruvate and supplemented with 1% FBS to serum starve the cells. 120 hours post-transfection, the media and cells were harvested, subjected to three freeze/thaw rounds, incubated with Benzonase nuclease, and concentrated to ∼150 µl using the Amicon Ultra-15 centrifugal filter unit. Virus titers were measured using qPCR or ddPCR. For qPCR, a primer pair that recognizes a region of 117 bp in the AAV2 ITRs (Forward: 5’-GGAACCCCTAGTGATGGAGTT-3’; Reverse: 5’-CGGCCTCAGTGAGCGA-3’) was utilized and reactions were performed using QuantiTect SYBR Green PCR Master Mix (Qiagen Cat#204145) and 500 nM of each primer. A positive control AAV with known titer and newly produced viruses with unknown titers were treated with DNase I (NEB Cat#M0303S) to degrade any plasmid DNA and serial dilutions (1/10, 1/100, 1/500, 1/2500, 1/12500, and 1/62500) were made and loaded onto the same qPCR plate. A standard curve of virus particle concentrations vs. quantitation cycle (C_q_) values was generated from the positive control virus and the titers of the new viruses were calculated based on this standard curve. To measure virus titers by ddPCR, an instrument manufactured by BIO-RAD and the AAV2 ITR primer pair specific above and a FAM-labeled probe (5’ 6-FAM/CGCGCAGAG/ZEN/AGGGAGTGG/3’ IABkFQ) was utilized. Serial dilutions of AAV samples (2.50E-05, 2.50E-06, 2.50E-07, and 2.50E-08) were used for measurement to fit the dynamic linear range of the ddPCR assay. ddPCR reaction assembly, droplet generation, PCR amplification of the droplets, plate scanning and data analysis were conducted according to the manufacturer’s instructions. Virus concentration was calculated for each diluted sample within the dynamic linear range of ddPCR, and AAV titer was reported as the mean of the calculated concentrations.

### Retro-orbital injections

We used 21 day-old or older C57BL/6J, *Ai14, Ai65F, Ai63, or Ai213* mice (Daigle et al., 2018; Madisen et al., 2015; Madisen et al., 2010). Mice were briefly anesthetized using isoflurane, and 1×10^10^ to 1×10^11^ viral genome copies (GC) were delivered into the retro-orbital sinus in a volume of 50 µL or less. This approach has been utilized previously to deliver AAVs across the blood-brain barrier and into the murine brain with high efficiency (Chan et al., 2017). For delivery of multiple viruses into one animal, the rAAVs were mixed beforehand and then delivered into the retro-orbital sinus in a total volume of 50 µL or less. Animals recovered the same day due to the minimally invasive nature of the procedure and were euthanized 1-3 weeks post-infection for analysis. For injection details, see **Table S3**.

### Stereotaxic and intracerebroventricular (ICV) injections

Purified PHP.eB serotyped rAAVs were produced as described above for mscRE4-SYFP2 (titer = 1.34 × 10^13^ GC/ml), mscRE4-EGFP (1.64 × 10^14^ GC/ml), mscRE16-EGFP (1.94 × 10^13^ GC/ml), Syn-Cre (2.0 × 10^13^ GC/ml), Syn-FlpO (3.0 × 10^13^ GC/ml), or Syn-oNigri (2.4 × 10^13^ GC/ml). For stereotaxic injections, each virus was delivered bilaterally at 250 nl, 50 nl, or 25 nl volumes into the primary visual cortex (VISp; coordinates: A/P: −3.8, ML: −2.5, DV: 0.6) of C57BL/6J mice using a pressure injection system (Nanoject II, Drummond Scientific Company, Cat# 3-000-204). To mark the injection site, a DJ serotyped rAAV expressing dTomato from the EF1α promoter was co-injected at a dilution of 1:10 with the enhancer-driven fluorophore virus. For ICV injections, the three rAAVs (Syn-Cre, Syn-FlpO, and Syn-oNigri) were mixed together to yield a final concentration of 2.0 × 10^10^ GC of each virus and injected into the right cerebral ventricle of *Ai213* heterozygous mice at P3. All mice were sacrificed 1-3 weeks post-injection; see **Table S3** for a list of donors and injection details. To evaluate the extent of viral labeling in the brain, mice were transcardially perfused with PBS followed by 4% paraformaldehyde (PFA) and the dissected brains were subsequently post-fixed in 4% PFA overnight at 4°C followed by cryoprotection for 1-2 days in a 30% sucrose solution. 50 µm sections were prepared using a freezing microtome (Leica SM2000R) and epifluorescence or confocal images were acquired from mounted sections using a Nikon Eclipse TI epi-fluorescence microscope or a Fluoview FV3000 series confocal laser scanning microscope. For scRNA-seq, virally infected, reporter-expressing cells were processed and analyzed as described below.

### Immunohistochemistry

Mice were anesthetized with isoflurane and transcardially perfused with 0.1 M phosphate buffered saline (PBS) followed by 4% PFA. Brains were removed, post-fixed in 4% PFA overnight, followed by an additional incubation for 2-3 days in 30% sucrose. Coronal sections (50 µm) were cut using a freezing microtome (Leica SM2000R) and native fluorescence or antibody-enhanced fluorescence was analyzed in mounted sections. To enhance the EGFP fluorescence, a rabbit anti-GFP antibody was used to stain free floating brain sections. Briefly, sections were rinsed three times in PBS, blocked for 1 hour in PBS containing 5% goat serum (Sigma-Aldrich Cat#G9023-10ML), 2% bovine serum albumin (BSA; Sigma-Aldrich Cat#A9418-50G) and 0.2% Triton X-100 (Sigma-Aldrich Cat#X100-500ML), and incubated overnight at 4°C in the anti-GFP primary antibody (1:3000; Abcam Cat#ab6556). The following day, sections were washed three times in PBS and incubated in blocking solution containing an Alexa 488 conjugated secondary antibody (1:1500; Invitrogen Cat#A-11034), washed in PBS, and mounted in Vectashield containing DAPI (Vector Labs Cat#H-1500). Epifluorescence images of native or antibody-enhanced fluorescence were acquired on a Nikon Eclipse Ti microscope.

### Single-cell RNA sequencing (scRNA-seq) and cell type mapping

scRNA-seq was performed using the SMART-Seq v4 kit (Takara Cat#634894) as described previously (Tasic et al., 2018). In brief, single cells were sorted into 8-well strips containing SMART-Seq lysis buffer with RNase inhibitor (0.17 U/uL; Takara Cat#ST0764), and were immediately frozen on dry ice for storage at −80°C. SMART-Seq reagents were used for reverse transcription and cDNA amplification. Samples were tagmented and indexed using a NexteraXT DNA Library Preparation kit (Illumina Cat#FC-131-1096) with NexteraXT Index Kit V2 Set A (Illumina Cat#FC-131-2001) according to manufacturer’s instructions except for decreases in volumes of all reagents, including cDNA, to 0.4x recommended volume. Full documentation for the scRNA-seq procedure is available in the ‘Documentation’ section of the Allen Institute data portal at http://celltypes.brain-map.org/. Samples were sequenced on an Illumina HiSeq 2500 as 50 bp paired-end reads. Reads were aligned to GRCm38 (mm10) using STAR v2.5.3 (Dobin et al., 2013) with the parameter “twopassMode,” and exonic read counts were quantified using the GenomicRanges package for R as described in Tasic *et al.* (2018) (Tasic et al., 2018). To determine the corresponding cell type for each scRNA-seq dataset, we utilized the scrattch.hicat package for R (Tasic et al., 2018). We selected marker genes that distinguished each cluster, then used this panel of genes in a bootstrapped centroid classifier which performed 100 rounds of correlation using 80% of the marker panel selected at random in each round. For plotting, we retained only cells that were assigned to the same cluster in ≥ 80 of 100 rounds. Mapping results and scRNA-seq sample metadata, including the most-frequently assigned cell type and the fraction of times each cell was assigned to that type, are included in **Table S8**.

### Differential gene expression analysis

To identify changes in gene expression induced by our viral genetic tools, we performed pairwise differential gene expression tests between virally labeled cells and cells labeled by transgenic mouse driver and reporter lines. For each viral scRNA-seq experiment, we selected all cell types to which at least 10 cells were reliably mapped (above), then used the DE_genes_pw() function from the scrattch.hicat package (Tasic et al., 2018) to perform differential gene expression analysis. This function utilized the limma package for R (Ritchie et al., 2015) to perform differential expression tests, along with some additional processing.

### Comparisons to previous scATAC-seq studies

For comparisons to GM12878 datasets, raw data from Cusanovich *et al.* (2015) (Cusanovich et al., 2015) was downloaded from GEO accession GSE67446, Buenrostro *et al.* (2015) (Buenrostro et al., 2015) from GEO accession GSE65360, and Pliner *et al.* (2018) (Pliner et al., 2018) from GEO accession GSE109828. Processed 10x Genomics data was retrieved from the 10x Genomics website for the experiment “5k 1:1 mixture of fresh frozen human (GM12878) and mouse (A20) cells.” Buenrostro, Cusanovich, Pliner, and our own GM12878 samples were aligned to the hg38 human genome using the same bowtie pipeline described above for mouse samples to obtain per-cell fragment locations. 10x Genomics samples were analyzed using fragment locations provided by 10x Genomics aligned to hg19. For comparison to TSS regions, we used the RefSeq Genes tables provided by the UCSC Genome Browser database for hg19 (for 10x data) and for hg38 (for other datasets). To compare to ENCODE peaks, we used ENCODE GM12878 DNA-seq HotSpot results from ENCODE experiment ID ENCSR000EJD aligned to hg19 (ENCODE file ID ENCFF206HYT) or hg38 (ENCODE file ID ENCFF773SCF).

### Electrophysiology

#### Brain slice preparation

Human and mouse brain slices were prepared using the NMDG protective recovery method (Ting et al., 2014; Ting et al., 2018). Human neurosurgical specimens were obtained from a 45-year-old female patient that underwent temporal cortex resection for the treatment of drug resistant temporal lobe. Mice were deeply anesthetized by intraperitoneal administration of Advertin (20 mg/kg) and were perfused through the heart with an artificial cerebral spinal (aCSF) solution containing (in mM): 92 NMDG, 2.5 KCl, 1.25 NaH_2_PO_4_, 30 NaHCO_3_, 20 HEPES, 25 glucose, 2 thiourea, 5 Na-ascorbate, 3 Na-pyruvate, .5 CaCl_2_. 4H_2_O and 10 MgSO_4_ 7H_2_O. Upon surgical resection, human neurosurgical tissue was immediately placed in NMDG aCSF and transported from the hospital to the Institute. Slices (300 μm) were sectioned on a Compresstome VF-200 (Precisionary Instruments) using a zirconium ceramic blade (EF-INZ10, Cadence). Human brain slices were prepared under sterile conditions in a biosafety hood. Mouse brains were sectioned coronally, and human tissue was sectioned such that the angle of slicing was perpendicular to the pial surface. After sectioning, slices were transferred to a warmed (32-34°C) recovery chamber filled with NMDG aCSF under constant carbogenation. After 12 minutes, slices were transferred to a holding chamber containing an aCSF made of (in mM) 92 NaCl, 2.5 KCl, 1.25 NaH_2_PO_4_, 30 NaHCO_3_, 20 HEPES, 25 glucose, 2 thiourea, 5 Na-ascorbate, 3 Na-pyruvate,128 CaCl_2_·4H_2_O and 2 MgSO_4_·7H_2_O continuously bubbled with 95/5 O_2_/CO_2. Mouse slices were held in this solution_ for use in acute recordings whereas human slices were transferred to a 6-well plate for long-term culture and viral transduction.

#### Human slice culture and viral transduction

Human brain slices were placed on membrane inserts and wells were filled with culture medium consisting of 8.4 g/L MEM Eagle medium, 20% heat-inactivated horse serum, 30 mM HEPES, 13 mM D-glucose, 15 mM NaHCO_3_, 1 mM ascorbic acid, 2 mM MgSO_4_·7H_2_O, 1 mM CaCl_2_.4H_2_O, 0.5 mM GlutaMAX-I, and 1 mg/L insulin (Ting et al., 2018). The slice culture medium was carefully adjusted to pH 7.2-7.3, osmolality of 300-310 mOsmoles/Kg by addition of pure H_2_O, sterile-filtered and stored at 4°C for up to two weeks. Culture plates were placed in a humidified 5% CO_2_ incubator at 35°C and the slice culture medium was replaced every 2-3 days until end point analysis. 1-3 hours after brain slices were plated on cell culture inserts, brain slices were infected by direct application of concentrated AAV viral particles over the slice surface (Ting et al., 2018).

#### Patch clamp physiology and analysis

For patch clamp recordings, slices were placed in a submerged, heated (32-34°C) recording chamber that was continually perfused with aCSF under constant carbogenation containing (in mM): 119 NaCl, 2.5 KCl, 1.25 NaH_2_PO_4_, 24 NaHCO_3_, 12.5 glucose, 2 CaCl_2_·4H_2_O and 2 MgSO_4_·7H_2_O (pH 7.3-7.4). Neurons were viewed with an Olympus BX51WI microscope and infrared differential contrast optics and a 40x water immersion objective. Patch pipettes (3-6 MΩ) were pulled from borosilicate glass using a horizontal pipette puller (P1000, Sutter Instruments). EGFP+ and/or SYFP+ neurons were identified using appropriate excitation/emission filter sets. The pipette solution for mouse experiments consisted of (in mM): 130 K-gluconate, 10 HEPES, 0.3 EGTA, 4 Mg-ATP, 0.3 Na2-GTP and 2 MgCl2 and 0.5% biocytin, pH 7.3. The pipette solution for human experiments was modified for patch-seq analysis and consisted of: 110 K-gluconate, 10.0 HEPES, 0.2 EGTA, 4 KCl, 0.3 Na2-GTP, 10 phosphocreatine disodium salt hydrate, 1 Mg-ATP, 20 µg/ml glycogen, 0.5U/µL RNAse inhibitor (Takara, 2313A) and 0.5% biocytin (Sigma B4261), pH 7.3. Electrical signals were acquired using a Multiclamp 700B amplifier and PClamp 10 data acquisition software (Molecular Devices). Signals were digitized (Axon Digidata 1550B) at 10-50 kHz and filtered at 2-10 kHz. Pipette capacitance was compensated and the bridge balanced throughout whole-cell current clamp recordings. Access resistance was 8-25 MΩ).

Data were analyzed using custom scripts written in Igor Pro (Wavemetrics). All measurements were made at resting membrane potential. Input resistance (R_N_) was calculated from the linear portion of the voltage-current relationship generated in response to a series of 1s current injections. The maximum and steady state voltage deflections were used to determine the maximum and steady state of R_N_, respectively. Voltage sag was fined as the ratio of maximum to steady-state R_N_. Resonance frequency (*f*_R_) was determined from the voltage response to a constant amplitude sinusoidal current injection that either linearly increased from 1-15 Hz over 15 s or increased logarithmically from .2-40 Hz over 20 s. Impedance amplitude profiles were constructed from the ratio of the fast Fourier transform of the voltage response to the fast Fourier transform of the current injection. *f*_R_ corresponded to the frequency at which maximum impedance was measured. While the majority of neurons we included in this study were located in primary visual cortex (n=10 YFP+, 10 YFP-), we also made recordings from motor cortex (n=1 YFP+) and primary somatosensory cortex (n=4 YFP). For illustrative purposes, we also compared the properties of YFP+ and YFP-neurons to 32-L5 pyramidal neurons located in somatosensory cortex from an uninfected mouse. To classify these neurons as IT-like or PT-like, we used Ward’s method of clustering. I_h_-related membrane properties are known to differentiate IT and PT neurons across many brain regions (Baker et al., 2018). As such, features included in clustering were restricted to the I_h-_ related membrane properties - sag ratio, R_N_ and *f*_R_.

#### Processing of patch-seq samples

For experiments in human slice cultures, the nucleus was extracted into the recording pipette at the end of the whole cell recording for RNA-sequencing. Prior to data collection, all surfaces were thoroughly cleaned with RNase Zap. The contents of the pipette were expelled into a PCR tube containing lysis buffer. cDNA libraries were produced using the SMART-Seq v4 Ultra Low Input RNA kit for Sequencing according to the manufacturer’s instructions. These data were then used to map each cell to a reference cell type in a previously published transcriptomic cell type taxonomy (Hodge et al., 2020) using the same tree-based mapping approach used for mapping single cell RNA sequencing samples described above.

### RNAscope

We retro-orbitally injected mscRE4-FlpO or mscRE16-FlpO rAAVs into brains for *Ai65F* mice, as well as mscRE4-SYFP2 AAV into brains of wildtype mice. Mice were sacrificed two weeks post-injection. Fresh brains were dissected and immediately embedded in optimum cutting temperature compound (OCT; TissueTek Cat#4583). The OCT blocks were stored at −80°C until they were sectioned. Twenty μm coronal sections were cut using a cryostat and collected on SuperFrost slides (ThermoFisher Scientific Cat#J3800AMNZ). RNA fluorescent in situ hybridization (FISH) with RNAscope HiPlex Assays (Advanced Cell Diagnostics Cat#324100) was performed according to the manufacturer’s instructions. All the probes were hybridized and amplified together and the detection was achieved iteratively in groups of up to three targets per round. Probes against *Fam84b* (ACD #500991-T1) were used to label L5 PT cells and probes against *Rorb* (ACD #444271-T3) were used to label L5 IT cells. *Rorb* also labels L4 cells, so probes against *Scnn1a* (ACD #441391-T5) and *Hsd11b1* (ACD #496231-T7) were used to delineate the layer 4 and 5 boundaries for analysis. The SYFP2 protein from the virus and tdTomato protein from the *Ai65F* conditional reporter degrade in fresh frozen sections, making native fluorescence in the tissue undetectable. Therefore, probes against the *SYFP2* mRNA (ACD #590291-T1) *tdTomato* mRNA (ACD #317041-T2) were used instead. Mounted sections were imaged using a 40x objective on a Leica SP8 confocal microscope and maximum intensity projections of z-stacks (1-µm intervals, for the middle 6 stacks) were created from each round of imaging. Nuclei were labeled by DAPI before prior to imaging and nuclear signal was used for registration across experimental rounds using the HiPlex Registration software (Advanced Cell Diagnostics). CellProfiler (http://www.cellprofiler.org) (Lamprecht et al., 2007) was used to segment DAPI stained nuclei and to identify spots from the FISH signal. To cover the spatial area occupied by mRNA of a cell, segmented DAPI borders were expanded by 25 pixels or until touching an adjacent border. Identified mRNA spots were related to a cell using the expanded nuclei border as the boundary. The number of detected mRNA spots per gene per cell, centroid coordinates of each segmented nucleus, and layer assignments are used as input into R for plotting and quantification. Thresholds above background labeling were manually determined for each probe and tissue section and are provided in **Table S9**.

### Whole-brain two photon imaging and analysis

TissueCyte images from virally infected brains were collected, registered, and segmented as described previously (Oh et al., 2014). After registration to the Allen Brain Atlas Common Coordinate Framework (CCFv3), 3D arrays of signal binned to 25 µm voxels were analyzed in R by subtraction of background and averaging the signal in the finest structures in the Allen Brain Atlas structural ontology. To propagate signals from fine to coarse structure in the ontology, we performed hierarchical calculations that assigned the maximum value of child nodes in the ontology to each parent from the bottom (finest) to the top (coarsest) levels of the ontology. Functions to assist this analysis are provided in the cocoframer package for R. We then filtered the ontology to select broad structures, and used the taxa and metacodeR packages for R (Foster, 2016) to display the resulting ontological relationships and structure scores.

### Data analysis and visualization software

Analysis and visualization of scATAC-seq and transcriptomic datasets was performed using R v.3.5.0 and greater in the Rstudio IDE (Integrated Development Environment for R) or using the Rstudio Server Open Source Edition as well as the following packages: for general data analysis and manipulation, data.table (Dowle, 2019), dplyr (Wickham, 2018), Matrix (Bates, 2018), matrixStats (Bengtsson, 2018), purr (Henry, 2019), and reshape2 (Wickham, 2007); for analysis of genomic data, GenomicAlignments (Lawrence et al., 2013), GenomicRanges (Lawrence et al., 2013), and rtracklayer (Lawrence et al., 2009); for plotting and visualization, cowplot (Wilke, 2018), ggbeeswarm (Clark, 2017), ggExtra (Attali, 2018), ggplot (Wichkham, 2016), and rgl (Adler, 2018); for clustering and dimensionality reduction, Rphenograph (Chen, 2015) and Rtsne (Krijthe, 2015); for analysis of transcriptomic datasets: scrattch.hicat and scrattch.io (Tasic et al., 2018); for taxonomic analysis and visualization, metacodeR (Foster, 2016) and taxa (Zachary, 2018); and plater (Hughes, 2016) for management of plate-based experimental results and metadata.

### Generation of the *Ai213* transgenic line

To target multiple transgene expression units into the *TIGRE* locus (Zeng et al., 2008) we employed a recombinase-mediated cassette exchange (RMCE) strategy similar to that previously described (Madisen et al., 2015), but instead of using Flp recombinase for targeting, Bxb1 integrase (Zhu et al., 2014) was used to “free-up” Flp for transgene expression control. A new landing pad mouse embryonic stem (ES) cell line was generated by taking the 129S6B6F1 cell line, G4 (George et al., 2007), and engineering it to contain the components from 5’ to 3’ **Bxb1** *Att<u>P</u>* **-PhiC31 AttB-PGK promoter-gb2 promoter-Neomycin gene-PGK polyA-Bxb1 AttP-splice acceptor-3’ partial hygromycin gene-SV40 polyA-PhiC31** *Att<u>P</u>* within the *TIGRE* genomic region. Southern blot, qPCR and junctional PCR analyses were performed on genomic DNA (gDNA) samples from modified ES cell clones to confirm proper targeting, copy number, and orientation of the components within the *TIGRE* locus. A Bxb1-compatible targeting vector with three independent and conditional expression units was then generated by standard molecular cloning techniques. The vector contained the following components from 5’ to 3’: gb2 promoter-Neo gene-Bxb1 AttB-partial GFP-2X HS4 Insulators-CAG promoter-LoxP-stop-LoxP-EGFP-WPRE-BGH polyA-2X HS4 Insulators-CAG promoter-FRT-stop-FRT-mOrange2-HA-WPRE-BGH polyA-PhiC31 AttB-WPRE-BGH polyA-2X HS4 Insulators-CAG-nox-stop-nox-mKate2-P2A-WPRE-PGK polyA-PhiC31 AttB-PGK promoter-5’ hygromycin gene-splice donor-Bxb1 AttB. The sequence and integrity of the targeting vector was confirmed by Sanger sequencing, restriction digests and *in vitro* testing performed in HEK293 cells. The targeting vector (30 µg of DNA) was then co-electroporated with a plasmid containing a mouse codon optimized Bxb1 gene under the control of the cytomegalovirus (CMV) promoter (100 µg of DNA) into the Bxb1-landing pad ES cell line and following hygromycin drug selection at 100-150 µg/ml for 5 days, monoclonal populations of cells were hand-picked and expanded. gDNA was prepared from the modified ES cell clones using a kit (Zymo Research Cat#D4071) and it was screened by qPCR and junctional PCR assays to confirm proper targeting into the *TIGRE* locus. Correctly targeted clones were injected into fertilized blastocysts at the University of Washington Transgenic Research Program (TRP) core to generate high percentage chimeras and then the chimeras were imported to the Institute, bred to C57BL/6J mice to produce F1 heterozygous reporter mice, and subsequently maintained in a C57BL/6J congenic background.

**Fig. S1.**
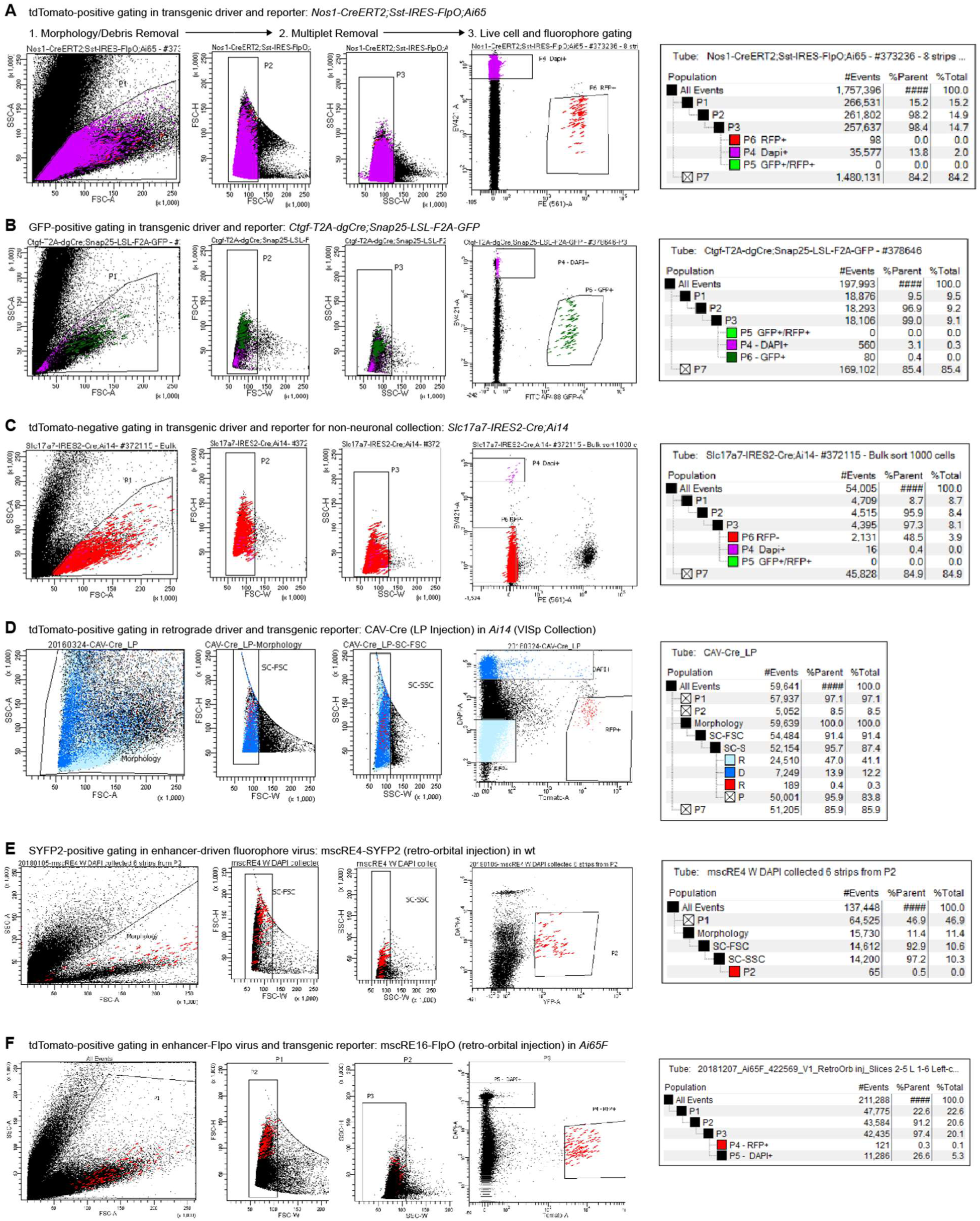
FACS Gating examples. All FACS collection followed a similar gating strategy: Morphology and debris exclusion using FSC-A and SSC-A; Exclusion of multiplets using FSC-W × FSC-H and SSC-W × SSC-H gating; and selection of live cells with or without fluorescent labels using DAPI exclusion and fluorophore signals. (**A**) Example gating for tdTomato+ cells from a transgenic driver and reporter. (**B**) Example gating for GFP+ cells from a transgenic driver and reporter. (**C**) Example gating for tdTomato-negative cells in a transgenic driver and reporter line for the collection of non-neuronal cells. (**D**) Example gating for tdTomato+ cells labeled by a retrograde driver in a transgenic reporter mouse line. (**E**) Example gating for direct SYFP2-labeling of cells from injection of mscRE4-SYFP2. (**F**) Example gating for tdTomato+ cells labeled by a FlpO virus injected into a transgenic reporter mouse line.

**Fig. S2.**
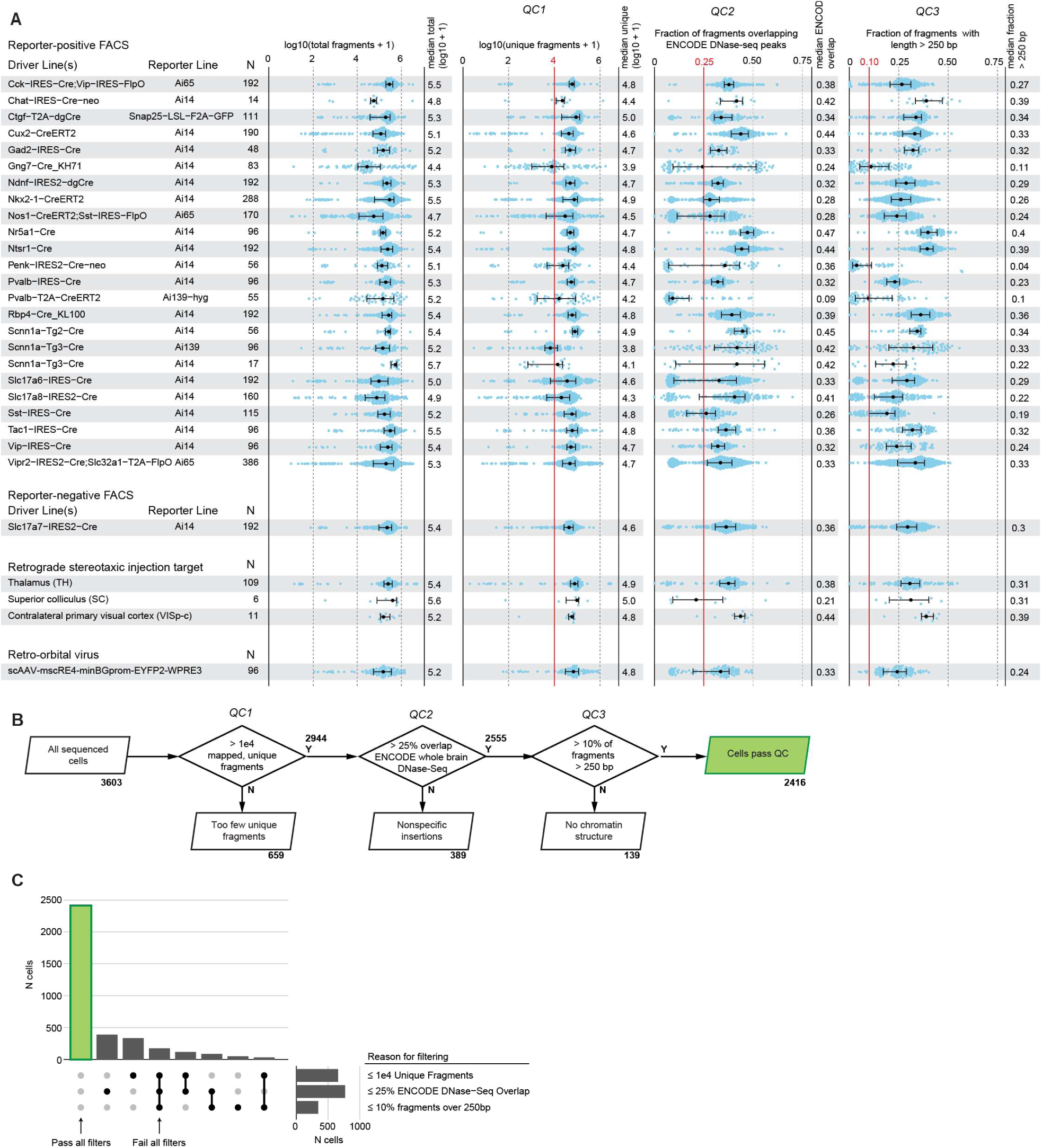
Cell sources and QC Statistics. (**A**) Fragment number and quality for each cell source in the scATAC-seq dataset. See **Table S1** for mouse line details. N, number of cells collected. ‘Total fragments’ is the count of all sequenced fragments for each cell. ‘Unique fragments’ is the number of uniquely mapped fragments, and was used for the first QC cutoff of (> 10,000 unique fragments; QC1). Fraction of fragments overlapping ENCODE DNase-seq peaks was computed for uniquely mapped fragments, and was used for the second QC cutoff (> 0.25; QC2). Fraction of fragments with length > 250 bp was computed for unique fragments, and was used for the third QC cutoff of (> 0.1; QC3). (**B**) Flow chart showing how many samples were sequentially filtered by these 3 QC criteria. (**C**) Set barplot showing how many cells were flagged with each combination of QC criteria.

**Fig. S3.**
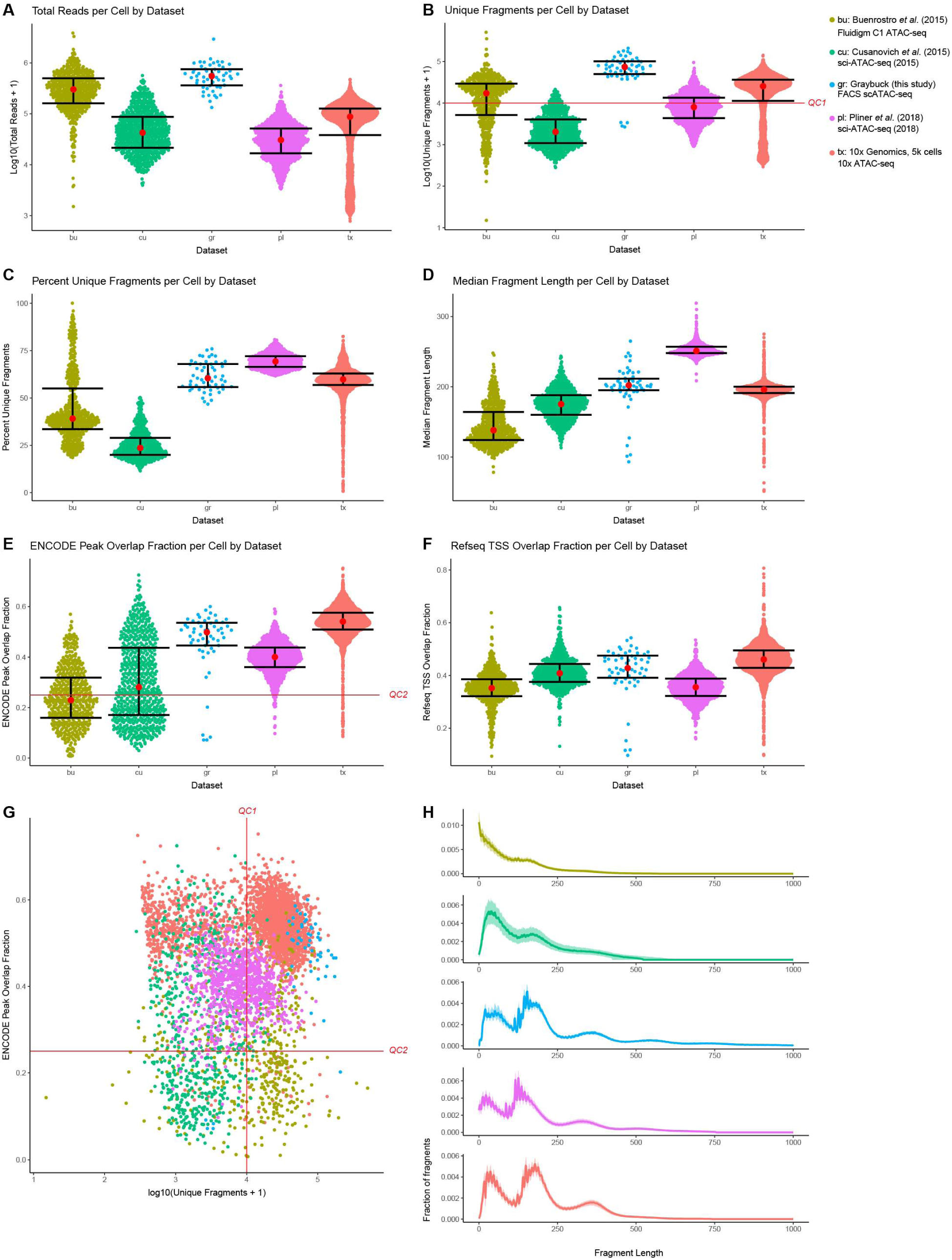
scATAC-seq data comparisons based on cell line GM12878. Comparison of FACS scATAC-seq libraries to those previously generated using Fluidigm C1 (Buenrostro et al., 2015), sci-ATAC-seq (Cusanovich et al., 2015; Pliner et al., 2018) or droplet-based indexing (10x Genomics) for which data using the common cell line of human GM12878 cells is available. To use in these comparisons, we generated scATAC-seq data using our FACS-based method for 60 GM12878 cells. For each published dataset, raw data was obtained from GEO and was aligned and analyzed using the same methods. For 10x Genomics, aligned fragment locations and metadata were obtained from the 10x genomics website for the “5k 1:1 mixture of fresh frozen human (GM12878) and mouse (A20) cells” dataset. Only samples labeled as GM12878 in sample metadata for each dataset were used for analysis. Colors and abbreviations at the top-right are used throughout the plots: **bu**, Buenrostro *et al.* (2015); **cu**, Cusanovich *et al.* (2015); **gr**, Graybuck *et al.* (this study); **pl**, Pliner *et al.* (2018), and **tx**, 10x Genomics. (**A**) Beeswarm plots showing the total reads per cell. For all similar plots (A-F), each point represents a single cell, with bars showing the 25^th^ and 75^th^ percentiles, and central red dots showing the median values. (**B**) Number of uniquely mapped fragments per cell for each dataset. (**C**) Percent uniquely mapped fragments for each dataset. A high fraction of unique fragments (as in Pliner *et al.*) suggests that deeper sequencing will yield additional useful data. (**D**) Median fragment length for each dataset. (**E**) Fraction of reads overlapping ENCODE DNase-seq data generated for GM12878 cells. (**F**) Fraction of reads overlapping RefSeq TSS regions (TSS ± 5kb) for each dataset. (**G**) Two-axis QC criteria plot, as in **Fig. 2**, showing the QC1 and QC2 cutoffs used for mouse cortical scATAC-seq data. (**H**) Aggregate fragment length frequency plots. Fragment length is shown on the x-axis, and the fraction of reads with fragments of each bp size was calculated for each sample in each dataset. For this analysis, the median fraction at each fragment size is shown as a solid line, with 25^th^ and 75^th^ percentiles shown in the shaded regions.

**Fig. S4.**
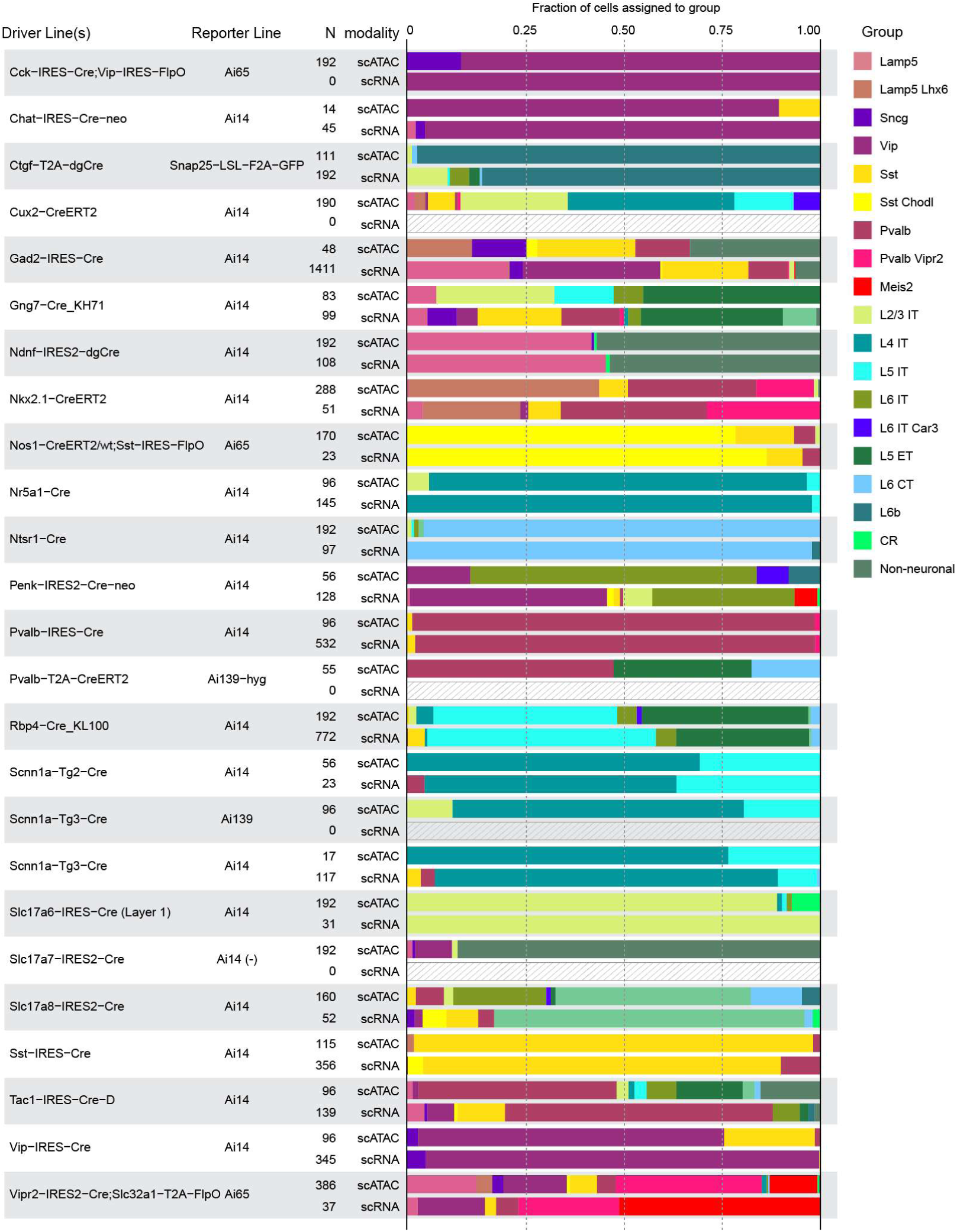
Comparison of scATAC-seq with scRNA-seq for same transgenic lines. To assess the accuracy of our scATAC-seq mapping to transcriptomic subclasses, we examined driver lines for which we collected both scATAC-seq and scRNA-seq data (the latter from Tasic *et al.* (Tasic et al., 2018)). For each combination of driver and reporter mouse line, we show the number of samples obtained for each method (scATAC-seq or scRNA-seq), and plot the fraction of cells assigned to each group used in **Figure 2** as colored bars for each method. Differences in proportions may be due to differences in dissection and region sampling strategies. Gray hashed regions indicate that no cells were assessed by scRNA-seq for specific driver and reporter line combinations.

**Fig. S5.**
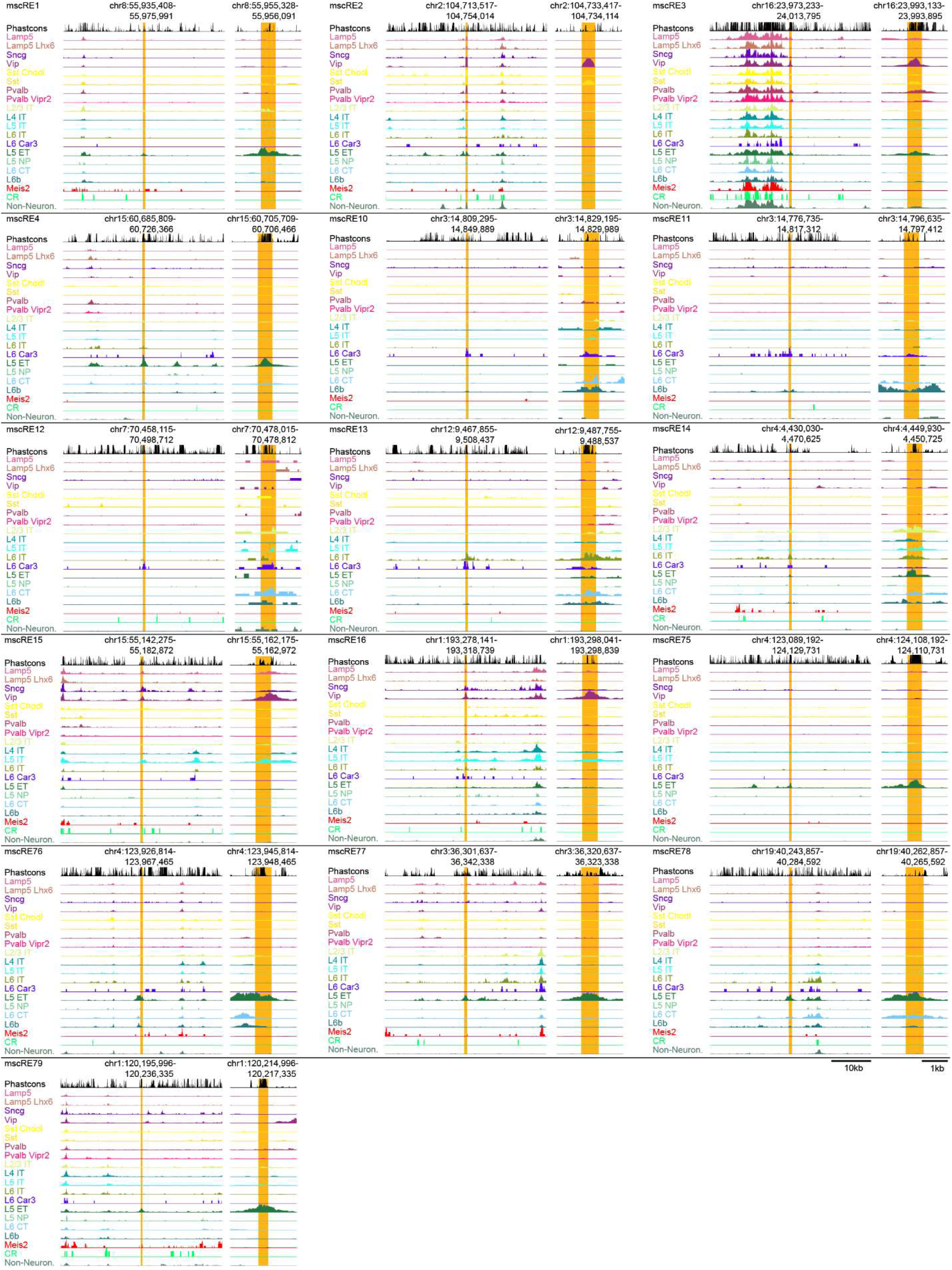
All mscREs examined in this study. Chromatin accessibility in clusters based on single cell ATAC-seq data for select genomic regions containing mscREs defined and examined in this study.

**Fig. S6.**
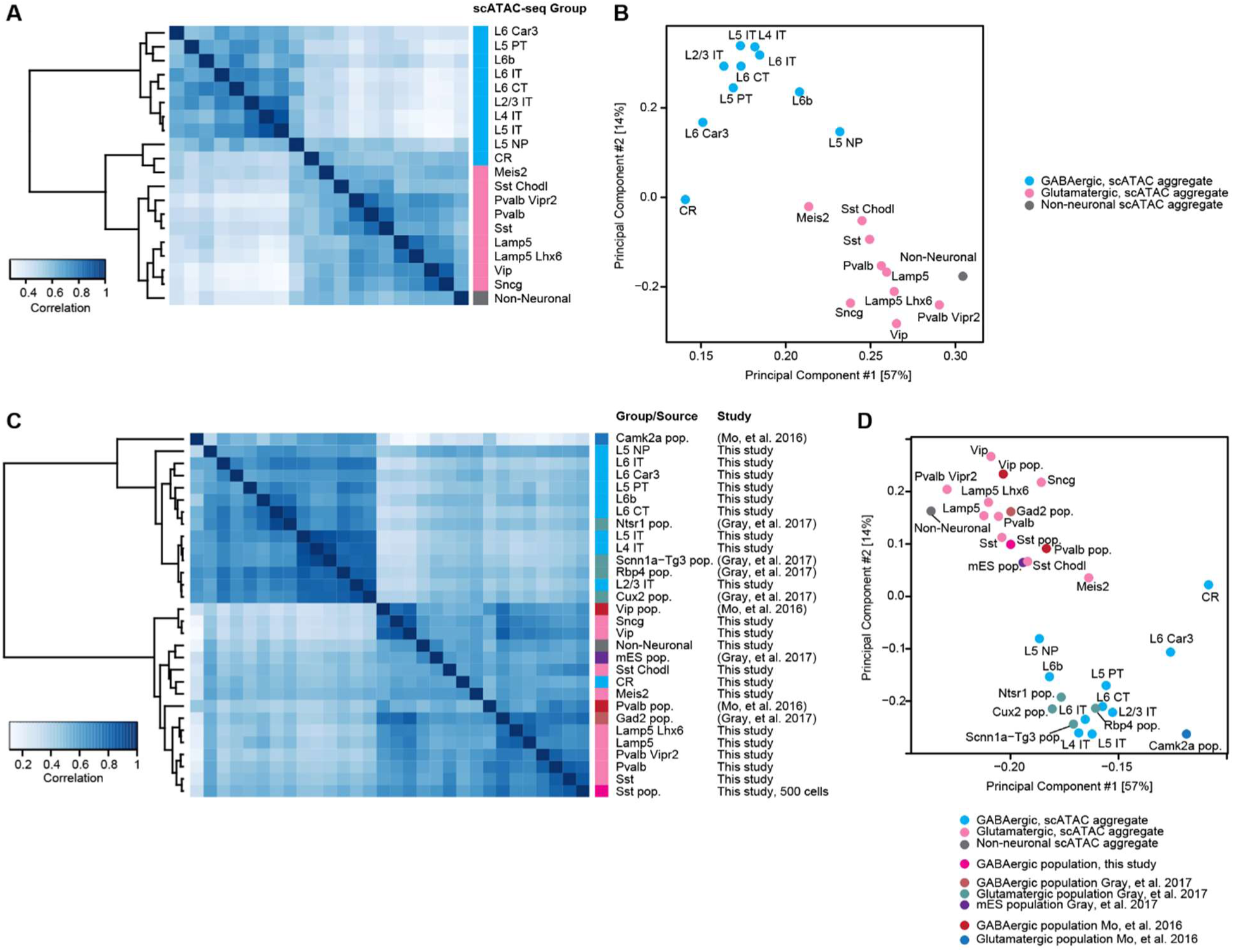
Comparison of our dataset with existing datasets. We compared aggregated scATAC-seq datasets to each other and to previously published neuronal ATAC-seq populations using the DiffBind package for R (Stark R, 2011) (**Methods**). (**A**) Comparisons of aggregated scATAC-seq datasets show expected grouping of GABAergic and glutamatergic cell classes. The heatmap shows weighted pairwise correlations between each pair of aggregated scATAC-seq groups. (**B**) PCA of the aggregated samples shown in (A). (**C**) Comparisons of aggregated scATAC-seq datasets with population ATAC-seq datasets generated by Mo *et al.* (Mo et al., 2015), Gray *et al.*, (Gray et al., 2017), or in this study (Sst pop.). (**D**) PCA of the datasets shown in (C). “pop.” indicates that the data were experimentally obtained from a population of cells, and not single cells.

**Fig. S7.**
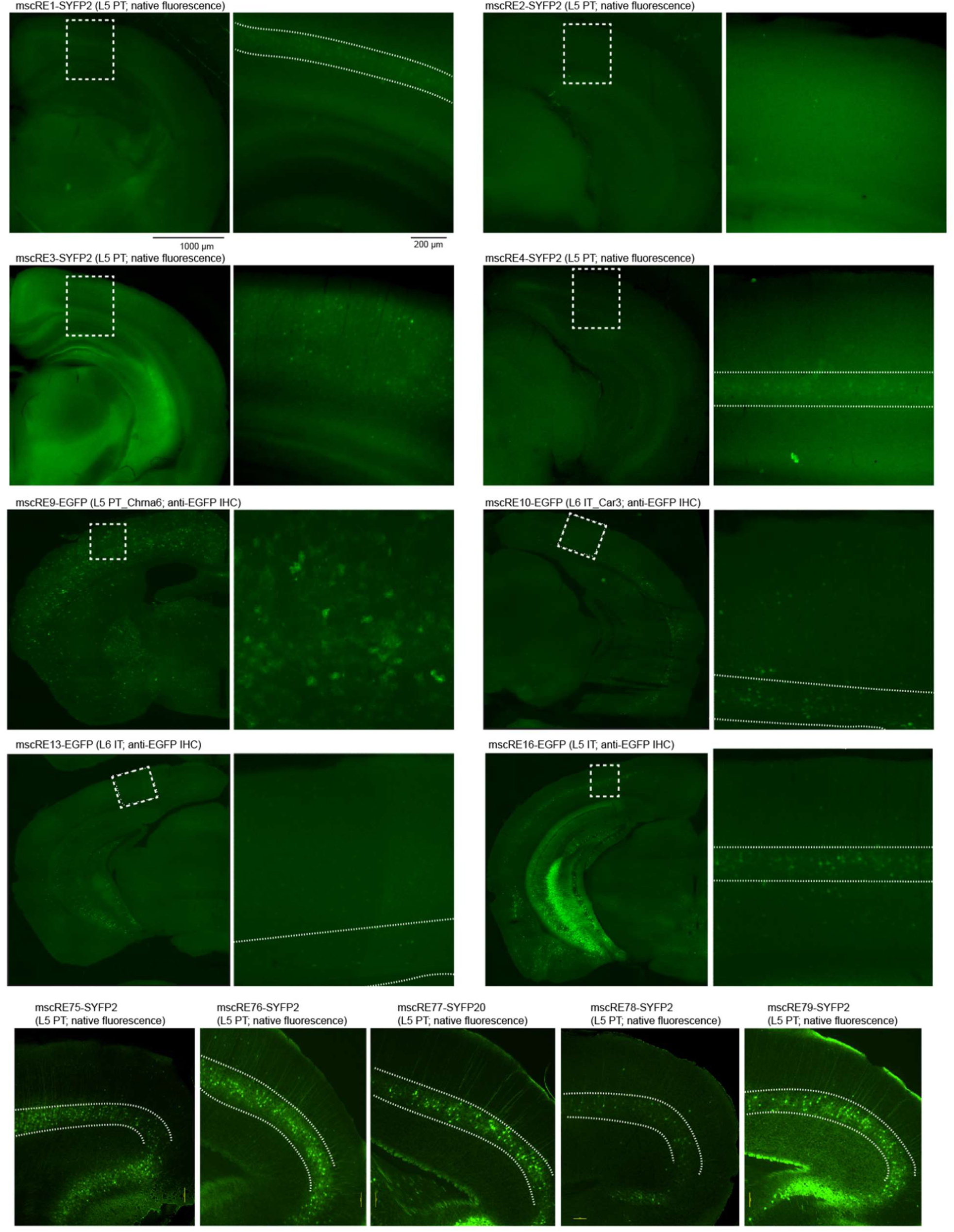
Initial screening of mscREs driving SYFP or EGFP. Candidate enhancers were first tested by retro-orbital injection of fluorophore-expressing viruses. The images show native fluorescence or antibody-enhanced immunohistochemistry (IHC) used to determine if fluorophores are expressed in an expected layer-specific pattern. Dashed boxes indicate regions of interest imaged at higher magnification (right panels) and dashed lines indicate the cortical layer expected to contain labeled cells. mscRE1-4 were screened using self-complementary AAVs (scAAV) driving SYFP2, whereas mscRE75-79 were screened using rAAVs driving SYFP2. Both groups were assessed using native fluorescence. mscRE10-16 were screened using rAAV driving EGFP, which was very weak, and needed anti-EGFP IHC to detect signal. For more detailed descriptions of viral constructs, see **Table S7**.

**Fig. S8.**
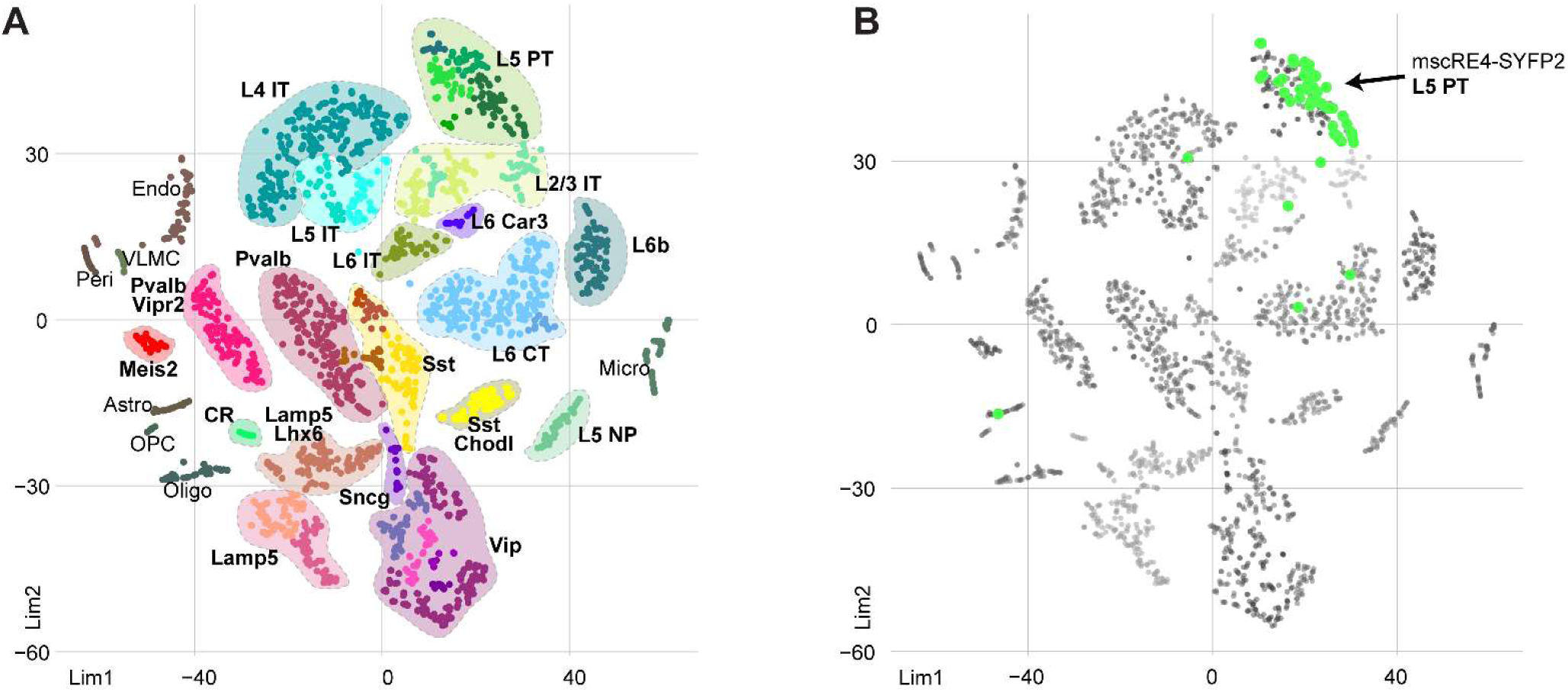
scATAC-seq of cells labeled by mscRE4-SYFP2. (**A**) scATAC-seq samples in a *t*-SNE projection with subclass and type labels for reference. (**B**) Cells in (A), highlighting samples collected from retro-orbital injection of mscRE4-SYFP2, VISp L5 dissection, and FACS (n = 61 high-quality scATAC-seq samples). 55 of 61 samples clustered with the L5 PT subclass.

**Fig. S9.**
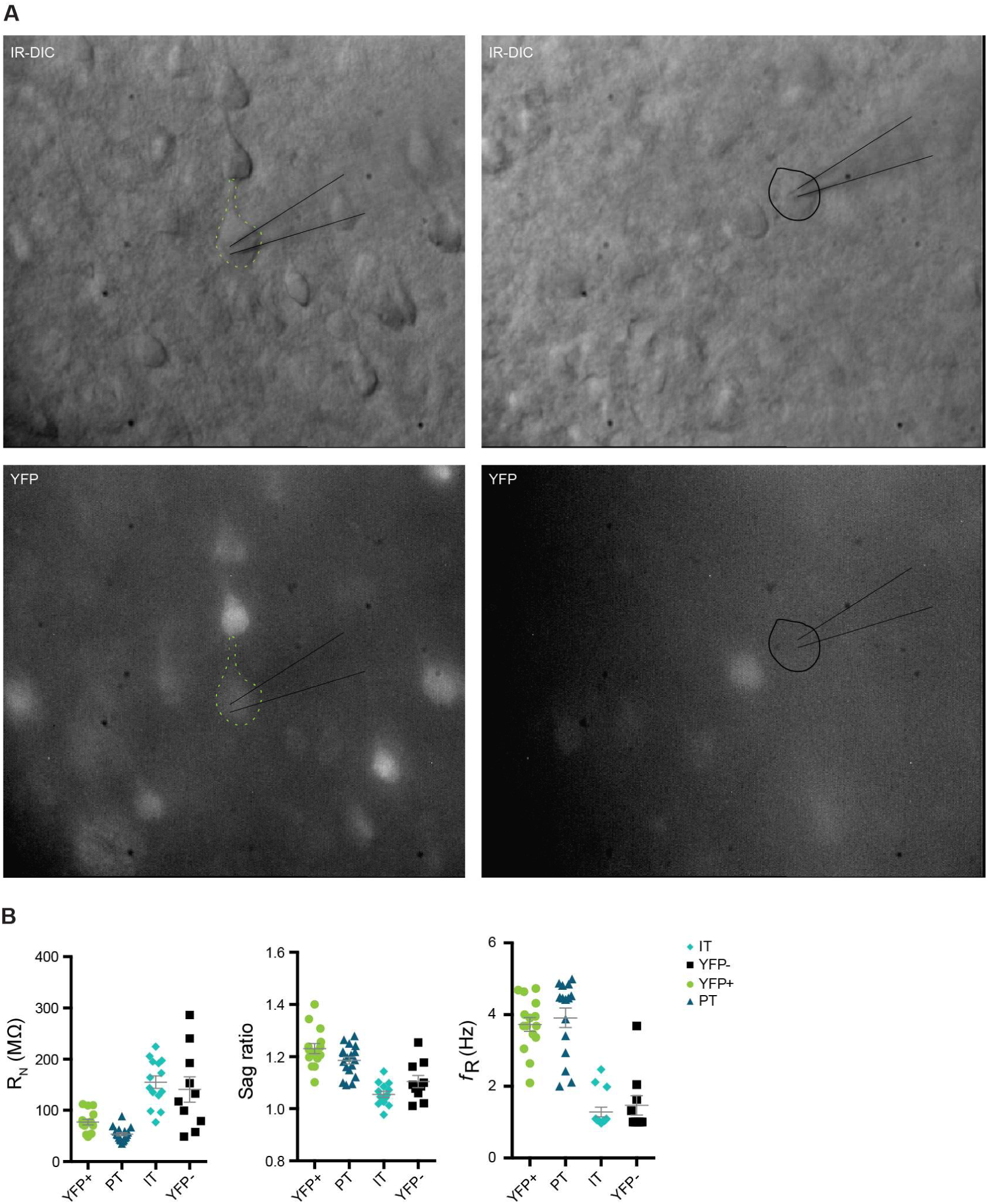
Additional electrophysiological characteristics of mouse neurons labeled by retro-orbital injection of mscRE4-SYFP2. (**A**) Microscopy of example cells characterized by patch electrophysiology. Left, a SYFP2-positive cell; right, a SFYP2-negative cell. Top images show differential interference contrast (DIC) microscopy of the same field shown in the bottom row. (**B**) Input resistance (R_N_), sag ratio, and resonance frequency (*f*_R_) for the four experimental conditions presented in **Figure 4D-E**.

**Fig. S10.**
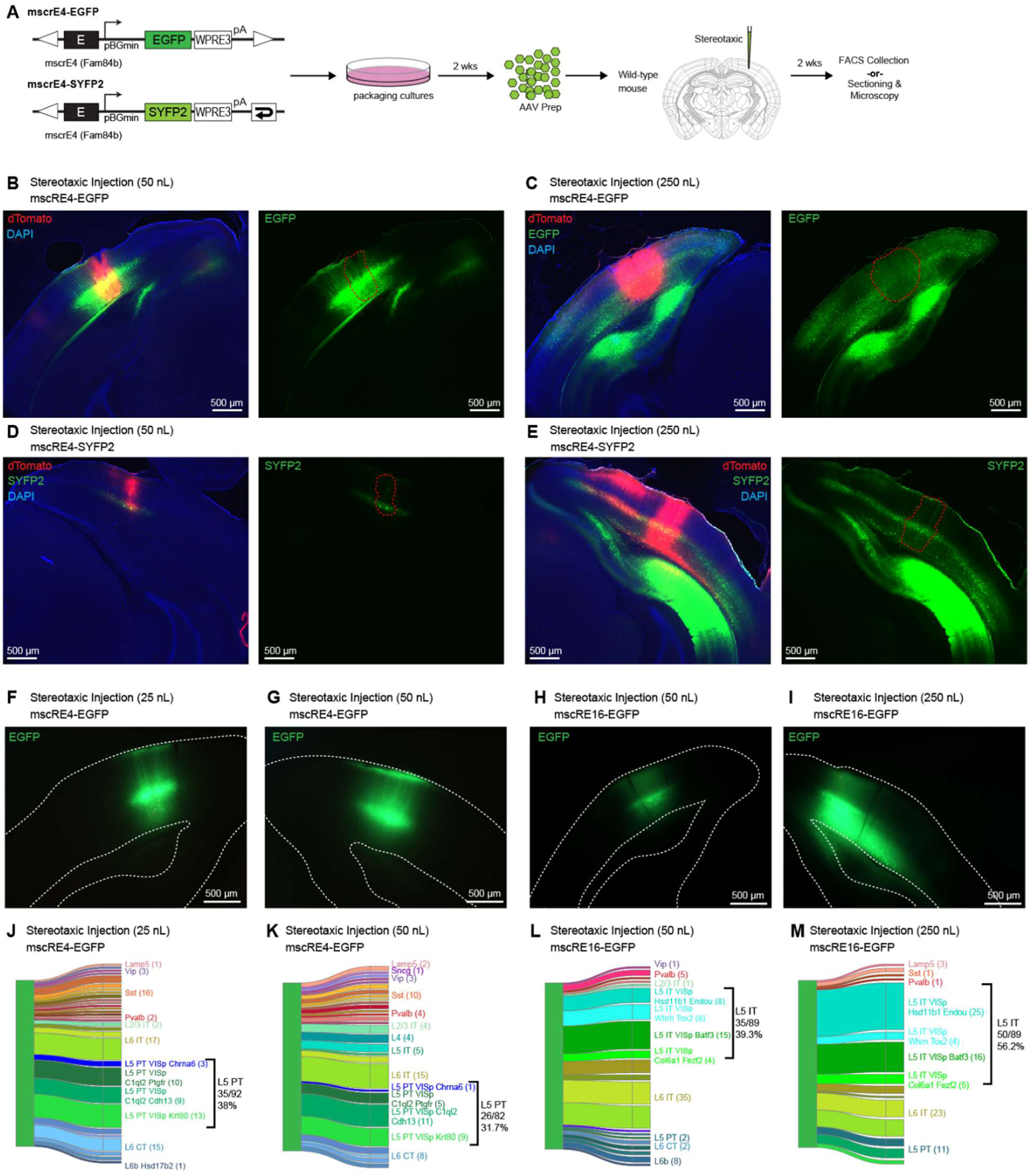
Enhancer virus labeling by stereotaxic injection. (**A**) For stereotaxic injection, enhancer-driven fluorophore viruses were generated by cloning mscRE sequences in rAAV (EGFP viruses) or scAAV (SYFP2 virus) constructs. After packaging, purification, and titering, viruses were stereotaxically targeted to VISp into wild-type mice. (**B-C**) Native fluorescence imaging of animals with stereotaxic injection of mscRE4-EGFP in VISp at two injection volumes (50 nL and 250 nL). Enhancer-driven viruses were co-injected with a constitutively expressed dTomato virus, rAAVDJ-EF1a-dTomato at 0.1X of the volumes of the mscRE viruses, to provide injection site location (red outlines). (**D-E**) Native fluorescence imaging of animals with stereotaxic injection of mscRE4-SYFP2 into primary visual cortex at two injection volumes (50 nL and 250 nL). The same injection site labeling strategy used in (A) was also utilized for these experiments. (**F-G**) Native fluorescence imaging of stereotaxic injections into the primary visual cortex using mscRE4-EGFP at 25 nL and 50 nL volumes injected into the left and right hemisphere, respectively. (**H-I**) as for (**E-F**) using mscRE16-EGFP at 50 nL and 250 nL volumes. (**J-M**) scRNA-seq mapping results for cells collected from panels above each river plot (e.g., cells in (**I**) were collected from the injections matching the conditions shown in (**E**) scRNA-seq data were mapped to a VISp cell type reference from Tasic *et al.* (Tasic et al., 2018). using a centroid classifier, and only cells that mapped to a single cluster in ≥ 80 of 100 bootstrapped classifications were retained for figures: (**J**), n = 92 of 96; (**K**) n = 82 of 96; (**L**) n = 89/96; and (**M**) n = 89/96.

**Fig. S11.**
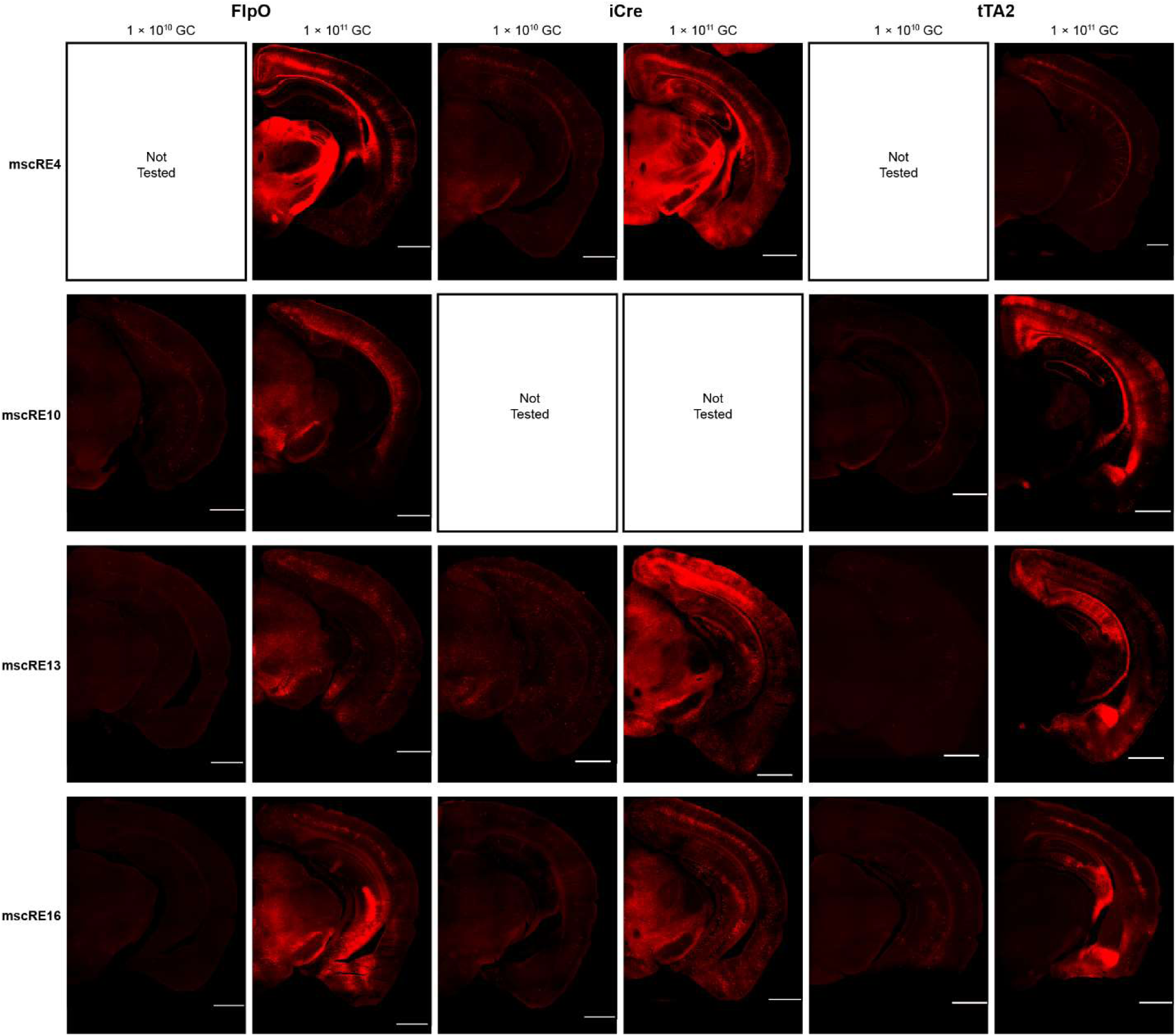
Labeling of retro-orbitally delivered mscRE viruses. Native fluorescence of tdTomato imaged in reporter mice injected retro-orbitally with enhancer-driven recombinase or tTA2; 1 ×10^10^ genome copies (GC) or 1×10^11^ GC of virus was delivered. Each enhancer is shown in a row, and each driver/reporter/dose combination is shown in a column. Reporter mouse lines used: *Ai65F* for FlpO, *Ai14* for iCre, and *Ai63* for tTA2. All scale bars (white) are 500 µm.

**Fig. S12.**
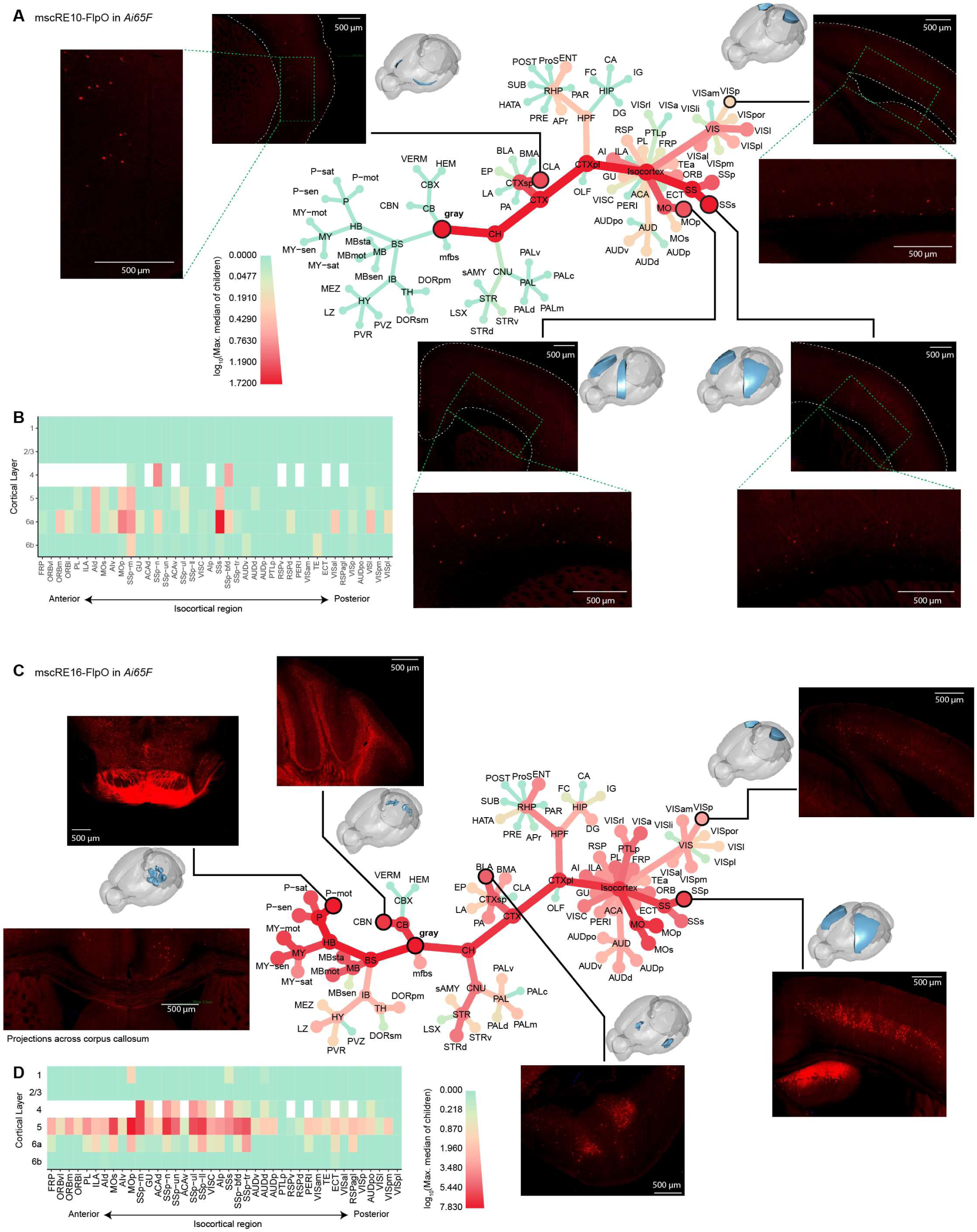
Whole-brain characterization of mscRE10-FlpO and mscRE16-FlpO. (**A**) TissueCyte imaging of an *Ai65F* mouse 2 weeks after retro-orbital injection of mscRE10-FlpO was registered to the Allen Institute Common Coordinate Framework (CCFv3), and each structure in the adult mouse structural ontology was scored. Similar to in **Figure 6**, this panel provides a high-level overview of cell labeling throughout the structural ontology. The size and color of nodes represents the maximum signal found among all children of each node, which allows one to follow the tree to the source of high signal within each structure. “Gray” is the root of the tree, representing all gray matter regions. Insets display selected regions of high or specific signal. (**B**) Layer quantification for the same TissueCyte image registered to the CCFv3 for all isocortical regions. Agranular regions that lack L4 are shown with a white box in the L4 row. Region acronyms correspond to the Adult Mouse Allen Brain Reference Atlas. (**C**) TissueCyte results for mscRE16-FlpO injected into *Ai65F*, as in (A). The inset at the bottom-left shows projection of IT neurons across the corpus callosum. (**D**) Heatmap of layer quantification for isocortical regions labeled by mscRE16-FlpO injected into *Ai65F*, as in (B).

**Fig. S13.**
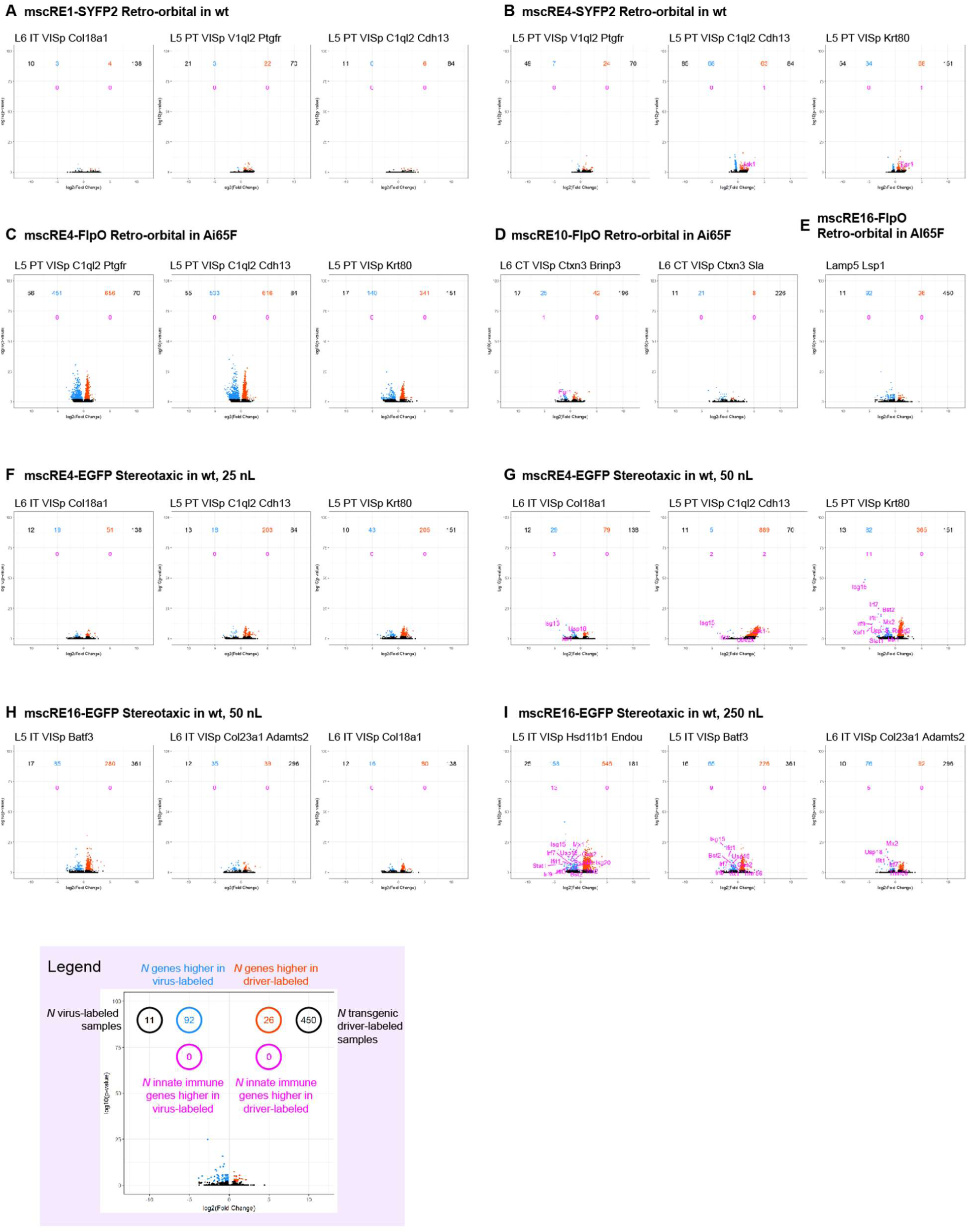
Differential gene expression induced by viral labeling. For each set of scRNA-seq data collected from virally labeled cells (**A–I**), we selected all cell types to which at least 10 cells were mapped. For each selected cell type, we performed pairwise differential gene expression analysis between virally labeled cells and cells labeled by transgenic recombinase driver mouse lines from the Tasic *et al.* (Tasic et al., 2018) (**Methods**). A volcano plot is shown for each comparison. Each gene is represented by a point. Log_2_(fold change) is displayed on the x-axis, with genes higher in virally labeled cells on the left and higher in transgenic mouse lines on the right. Log_10_(adjusted p-value) is displayed on the y-axis. Significantly differentially expressed genes (adjusted p-value < 0.01) are highlighted in blue (higher in virally labeled cells) and orange (higher in transgenic mouse lines). Differentially expressed innate immunity-related genes are highlighted in magenta. Counts for the number of cells in each group and the number of differentially expressed genes and immune-related genes are shown in each plot. The legend below panel (**H**) applies to all plots.

**Fig. S14.**
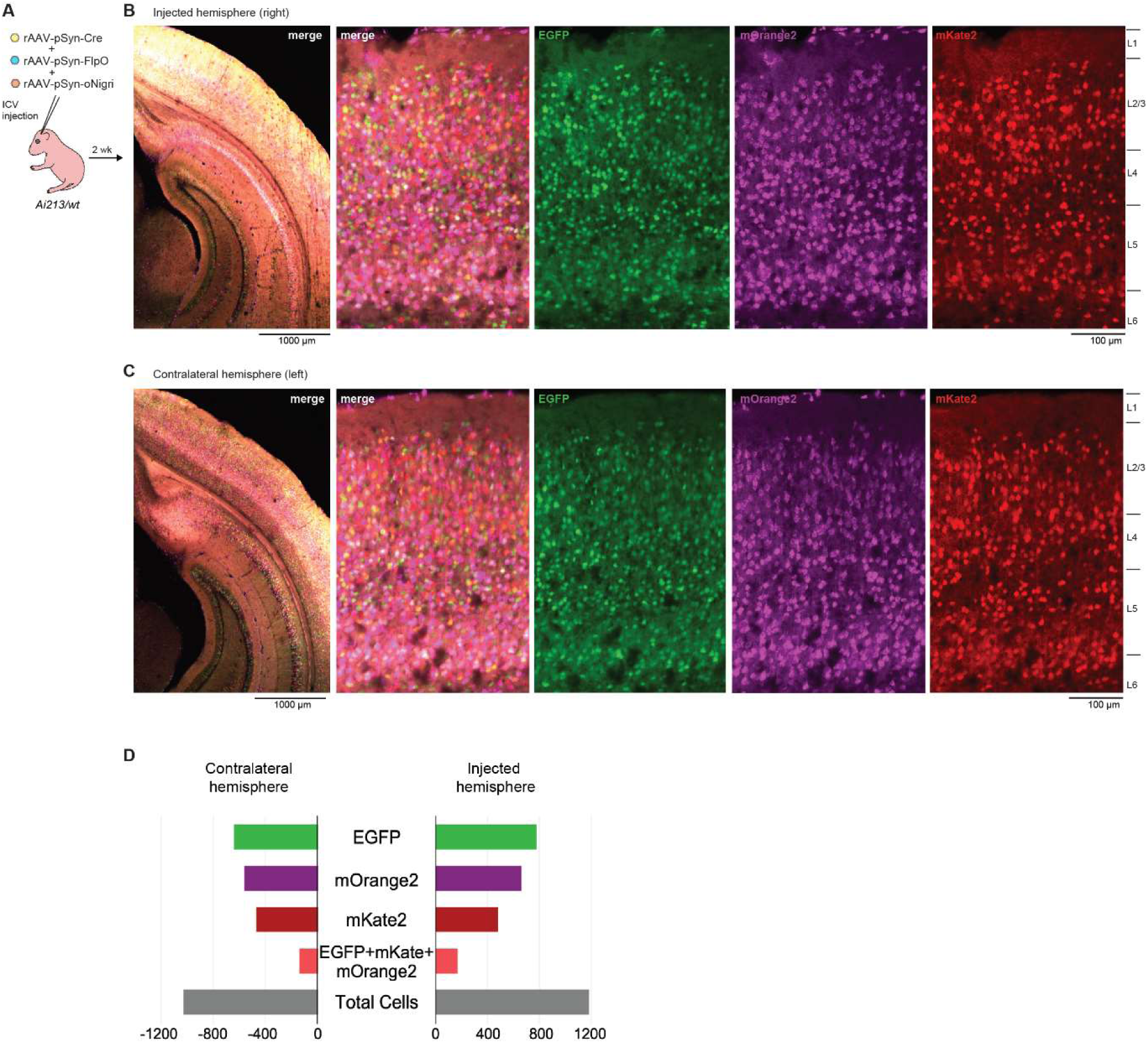
Increased labeling efficiency achieved with ICV injection of viruses. (**A-C**) *Ai213/wt* animals at postnatal day 3 (P3) were injected with a mixture of the three pan-neuronal recombinase viruses each at 2.1 × 10^10^ GC into the right cerebral ventricle and reporter expression in the brain was analyzed two-weeks post-injection. (**D**) The labelled cells for each fluorophore alone or the combination of all three were counted in confocal images collected from primary visual cortex of ICV injected animals. Quantification from both hemispheres is presented and shows a similar number of labeled cells in all groups between both regions.

**Fig. S15.**
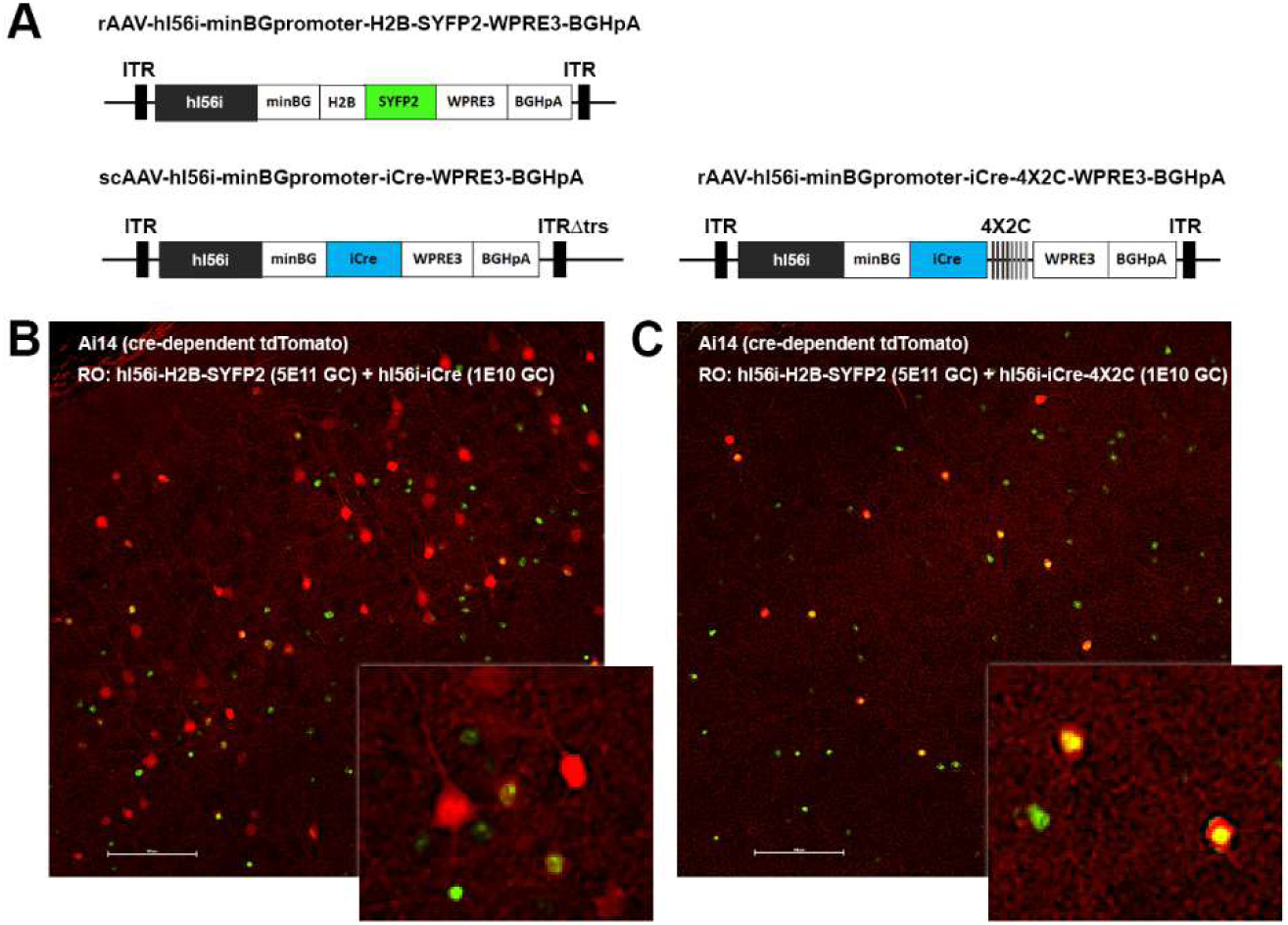
An optimized AAV vector for specific expression of Cre recombinase in forebrain GABAergic interneurons in the adult mouse brain. (**A**) Diagram of AAV vector designs. Top vector: *DLX* hl56i enhancer driving a histone 2B-tagged SYFP2 fluorophore; bottom left vector: hl56i enhancer driving iCre; bottom right vector: hl56i enhancer driving iCre with 3’ 4×2C mAGNET sequence to suppress excitatory neuron labeling (Sayeg et al., 2015). (**B-C**) Viruses were co-delivered (RO) into *Ai14* mice at the doses indicated and native tdTomato (red) and SYFP2 (green) fluorescence was evaluated in micrographs from adult mouse neocortex. Labeling of both excitatory and inhibitory populations was observed with hl56i-iCre virus (**B**), compared to exclusively GABAergic cell type labeling with hl56i-iCre-4×2C virus (**C**). Scale bars,100 microns.

## Supplementary Tables

**Table S1. Mouse lines.** Information about 27 driver lines and 5 reporter lines used in this study, with originating labs, references, generation method, RRID, and repository availability. All lines are available from the Jackson Laboratory or MMRRC, except for *Ai63*, which is available directly from the Allen Institute. The references included in this table are: (Daigle et al., 2018); (He et al., 2016); (Tasic et al., 2016); (Madisen et al., 2015); (Harris et al., 2014); (Gerfen et al., 2013); (Taniguchi et al., 2013); (Franco and Muller, 2013); (Taniguchi et al., 2011); (Vong et al., 2011);(Madisen et al., 2010); (Dhillon et al., 2006); (Hippenmeyer et al., 2005).

**Table S2. Donor metadata for cells labeled by transgenic mouse lines.** Information about 68 donor mice used for scATAC-seq experiments using cells labeled by transgenic drivers and reporters, with donor IDs, full genotypes, sex, birth date, age at euthanasia, final FACS gate used for sorting, the protease used for cell dissociation, and whether trehalose was included in dissociation and sorting buffers as described in **Methods**.

**Table S3. Stereotaxic,retro-orbital, and intracerebroventricular injection donor metadata.** Information about 14 stereotaxic and 33 retroorbital injection experiments, with donor IDs, full genotypes, sex, age at injection, age at euthanasia, virus incubation time, injected material, injection target, Paxinos stereotaxic coordinates (Paxinos and Franklin, 2013), injection titers and volumes, and the experimental modalities for which each donor was used.

**Table S4. scATAC-seq cell alignment and QC statistics.** Metadata for 3,603 scATAC-seq samples, including sample IDs, donor IDs corresponding to **table S2** or **table S3**, dissected region, dissected cortical layer(s), dissection date, age at dissection, MiSeq batch, total sequenced reads, number of mapped reads, number of mapped fragments, percent of reads mapped to the mm10 genome, percent of mapped reads, percent of unmapped reads, number of unique reads, number of unique fragments (used for QC1), percent unique fragments, percent duplicate fragments, number of unique fragments overlapping ENCODE DNase-Seq peaks, fraction of unique fragments overlapping ENCODE DNase-Seq peaks (used for QC2), fraction of unique fragments with insert size > 250bp (used for QC3), and pass/fail flags for QC criteria.

**Table S5. scATAC-seq clustering, cell type mapping, and *t*-SNE embedding coordinates.** Results for the 2,416 scATAC-seq samples passing QC criteria that were used for *t*-SNE embedding, Phenograph clustering, and mapping to scRNA-seq cell types, including scATAC-seq sample IDs corresponding to **Table S4**, group IDs, labels, and colors used for plots in figures, Phenograph cluster IDs and colors used for scRNA-seq correlation, correlation scores for the highest-scoring scRNA-seq cluster for each phenograph cluster, cluster IDs, cluster labels, and cluster colors used for plotting, and *t*-SNE embedding coordinates (tSNE1 and tSNE2).

**Table S6. mscRE genomic locations and cloning primers.** Information about 16 putative regulatory elements that were cloned and tested, with region IDs, coordinates in the mm10 genome, targeted cell subclass/type populations, nearest gene, cloned genomic sequence length, and 5’ and 3’ primer sequences used for cloning.

**Table S7. Virus descriptions and sources.** Information about 23 enhancer-driven viruses and two additional viruses used for retrograde and stereotaxic injections, including full virus names, abbreviations used throughout the manuscript, virus sources, serotypes, and references for packaging lines and viral genomes for viruses from other studies. The references in this table are:(Chan et al., 2017); (Hnasko et al., 2006); (Wertz et al., 2015); and (Grimm et al., 2008).

**Table S8. scRNA-seq sample alignment and mapping statistics.** Metadata for 989 scRNA-seq samples labeled by enhancer-driven viruses by stereotaxic or retro-orbital injections, including experiment IDs, sample IDs, mapping results for cell type assignments based on Tasic et al., (Tasic et al., 2018) (mapping confidence, cluster IDs, cluster labels, cluster colors, class IDs, class labels, class colors, subclass IDs, subclass labels, and subclass colors), donor metadata (donor IDs, sex, and genotype), injection type (ro = retro-orbital; st = stereotaxic), injected virus, dissected hemisphere, dissected ROI, sorted FACS population, FACS container (an ID for each FACS strip), FACS well, RNA amplification set ID and library prep set ID (batches used during sample processing), number of PCR cycles, fraction of cDNA with length > 400 bp, RNA amplification pass/fail flag, ng of amplified cDNA, library multiplexing level, sequencing batch ID, pooled library tube ID, average final sequence length, sequencing sample quantification (quantification2_ng, quantification_fmol, quantification2_nM), library prep pass/fail flag, alignment statistics (total reads, percent of reads aligned to exons/rRNA/tRNA/introns/intergenic regions/E. coli/synthetic constructs, percent unique reads, percent aligned to any listed target), number of genes detected (including intronic reads = premRNA_genes_detected; including only exonic reads = mRNA_genes_detected).

**Table S9. Statistics for RNAscope.** Counts, thresholds, and on/off-target criteria, counts, and percentages for each RNA ISH experiment. Probe channel, specificity, and cutoffs are shown in columns beginning with R[round number]C[channel number] (e.g. R1C1 Probe is the probe used in round 1, channel 1). Probes are described using fluorescence wavelength and target gene (e.g. 488 Fam84b corresponds to a probe for the *Fam84b* gene with 488 nm excitation).

## References

Arnold, C.D., Gerlach, D., Stelzer, C., Boryn, L.M., Rath, M., and Stark, A. (2013). Genome-wide quantitative enhancer activity maps identified by STARR-seq. Science (80-.). 339, 1074–1077.

Baker, A., Kalmbach, B., Morishima, M., Kim, J., Juavinett, A., Li, N., and Dembrow, N. (2018). Specialized Subpopulations of Deep-Layer Pyramidal Neurons in the Neocortex: Bridging Cellular Properties to Functional Consequences. J. Neurosci. 38, 5441–5455.

Blankvoort, S., Witter, M.P., Noonan, J., Cotney, J., and Kentros, C. (2018). Marked Diversity of Unique Cortical Enhancers Enables Neuron-Specific Tools by Enhancer-Driven Gene Expression. Curr. Biol. 28, 2103-2114.e5.

Buenrostro, J.D., Wu, B., Litzenburger, U.M., Ruff, D., Gonzales, M.L., Snyder, M.P., Chang, H.Y., and Greenleaf, W.J. (2015). Single-cell chromatin accessibility reveals principles of regulatory variation. Nature 523, 486–490.

Cao, J., Cusanovich, D.A., Ramani, V., Aghamirzaie, D., Pliner, H.A., Hill, A.J., Daza, R.M., McFaline-Figueroa, J.L., Packer, J.S., Christiansen, L., et al. (2018). Joint profiling of chromatin accessibility and gene expression in thousands of single cells. Science (80-).. 1385, 1380–1385.

Chan, K.Y., Jang, M.J., Yoo, B.B., Greenbaum, A., Ravi, N., Wu, W., Sánchez-guardado, L., Lois, C., Mazmanian, S.K., Deverman, B.E., et al. (2017). Engineered adeno-associated viruses for efficient and noninvasive gene delivery throughout the central and peripheral nervous systems. Nat. Neurosci. 20, 1172–1179.

Chen, S., Lake, B.B., and Zhang, K. (2019). High-throughput sequencing of the transcriptome and chromatin accessibility in the same cell. Nat. Biotechnol. 37, 1452–1457.

Cusanovich, D. a, Daza, R., Adey, A., Pliner, H. a, Christiansen, L., Gunderson, K.L., Steemers, F.J., Trapnell, C., and Shendure, J. (2015). Epigenetics. Multiplex single-cell profiling of chromatin accessibility by combinatorial cellular indexing. Science 348, 910–914.

Cusanovich, D.A., Reddington, J.P., Garfield, D.A., Daza, R.M., Aghamirzaie, D., Marco-Ferreres, R., Pliner, H.A., Christiansen, L., Qiu, X., Steemers, F.J., et al. (2018). The cis-regulatory dynamics of embryonic development at single-cell resolution. Nature 555, 538–542.

Daigle, T.L., Madisen, L., Hage, T.A., Valley, M.T., Knoblich, U., Larsen, R.S., Takeno, M.M., Huang, L., Gu, H., Larsen, R., et al. (2018). A Suite of Transgenic Driver and Reporter Mouse Lines with Enhanced Brain-Cell-Type Targeting and Functionality. Cell 174, 465-480.e22.

Dembrow, N.C., Chitwood, R.A., and Johnston, D. (2010). Projection-specific neuromodulation of medial prefrontal cortex neurons. J. Neurosci. 30, 16922–16937.

Dickel, D.E., Ypsilanti, A.R., Pla, R., Zhu, Y., Barozzi, I., Mannion, B.J., Khin, Y.S., Fukuda-Yuzawa, Y., Plajzer-Frick, I., Pickle, C.S., et al. (2018). Ultraconserved Enhancers Are Required for Normal Development. Cell 172, 491-499.e15.

Dimidschstein, J., Chen, Q., Tremblay, R., Rogers, S.L., Saldi, G.-A., Guo, L., Xu, Q., Liu, R., Lu, C., Chu, J., et al. (2016). A viral strategy for targeting and manipulating interneurons across vertebrate species. Nat. Neurosci. 19, 1743–1749.

Economo, M.N., Viswanathan, S., Tasic, B., Bas, E., Winnubst, J., Menon, V., Graybuck, L.T., Nguyen, T.N., Smith, K.A., Yao, Z., et al. (2018). Distinct descending motor cortex pathways and their roles in movement. Nature 563, 79–84.

Gasperini, M., Tome, J.M., and Shendure, J. (2020). Towards a comprehensive catalogue of validated and target-linked human enhancers. Nat. Rev. Genet. 40.

Gong, S., Doughty, M., Harbaugh, C.R., Cummins, A., Hatten, M.E., Heintz, N., and Gerfen, C.R. (2007). Targeting Cre recombinase to specific neuron populations with bacterial artificial chromosome constructs. J. Neurosci. 27, 9817–9823.

Gray, L.T., Yao, Z., Nguyen, T.N., Kim, T.-K., Zeng, H., and Tasic, B. (2017). Layer-specific chromatin accessibility landscapes reveal regulatory networks in adult mouse visual cortex. Elife e21883.

Harris, J.A., Mihalas, S., Hirokawa, K.E., Whitesell, J.D., Knox, J., Bernard, A., Bohn, P., Caldejon, S., Casal, L., Cho, A., et al. (2018). The organization of intracortical connections by layer and cell class in the mouse brain. BioRxiv 292961.

Hartl, D., Krebs, A.R., Jüttner, J., Roska, B., and Schübeler, D. (2017). Cis-regulatory landscapes of four cell types of the retina. Nucleic Acids Res. 45, 11607–11621.

Jüttner, J., Szabo, A., Gross-Scherf, B., Morikawa, R.K., Rompani, S.B., Hantz, P., Szikra, T., Esposti, F., Cowan, C.S., Bharioke, A., et al. (2019). Targeting neuronal and glial cell types with synthetic promoter AAVs in mice, non-human primates and humans. Nat. Neurosci. doi.org/10.1101/434720.

Karimova, M., Splith, V., Karpinski, J., Pisabarro, M.T., and Buchholz, F. (2016). Discovery of Nigri/nox and Panto/pox site-specific recombinase systems facilitates advanced genome engineering. Sci. Rep. 6, 1–13.

Kelsch, W., Stolfi, A., and Lois, C. (2012). Genetic labeling of neuronal subsets through enhancer trapping in mice. PLoS One 7, 3–10.

Kishi, J.Y., Lapan, S.W., Beliveau, B.J., West, E.R., Zhu, A., Sasaki, H.M., Saka, S.K., Wang, Y., Cepko, C.L., and Yin, P. (2019). SABER amplifies FISH: enhanced multiplexed imaging of RNA and DNA in cells and tissues. Nat. Methods 16, 533–544.

Klemm, S.L., Shipony, Z., and Greenleaf, W.J. (2019). Chromatin accessibility and the regulatory epigenome. Nat. Rev. Genet. 20, 207–220.

Levine, J.H., Simonds, E.F., Bendall, S.C., Davis, K.L., Amir, E.A.D., Tadmor, M.D., Litvin, O., Fienberg, H.G., Jager, A., Zunder, E.R., et al. (2015). Data-Driven Phenotypic Dissection of AML Reveals Progenitor-like Cells that Correlate with Prognosis. Cell 162, 184–197.

Liu, L., Liu, C., Quintero, A., Wu, L., Yuan, Y., Wang, M., Cheng, M., Leng, L., Xu, L., Dong, G., et al. (2019). Deconvolution of single-cell multi-omics layers reveals regulatory heterogeneity. Nat. Commun. 10.

Luo, L., Callaway, E.M., and Svoboda, K. (2018). Genetic Dissection of Neural Circuits: A Decade of Progress. Neuron 98, 256–281.

Madisen, L., Zwingman, T.A., Sunkin, S.M., Oh, S.W., Zariwala, H.A., Gu, H., Ng, L.L., Palmiter, R.D., Hawrylycz, M.J., Jones, A.R., et al. (2010). A robust and high-throughput Cre reporting and characterization system for the whole mouse brain. Nat. Neurosci. 13, 133–140.

Madisen, L., Garner, A.R.R., Shimaoka, D., Chuong, A.S.S., Klapoetke, N.C.C., Li, L., van der Bourg, A., Niino, Y., Egolf, L., Monetti, C., et al. (2015). Transgenic Mice for Intersectional Targeting of Neural Sensors and Effectors with High Specificity and Performance. Neuron 85, 942–958.

Mich, J.K., Hess, E.E., Graybuck, L.T., Somasundaram, S., Miller, A., Ding, Y., Shapovalova, N. V, Fong, O., Yao, S., Mortrud, M., et al. (2019). Epigenetic landscape and AAV targeting of human neocortical cell classes. BioRxiv.

Mo, A., Mukamel, E.A., Davis, F.P., Luo, C., Henry, G.L., Picard, S., Urich, M.A., Nery, J.R., Sejnowski, T.J., Lister, R., et al. (2015). Epigenomic Signatures of Neuronal Diversity in the Mammalian Brain. Neuron 86, 1369–1384.

Nagano, T., Lubling, Y., Stevens, T.J., Schoenfelder, S., Yaffe, E., Dean, W., Laue, E.D., Tanay, A., and Fraser, P. (2013). Single-cell Hi-C reveals cell-to-cell variability in chromosome structure. Nature 502, 59–64.

Pfeiffer, B.D., Jenett, A., Hammonds, A.S., Ngo, T.-T.B., Misra, S., Murphy, C., Scully, A., Carlson, J.W., Wan, K.H., Laverty, T.R., et al. (2008). Tools for neuroanatomy and neurogenetics in Drosophila. Proc. Natl. Acad. Sci. 105, 9715–9720.

Pfeiffer, B.D., Ngo, T.T.B., Hibbard, K.L., Murphy, C., Jenett, A., Truman, J.W., and Rubin, G.M. (2010). Refinement of tools for targeted gene expression in Drosophila. Genetics 186, 735–755.

Pliner, H.A., Packer, J.S., McFaline-Figueroa, J.L., Cusanovich, D.A., Daza, R.M., Aghamirzaie, D., Srivatsan, S., Qiu, X., Jackson, D., Minkina, A., et al. (2018). Cicero Predicts cis-Regulatory DNA Interactions from Single-Cell Chromatin Accessibility Data. Mol. Cell 71, 858-871.e8.

Porrero, C., Rubio-Garrido, P., Avendaño, C., and Clascá, F. (2010). Mapping of fluorescent protein-expressing neurons and axon pathways in adult and developing Thy1-eYFP-H transgenic mice. Brain Res. 1345, 59–72.

Preissl, S., Fang, R., Huang, H., Zhao, Y., Raviram, R., Gorkin, D.U., Zhang, Y., Sos, B.C., Afzal, V., Dickel, D.E., et al. (2018). Single-nucleus analysis of accessible chromatin in developing mouse forebrain reveals cell-type-specific transcriptional regulation. Nat. Neurosci. 21, 432–439.

Ramani, V., Deng, X., Gunderson, K.L., Steemers, F.J., Disteche, C.M., Noble, W.S., Duan, Z., and Shendure, J. (2016). Massively multiplex single-cell Hi-C. BioRxiv 065052.

Salva, M.Z., Himeda, C.L., Tai, P.W.L., Nishiuchi, E., Gregorevic, P., Allen, J.M., Finn, E.E., Nguyen, Q.G., Blankinship, M.J., Meuse, L., et al. (2007). Design of tissue-specific regulatory cassettes for high-level rAAV-mediated expression in skeletal and cardiac muscle. Mol. Ther. 15, 320–329.

Shen, S.Q., Myers, C.A., Hughes, A.E.O., Byrne, L.C., Flannery, J.G., and Corbo, J.C. (2016). Massively parallel cis-regulatory analysis in the mammalian central nervous system. Genome Res. 26, 238–255.

Shima, Y., Sugino, K., Hempel, C.M., Shima, M., Taneja, P., Bullis, J.B., Mehta, S., Lois, C., and Nelson, S.B. (2016). A mammalian enhancer trap resource for discovering and manipulating neuronal cell types. Elife 5, 1–32.

Siepel, A., Bejerano, G., Pedersen, J.S., Hinrichs, A.S., Hou, M., Rosenbloom, K., Clawson, H., Spieth, J., Hillier, L.D.W., Richards, S., et al. (2005). Evolutionarily conserved elements in vertebrate, insect, worm, and yeast genomes. Genome Res. 15, 1034–1050.

Sorensen, S. a, Bernard, A., Menon, V., Royall, J.J., Glattfelder, K.J., Desta, T., Hirokawa, K., Mortrud, M., Miller, J. a, Zeng, H., et al. (2013). Correlated Gene Expression and Target Specificity Demonstrate Excitatory Projection Neuron Diversity. Cereb. Cortex 1–17.

Taniguchi, H., He, M., Wu, P., Kim, S., Paik, R., Sugino, K., Kvitsani, D., Fu, Y., Lu, J., Lin, Y., et al. (2011). A Resource of Cre Driver Lines for Genetic Targeting of GABAergic Neurons in Cerebral Cortex. Neuron 71, 995–1013.

Tasic, B. (2018). Single cell transcriptomics in neuroscience: cell classification and beyond. Curr. Opin. Neurobiol. 50, 242–249.

Tasic, B., Menon, V., Nguyen, T.N, Kim, T.K., Jarsky, T., Yao, Z., Levi, B.B., Gray, L.T., Sorensen, S.A., Dolbeare, T., et al. (2016). Adult mouse cortical cell taxonomy revealed by single cell transcriptomics. Nat. Neurosci. 19, 335–346.

Tasic, B., Yao, Z., Graybuck, L.T., Smith, K.A., Nguyen, T.N., Bertagnolli, D., Goldy, J., Garren, E., Economo, M.N., Viswanathan, S., et al. (2018). Shared and distinct transcriptomic cell types across neocortical areas. Nature 563, 72–78.

Visel, A., Taher, L., Girgis, H., May, D., Golonzhka, O., Hoch, R.V., McKinsey, G.L., Pattabiraman, K., Silberberg, S.N., Blow, M.J., et al. (2013). A High-Resolution Enhancer Atlas of the Developing Telencephalon. Cell 152, 895–908.

Vormstein-Schneider, D., Lin, J., Pelkey, K., Chittajallu, R., Guo, B., Garcia, M.A., Sakopoulos, S., Stevenson, O., Schneider, G., Zhang, Q., et al. (2019). Viral manipulation of functionally distinct neurons from mice to humans. BioRxiv 808170.

Yee, S.P., and Rigby, P.W.J. (1993). The regulation of myogenin gene expression during the embryonic development of the mouse. Genes Dev. 7, 1277–1289.

Yue, F., Cheng, Y., Breschi, A., Vierstra, J., Wu, W., Ryba, T., Sandstrom, R., Ma, Z., Davis, C., Pope, B.D., et al. (2014). A comparative encyclopedia of DNA elements in the mouse genome. Nature 515, 355–364.

Zeng, H., and Sanes, J.R. (2017). Neuronal cell-type classification: Challenges, opportunities and the path forward. Nat. Rev. Neurosci. 18, 530–546.

Zeng, H., Horie, K., Madisen, L., Pavlova, M.N., Gragerova, G., Rohde, A.D., Schimpf, B.A., Liang, Y., Ojala, E., Kramer, F., et al. (2008). An inducible and reversible mouse genetic rescue system. PLoS Genet. 4.

## References

Adler, D., Murdoch, D., et al. (2018). rgl: 3D Visualization Using OpenGL.

Attali, D.a.B. C. (2018). ggExtra: Add Marginal Histograms to “ggplot2”, And more “ggplot2” Enhancements.

Baker, A., Kalmbach, B., Morishima, M., Kim, J., Juavinett, A., Li, N., and Dembrow, N. (2018). Specialized Subpopulations of Deep-Layer Pyramidal Neurons in the Neocortex: Bridging Cellular Properties to Functional Consequences. J Neurosci 38, 5441–5455.

Bates, D., and Maechler, M. (2018). Matrix: Sparse and Dense Matrix Classes and Methods.

Bengtsson, H. (2018). matrixStats: Functions that Apply to Rows and Columns of Matrices (and to Vectors).

Chan, K.Y., Jang, M.J., Yoo, B.B., Greenbaum, A., Ravi, N., Wu, W.L., Sanchez-Guardado, L., Lois, C., Mazmanian, S.K., Deverman, B.E., et al. (2017). Engineered AAVs for efficient noninvasive gene delivery to the central and peripheral nervous systems. Nat Neurosci 20, 1172–1179.

Chen, H. (2015). Rphenograph: R implementation of the phenograph algorithm.

Clark, E., and Sherrill-Mix, S. (2017). ggbeeswarm: Categorical scatter (Violin plots) Plots.

Cusanovich, D.A., Daza, R., Adey, A., Pliner, H.A., Christiansen, L., Gunderson, K.L., Steemers, F.J., Trapnell, C., and Shendure, J. (2015). Multiplex single cell profiling of chromatin accessibility by combinatorial cellular indexing. Science 348, 910–914.

Daigle, T.L., Madisen, L., Hage, T.A., Valley, M.T., Knoblich, U., Larsen, R.S., Takeno, M.M., Huang, L., Gu, H., Larsen, R., et al. (2018). A Suite of Transgenic Driver and Reporter Mouse Lines with Enhanced Brain-Cell-Type Targeting and Functionality. Cell 174, 465–480 e422.

Dhillon, H., Zigman, J.M., Ye, C., Lee, C.E., McGovern, R.A., Tang, V., Kenny, C.D., Christiansen, L.M., White, R.D., Edelstein, E.A., et al. (2006). Leptin directly activates SF1 neurons in the VMH, and this action by leptin is required for normal body-weight homeostasis. Neuron 49, 191–203.

Dobin, A., Davis, C.A., Schlesinger, F., Drenkow, J., Zaleski, C., Jha, S., Batut, P., Chaisson, M., and Gingeras, T.R. (2013). STAR: ultrafast universal RNA-seq aligner. Bioinformatics 29, 15–21.

Dowle, M., Srinivasan, A. (2019). data.table: Extension of `data.framè.

Foster, Z.S.L., Sharpton, T., Grunwald, N.J. (2016). MetacodeR : An R package for manipulation and heat tree visualization of community taxonomic data from metabarcoding. BioRxiv.

Franco, S.J., and Muller, U. (2013). Shaping our minds: stem and progenitor cell diversity in the mammalian neocortex. Neuron 77, 19–34.

George, S.H., Gertsenstein, M., Vintersten, K., Korets-Smith, E., Murphy, J., Stevens, M.E., Haigh, J.J., and Nagy, A. (2007). Developmental and adult phenotyping directly from mutant embryonic stem cells. Proc Natl Acad Sci U S A 104, 4455–4460.

Gerfen, C.R., Paletzki, R., and Heintz, N. (2013). GENSAT BAC cre-recombinase driver lines to study the functional organization of cerebral cortical and basal ganglia circuits. Neuron 80, 1368–1383.

Gray, L.T., Yao, Z., Nguyen, T.N., Kim, T.K., Zeng, H., and Tasic, B. (2017). Layer-specific chromatin accessibility landscapes reveal regulatory networks in adult mouse visual cortex. Elife 6.

Grimm, D., Lee, J.S., Wang, L., Desai, T., Akache, B., Storm, T.A., and Kay, M.A. (2008). In vitro and in vivo gene therapy vector evolution via multispecies interbreeding and retargeting of adeno-associated viruses. J Virol 82, 5887–5911.

Harris, J.A., Hirokawa, K.E., Sorensen, S.A., Gu, H., Mills, M., Ng, L.L., Bohn, P., Mortrud, M., Ouellette, B., Kidney, J., et al. (2014). Anatomical characterization of Cre driver mice for neural circuit mapping and manipulation. Front Neural Circuits 8, 76.

He, M., Tucciarone, J., Lee, S., Nigro, M.J., Kim, Y., Levine, J.M., Kelly, S.M., Krugikov, I., Wu, P., Chen, Y., et al. (2016). Strategies and Tools for Combinatorial Targeting of GABAergic Neurons in Mouse Cerebral Cortex. Neuron 92, 555.

Heinz, S., Benner, C., Spann, N., Bertolino, E., Lin, Y.C., Laslo, P., Cheng, J.X., Murre, C., Singh, H., and Glass, C.K. (2010). Simple combinations of lineage-determining transcription factors prime cis-regulatory elements required for macrophage and B cell identities. Mol Cell 38, 576–589.

Henry, L., and Wickham, H. (2019). purrr: Functional Programming Tools.

Hippenmeyer, S., Vrieseling, E., Sigrist, M., Portmann, T., Laengle, C., Ladle, D.R., and Arber, R. (2005). A developmental switch in the response of DRG neurons to ETS transcription factor signaling. PLoS Biol 3, e159.

Hnasko, T.S., Perez, F.A., Scouras, A.D., Stoll, E.A., Gale, S.D., Luquet, S., Phillips, P.E., Kremer, E.J., and Palmiter, R.D. (2006). Cre recombinase-mediated restoration of nigrostriatal dopamine in dopamine-deficient mice reverses hypophagia and bradykinesia. Proc Natl Acad Sci U S A 103, 8858–8863.

Hodge, R.D., Miller, J.A., Novotny, M., Kalmbach, B.E., Ting, J.T., Bakken, T.E., Aevermann, B.D., Barkan, E.R., Berkowitz-Cerasano, M.L., Cobbs, C., et al. (2020). Transcriptomic evidence that von Economo neurons are regionally specialized extratelencephalic-projecting excitatory neurons. Nat Commun 11, 1172.

Hughes, S.M. (2016). plater: Read, Tidy, and Display Data from Microtiter Plates. Journal of Open Source Software, 106.

Jüttner, J., Szabo, A., Gross-Scherf, B., Morikawa, R.K., Rompani, S.B., Hantz, P., Szikra, T., Esposti, F., Cowan, C.S., Bharioke, A., et al. (2019). Targeting neuronal and glial cell types with synthetic promoter AAVs in mice, non-human primates and humans. Nat Neurosci 22, 1345–1356.

Krijthe, J.H. (2015). {Rtsne}: T-Distributed Stochastic Neighbor Embedding using Barnes-Hut Implementation.

Lamprecht, M.R., Sabatini, D.M., and Carpenter, A.E. (2007). CellProfiler: free, versatile software for automated biological image analysis. Biotechniques 42, 71–75.

Langmead, B., Trapnell, C., Pop, M., and Salzberg, S.L. (2009). Ultrafast and memory-efficient alignment of short DNA sequences to the human genome. Genome Biol 10, R25.

Lawrence, M., Gentleman, R., and Carey, V. (2009). rtracklayer: an R package for interfacing with genome browsers. Bioinformatics 25, 1841–1842.

Lawrence, M., Huber, W., Pagès, H., Aboyoun, P., Carlson, M., Gentleman, R., Morgan, M.T., and Carey, V.J. (2013). Software for computing and annotating genomic ranges. PLoS Comput Biol 9, e1003118.

Levine, J.H., Simonds, E.F., Bendall, S.C., Davis, K.L., Amir, e.-A., Tadmor, M.D., Litvin, O., Fienberg, H.G., Jager, A., Zunder, E.R., et al. (2015). Data-Driven Phenotypic Dissection of AML Reveals Progenitor-like Cells that Correlate with Prognosis. Cell 162, 184–197.

Li, H., Handsaker, B., Wysoker, A., Fennell, T., Ruan, J., Homer, N., Marth, G., Abecasis, G., Durbin, R., and Subgroup, G.P.D.P. (2009). The Sequence Alignment/Map format and SAMtools. Bioinformatics 25, 2078–2079.

Madisen, L., Garner, A.R., Shimaoka, D., Chuong, A.S., Klapoetke, N.C., Li, L., van der Bourg, A., Niino, Y., Egolf, L., Monetti, C., et al. (2015). Transgenic mice for intersectional targeting of neural sensors and effectors with high specificity and performance. Neuron 85, 942–958.

Madisen, L., Zwingman, T.A., Sunkin, S.M., Oh, S.W., Zariwala, H.A., Gu, H., Ng, L.L., Palmiter, R.D., Hawrylycz, M.J., Jones, A.R., et al. (2010). A robust and high-throughput Cre reporting and characterization system for the whole mouse brain. Nat Neurosci 13, 133–140.

Oh, S.W., Harris, J.A., Ng, L., Winslow, B., Cain, N., Mihalas, S., Wang, Q., Lau, C., Kuan, L., Henry, A.M., et al. (2014). A mesoscale connectome of the mouse brain. Nature 508, 207–214.

Paxinos, G., and Franklin, K.B.J. (2013). Paxinos and Franklin’s the mouse brain in stereotaxic coordinates, 4th edn (Amsterdam: Elsevier/AP).

Pliner, H.A., Packer, J.S., McFaline-Figueroa, J.L., Cusanovich, D.A., Daza, R.M., Aghamirzaie, D., Srivatsan, S., Qiu, X., Jackson, D., Minkina, A., et al. (2018). Cicero Predicts cis-Regulatory DNA Interactions from Single-Cell Chromatin Accessibility Data. Mol Cell 71, 858-871.e858.

Ritchie, M.E., Phipson, B., Wu, D., Hu, Y., Law, C.W., Shi, W., and Smyth, G.K. (2015). limma powers differential expression analyses for RNA-sequencing and microarray studies. Nucleic Acids Res 43, e47.

Sayeg, M.K., Weinberg, B.H., Cha, S.S., Goodloe, M., Wong, W.W., and Han, X. (2015). Rationally Designed MicroRNA-Based Genetic Classifiers Target Specific Neurons in the Brain. ACS Synth Biol 4, 788–795.

Stark R, B. G (2011). DiffBind: differential binding analysis of ChIP-Seq peak data.

Taniguchi, H., He, M., Wu, P., Kim, S., Paik, R., Sugino, K., Kvitsiani, D., Kvitsani, D., Fu, Y., Lu, J., et al. (2011). A resource of Cre driver lines for genetic targeting of GABAergic neurons in cerebral cortex. Neuron 71, 995–1013

Taniguchi, H., Lu, J., and Huang, Z.J. (2013). The spatial and temporal origin of chandelier cells in mouse neocortex. Science 339, 70–74.

Tasic, B., Menon, V., Nguyen, T.N., Kim, T.K., Jarsky, T., Yao, Z., Levi, B., Gray, L.T., Sorensen, S.A., Dolbeare, T., et al. (2016). Adult mouse cortical cell taxonomy revealed by single cell transcriptomics. Nat Neurosci 19, 335–346.

Ting, J.T., Daigle, T.L., Chen, Q., and Feng, G. (2014). Acute brain slice methods for adult and aging animals: application of targeted patch clamp analysis and optogenetics. Methods Mol Biol 1183, 221–242.

Ting, J.T., Kalmbach, B., Chong, P., de Frates, R., Keene, C.D., Gwinn, R.P., Cobbs, C., Ko, A.L., Ojemann, J.G., Ellenbogen, R.G., et al. (2018). A robust ex vivo experimental platform for molecular-genetic dissection of adult human neocortical cell types and circuits. Sci Rep 8, 8407.

Vong, L., Ye, C., Yang, Z., Choi, B., Chua, S., Jr., and Lowell, B.B. (2011). Leptin action on GABAergic neurons prevents obesity and reduces inhibitory tone to POMC neurons. Neuron 71, 142–154.

Wertz, A., Trenholm, S., Yonehara, K., Hillier, D., Raics, Z., Leinweber, M., Szalay, G., Ghanem, A., Keller, G., Rózsa, B., et al. (2015). PRESYNAPTIC NETWORKS. Single-cell-initiated monosynaptic tracing reveals layer-specific cortical network modules. Science 349, 70–74.

Wichkham, H. (2016). ggplot2: Elegant Graphics for Data Analysis. Springer-Verlag New York.

Wickham, H. (2007). Reshaping Data with the {reshape} Package. Journal of Statistics Software 21, 1–20.

Wickham, H., François, R., Henry, L., and Müller, K. (2018). dplyr: A Grammar of Data Manipulation.

Wilke, C.O. (2018). cowplot: Streamlined Plot Theme and Plot Annotations for “ggplot2.”.

Zachary, F., Scott, C., and Niklaus, G. (2018). Taxa: An R package implementing data standards and methods for taxonomic data. F1000Research 7.

Zeng, H., Horie, K., Madisen, L., Pavlova, M.N., Gragerova, G., Rohde, A.D., Schimpf, B.A., Liang, Y., Ojala, E., Kramer, F., et al. (2008). An inducible and reversible mouse genetic rescue system. PLoS Genet 4, e1000069.

Zhu, F., Gamboa, M., Farruggio, A.P., Hippenmeyer, S., Tasic, B., Schüle, B., Chen-Tsai, Y., and Calos, M.P. (2014). DICE, an efficient system for iterative genomic editing in human pluripotent stem cells. Nucleic Acids Res 42, e34.

